# Novel prokaryotic sensing and regulatory system employing previously unknown nucleic acids-based receptors

**DOI:** 10.1101/2021.09.11.459467

**Authors:** Victor Tetz, George Tetz

## Abstract

The present study describes a previously unknown universal signaling and regulatory system, which we named TRB receptor system. This system is responsible for sensing, remembering, and regulating cell responses to various chemical, physical or biological stimuli. It controls cell survival, variability, reproduction, adaptation, genome changes, and gene transfer. Importantly, the TRB-receptor system is responsible for the formation and maintenance of cell memory, as well the ability to “forget” preceding events. The system is composed of DNA- and RNA-based receptors located outside the membrane named “TezRs”, as well as reverse transcriptases and integrases. The sensory and regulatory functions of TezRs enable the TRB-receptor system to control all major aspects of bacterial behavior, such as growth, biofilm formation and dispersal, utilization of nutrients including xenobiotics, virulence, chemo- and magnetoreception, response to external factors (e.g., temperature, UV, light and gas content), mutation events, phage-host interaction and recombination activity. Additionally, it supervises the function of other receptor-mediated signaling pathways. Transcriptome analysis revealed that the loss of different TezRs instigates significant alterations in gene expression.

**HIGHLIGHTS:** The TRB-receptor system regulates bacterial sensing and response to various stimuli.

The TRB-receptor system is responsible for maintenance and loss of cell memory.

The TRB-receptor system comprises DNA- and RNA-based “TezRs” receptors.

The TRB-receptor system relies on reverse transcriptases and recombinases.

The TRB-receptor system oversees other receptor-mediated signaling pathways.

TezRs are implicated in cell mutation and recombination events.

## INTRODUCTION

To ensure survival, bacteria need to adapt to a constantly changing environment. Yet, the details of sensory and biophysical processes involved in reception have remained elusive (1–3). At present, these adaptations are known to be mediated by a variety of predominantly transmembrane receptors consist of a protein structure, which control different key aspects of the interaction with the environment, cell-to-cell signaling, and multicellular behavior. Chemoreceptors represent the most well studied type of bacterial receptors (3–7). They recognize various signals, primarily growth substrates or toxins (8,9). Chemoreception is tightly linked to chemotaxis and provides bacteria with the capacity to approach or escape different compounds, thus favoring the movement toward optimal ecological niches (10). However, many aspects of chemoreception remain unclear, including details of the mechanisms underlying high sensitivity, sensing of multiple stimuli, and recognition of previously unknown nutrients or xenobiotics (11–13).

Bacterial receptive function and interaction with the environment is coupled to bacterial memory, another poorly characterized phenomenon (14–20).

Cell memory is viewed as a part of history-dependent behavior and is intended as a means for the efficient adaptation to recurring stimuli. It can be encoded by membrane potential, which is also associated with transmembrane receptors in bacteria (21).

Sensing of physical factors by bacteria remains even more elusive. For example, the mechanism of magnetoreception, whereby microorganisms sense the geomagnetic field, has been well described only in magnetotactic bacteria (22). These prokaryotes sense magnetic fields due to the biomineralization of nano-sized magnets, termed magnetosomes, within cells (23,24). However, existing studies have not explained why bacteria lacking these elements could still sense the magnetic field (25,26). Recent data suggest that intracellular DNA can be affected by magnetic fields and is able to interact with them, but the nature of such interactions remains enigmatic (27–29).

The mechanism and regulation of bacterial temperature sensing is also characterized by numerous unknowns. Different studies have pointed to Tar/Tsr receptors as responsible for controlling and regulating the temperature response, but the detailed mechanisms of their reception remain elusive (30–33). Some authors also highlight the sensing of the temperature that is associated with blue-light sensing through the BlsA Sensor (34,35).

Therefore, the question of how known receptors sense a diverse array of chemical, biological, and physical factors remains insufficiently explored. It has been suggested that certain protein receptors could be organized into sensory arrays, whereby cooperative interactions between receptors enable the sensing of a diverse range of stimuli (7,36–39). Still, even such clusters could not account for the totality of different stimuli sensed by bacteria. Even in the case of known receptive systems it remains to be determined how bacteria sense the whole plethora of available environmental factors including previously unknown exogenous stimuli, how remote sensing operates, what is the common sensor part of most receptors, and how signal transduction is mediated. Therefore, a better understanding of receptors and receptor systems could expand our knowledge of the regulation of bacterial physiology, virulence, and adaptation.

In this work, we report for the first time the identification of novel bacterial elements constituted by nucleic acid molecules (located outside the cell membrane and presumably also inside the cell), which can sense and amplify the signals from different chemical, physical, and biological stimuli into an integrated output (40). Because they possess the features of receptors and regulators, we named these elements Teazeled Receptors (TezRs). Here, we confirmed their receptor and regulatory activities, and also revealed their participation in cell memory formation, maintenance, and loss. Finally, we demonstrated that TezRs were part of a previously unknown receptor system, which we named TezR-based receptor system (TRB-receptor system).

## RESULTS

### Nucleases remove cell surface-bound nucleic acids

First, we confirmed the destruction of cell-surface bound DNA and RNA by studying the changes in fluorescence of washed planktonic *B. pumilus* VT1200 following their treatment with 10 µg/mL DNase I and RNase A for 15 min or a combination of the two. SYTOX Green-stained *B. pumilus* displayed clear green fluorescence, confirming the presence of cell surface-bound nucleic acids, which were not removed upon washing of culture medium or matrix (Fig. 1A, B).

**Figure 1.**
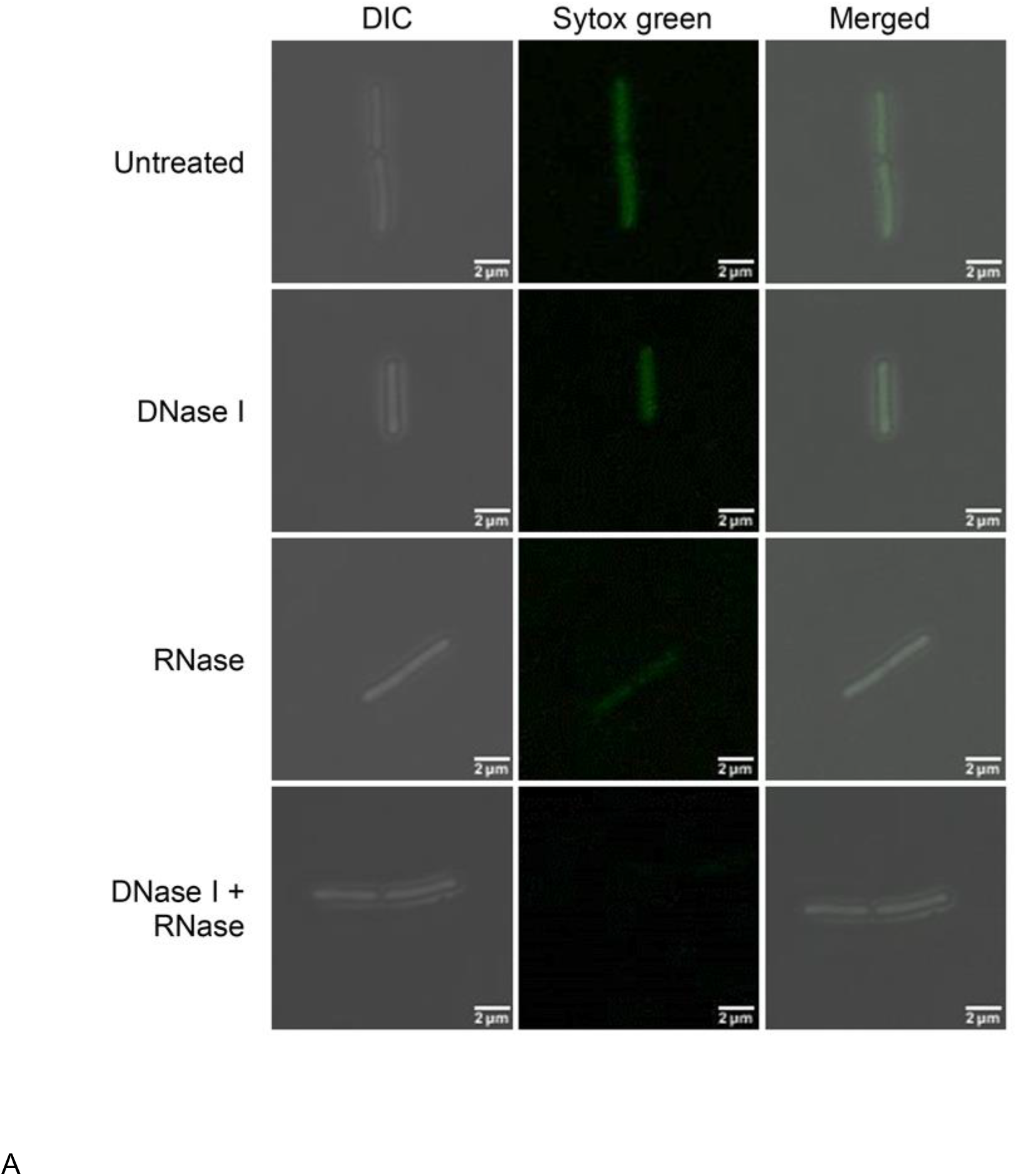
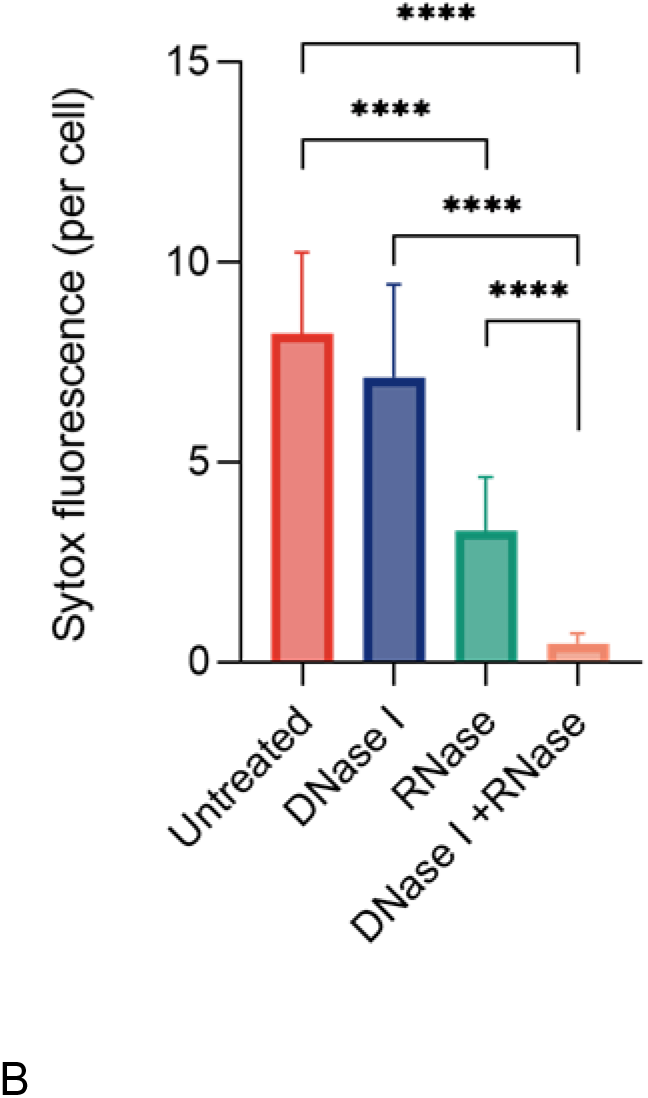
Removal of *B. pumilus* cell surface-bound DNA and RNA molecules with nucleases. Green fluorescence denotes cell surface-bound DNA and RNA of *B. pumilus* stained with the membrane impermeable SYTOX Green dye. (A) DIC (left), SYTOX Green (center), and merged (right) images of untreated and DNase/RNase-treated *B. pumilus* at 100× magnification. Scale bars represent 2 μm. (B) Quantification of SYTOX Green signal intensity per cell (n = 10; mean ± SD). ****p < 0.0001, two-tailed unpaired *t*-tests.

Bacteria treated with either DNase or RNase alone exhibited a decrease, but not the total disappearance of fluorescence compared to untreated cells (p < 0.0001). Instead, bacteria treated with a combination of DNase and RNase revealed the total disappearance of surface fluorescence compared to single-nuclease treatment (p < 0.0001).

As it was outside the scope of our study to evaluate which part of the cell surface-bound DNA or RNA exerted receptive functions, in the following experiments we applied the same nuclease treatment regimen that resulted in total removal of all cell surface-bound nucleic acids as observed here.

Next, we verified that the RNase A used in this study was not internalized by the bacteria. To examine the ability of RNase A to penetrate the bacterial cell wall we linked the enzyme with a fluorophore. To score the penetration capability of RNase in *B. pumilus* we incubated *B. pumilus* on agar media supplemented with fluorophore-linked RNase or cultivated pre-treated *B. pumilus* with the same RNase. However, in both experiments no signs of RNase internalization were observed. (Supplementary Fig. 1).

### TezR destruction has a global impact on gene expression

To gain insight into the consequences of TezRs loss on bacterial gene expression, RNA-seq analyses of *S. aureus* gene expression profile were examined following the removal of primary TezRs. Principal-component analysis (PCA) showed that *S. aureus* due to the loss of primary TezRs clustered separately from the control group of *S. aureus* where TezRs was intact. The largest difference in PCA was observed for *S. aureus* TezR_R1^d^ (Fig. 2A). These differences in gene expression datasets are also clearly evident in the hierarchical clustering and heatmaps of Euclidean distance. Strikingly, the largest pairwise Euclidean distance was observed between the control *S. aureus* and TezR_R1^d^ (Fig. 2B).

**Figure 2.**
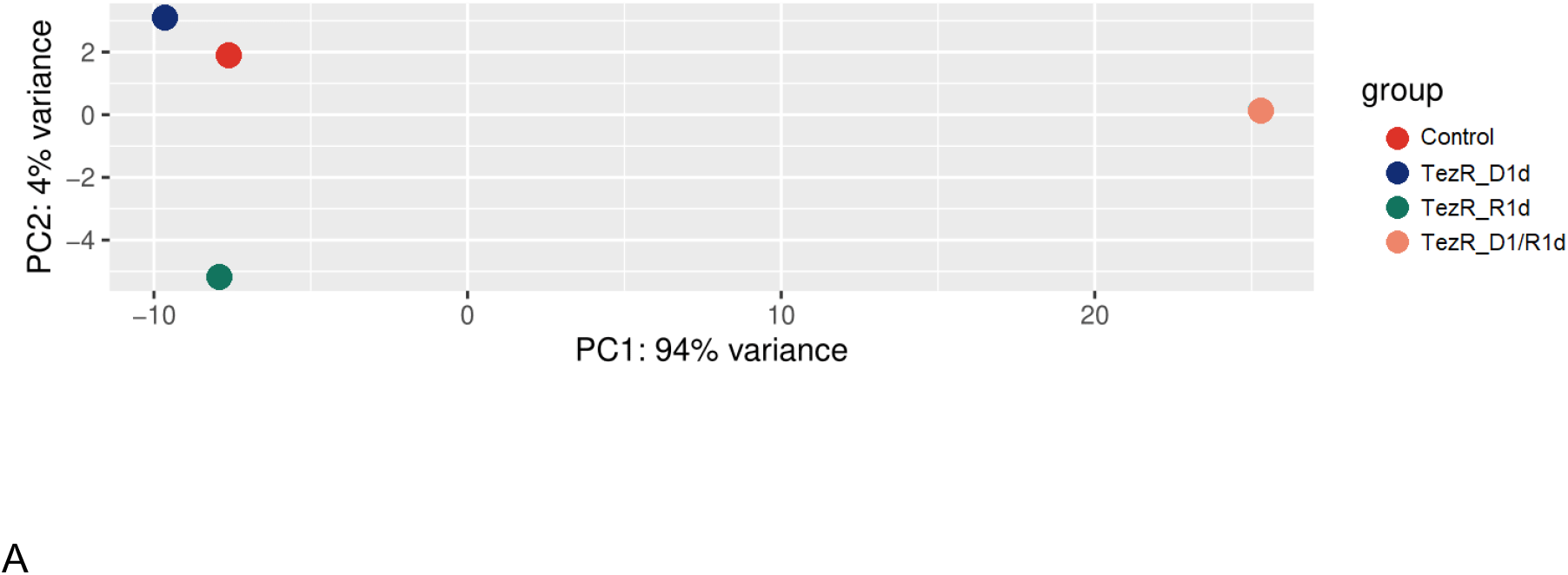
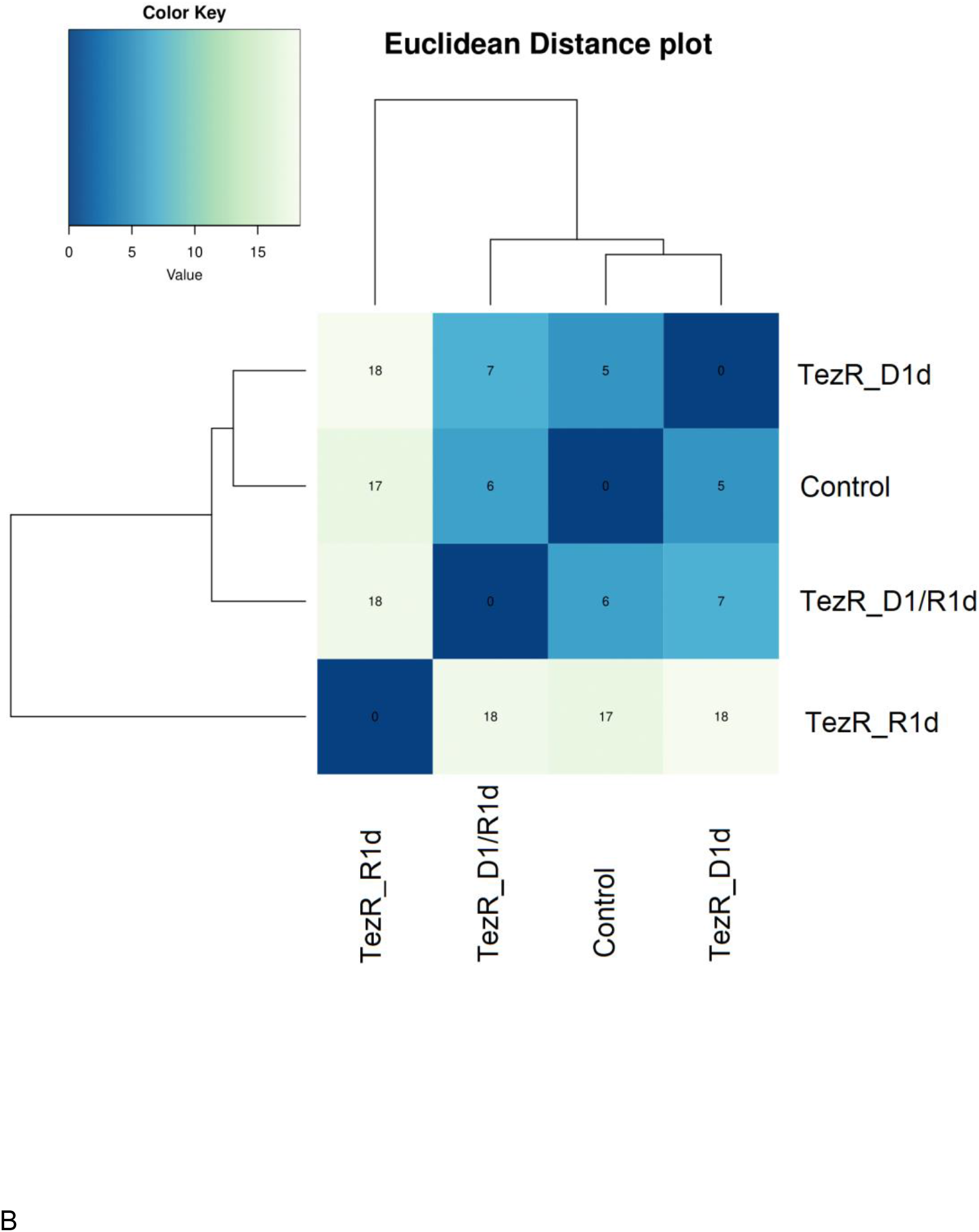
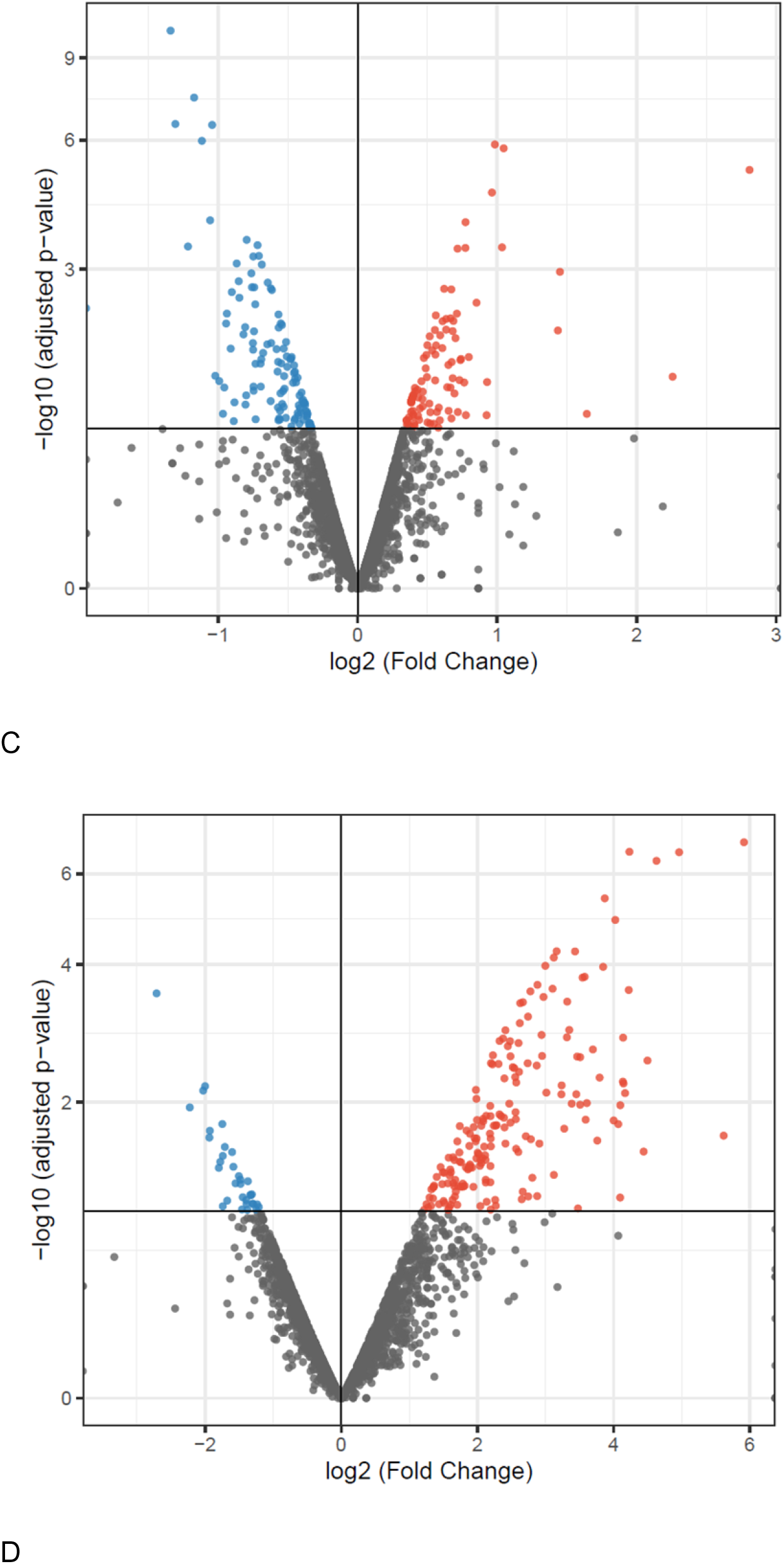
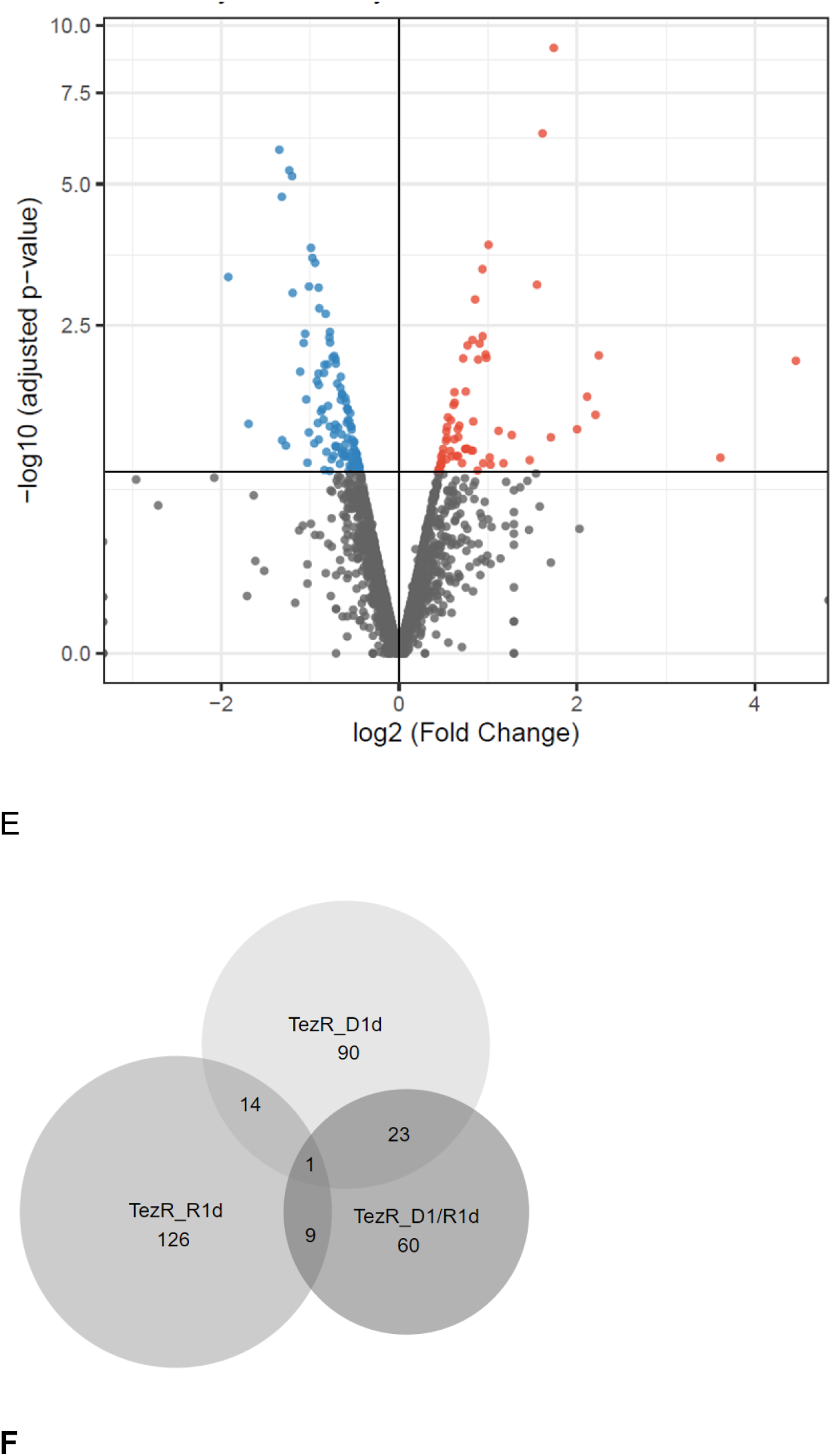
Transcriptome analysis of *S. aureus* following the removal of primary TezRs (A) PCA plot (B) Heatmap of Euclidean distance (C to E) Volcano plots highlighting the genes that are differentially expressed (log_2_ fold> 0.5 change plotted against the –log_10_ P-value). The results demonstrate the altered expression levels of the genes following primary TezRs loss (F) the overlap and unique DEPs in each group using Venn diagram

Next, we compared the results from each probe and analyzed the genes whose expressions were significantly altered (upregulated or downregulated) following the removal of different TezRs (Fig. 2C to D, Supplementary table S1). We identified 128, 150, and 93 differentially expressed proteins (DEPs) in *S.aureus* when compared to TezR_D1^d^/control, TezR_R1^d^/control, and TezR_D1/R1^d^/control, respectively (|log_2_-fold change| > 0.5 and p-value < 0.05). Among the DEPs, 55 proteins were upregulated, and 73 proteins were downregulated in *S.aureus* TezR_D1^d^ compared to those in the TezR_D1^d^/control (Fig. 2C). Among the DEPs in *S.aureus* TezR_R1^d^, 137 upregulated and 13 downregulated proteins are found compared to those in the TezR_R1^d^/control (Fig. 2D). Additionally, 62 upregulated proteins and 31 downregulated proteins are detected in TezR_D1/R1^d^/control compared to those in the TezR_D1/R1^d^/control. A minute overlap in differentially expressed transcripts were detected in bacteria after the removal of different TezRs. This non-redundancy signifies the individual regulatory roles of TezRs. These data evidently highlight the complex responses triggered by the loss of both primary DNA- and RNA-based TezRs, which cannot be justified by summing up the effects of individual TezRs losses (Fig. 2E). The only gene expression which significantly altered due to the loss of any of the primary TezRs was SA0532 encoding a *Staphylococcus*-specific hypothetical protein (41). Interestingly, following the loss of DNA-based TezRs alone or in combination with RNA-based TezRs, upregulation of proteins associated with type VII secretion system was observed (42,43).

### TezRs affect microbial growth

Stationary phase *S. aureus* VT209 and *E. coli* ATCC 25922 were left untreated or pretreated with nucleases to remove primary TezRs, after which they were diluted in fresh medium and allowed to grow. OD600 and CFU were measured hourly during the first 6 h of incubation. Growth curves are presented as OD600 values (Fig. 2A, B) or bacterial counts (Fig. 2C, D) as a function of time.

Removal of primary TezRs retarded bacterial growth in both *S. aureus* and *E. coli* compared with untreated bacteria as measured by OD600 (p < 0.001 and p < 0.05, respectively) and CFU. While the lag phase was 3-h longer for treated *S. aureus*, it was similar between untreated and treated *E. coli*; although the latter exhibited retarded growth by the end of the observation period. At that point, CFU/mL of *S. aureus* TezR_D1^d^ and *E. coli* TezR_D1^d^ were lower by 2.6 log10 (p < 0.05) and 2.1 log10 (p < 0.001) compared with control bacteria.

Loss of TezR_R1 in *S. aureus* inhibited bacterial growth, as indicated by OD600 values (p < 0.05), but it did not affect bacterial counts. Such a discrepancy points to dysregulation of *S. aureus* TezR_R1^d^ and can be explained by reduced production of extracellular matrix. A similar effect on growth was observed in *E. coli* following the removal of TezR_R1 (OD600, p < 0.05); however, unlike in *S. aureus*, it coincided with reduced CFU (p < 0.05).

Loss of both primary TezRs in *S. aureus* and *E. coli* extended the lag phase by 3 h; however, this was followed by very rapid growth from 3 to 6 h. Thus, by the end of the observation period, OD600 for *S. aureus* TezR_D1/R1^d^ was even higher than for control *S. aureus*; while OD600 for *E. coli* TezR_D1/R1^d^ was only marginally lower than for control *E. coli*. Surprisingly, bacterial counts of *S. aureus* TezR_D1/R1^d^ and *E. coli* TezR_D1/R1^d^ were lower throughout the observation period, amounting to 2.4 log10 CFU/mL and 1.2 log10 CFU/mL fewer counts compared with control bacteria after 6 h (p < 0.05). Cell size was also reduced at this time point (Supplementary Table 1).

The discrepancy between elevated OD600 levels along with delayed bacterial growth and a reduced cell size can be explained by the production of more extracellular matrix. Given similar OD600 values at the end of the observation period between control bacteria and those lacking TezR_D1/R1, we named the latter “drunk cells”.

Based on these data we conclude that primary TezRs play a critical regulatory role in bacterial growth by affecting multiple biosynthetic pathways.

### Biofilm growth and cell size are regulated by TezRs

We next investigated how TezRs affected biofilm morphology of *B. pumilus* VT1200 grown on agar plates. To analyze the role of primary TezRs, *B. pumilus* were pretreated with nucleases and then inoculated and grown on regular agar medium. To study the role of secondary TezRs, growth of *B. pumilus* was evaluated on medium supplemented with different nucleases. We also established that RNase A used in this study was not internalized by the bacteria under these experimental conditions (Supplementary Fig. 1).

Biofilms of control *B. pumilus* had a circular shape (Fig. 4A) with smooth margins; whereas those formed by *B. pumilus* TezR_D1^d^ (Fig. 4B) and *B. pumilus* TezR_R1^d^ (Fig. 4C) develop blebbing, and those of *B. pumilus* TezR_D1/R1^d^ exhibited filamentous (filiform) margins (Fig. 4D). *B. pumilus* TezR_D2^d^ biofilms were characterized by increased swarming motility and formation of significantly larger colonies (p < 0.001) with distinct phenotype and dendritic patterns (Fig. 4E); whereas *B. pumilus* TezR_R2^d^ biofilms had the same size as control *B. pumilus*, but irregular margins and wrinkled surface (Fig. 4F, Supplementary Table 2).

**Figure 3.**
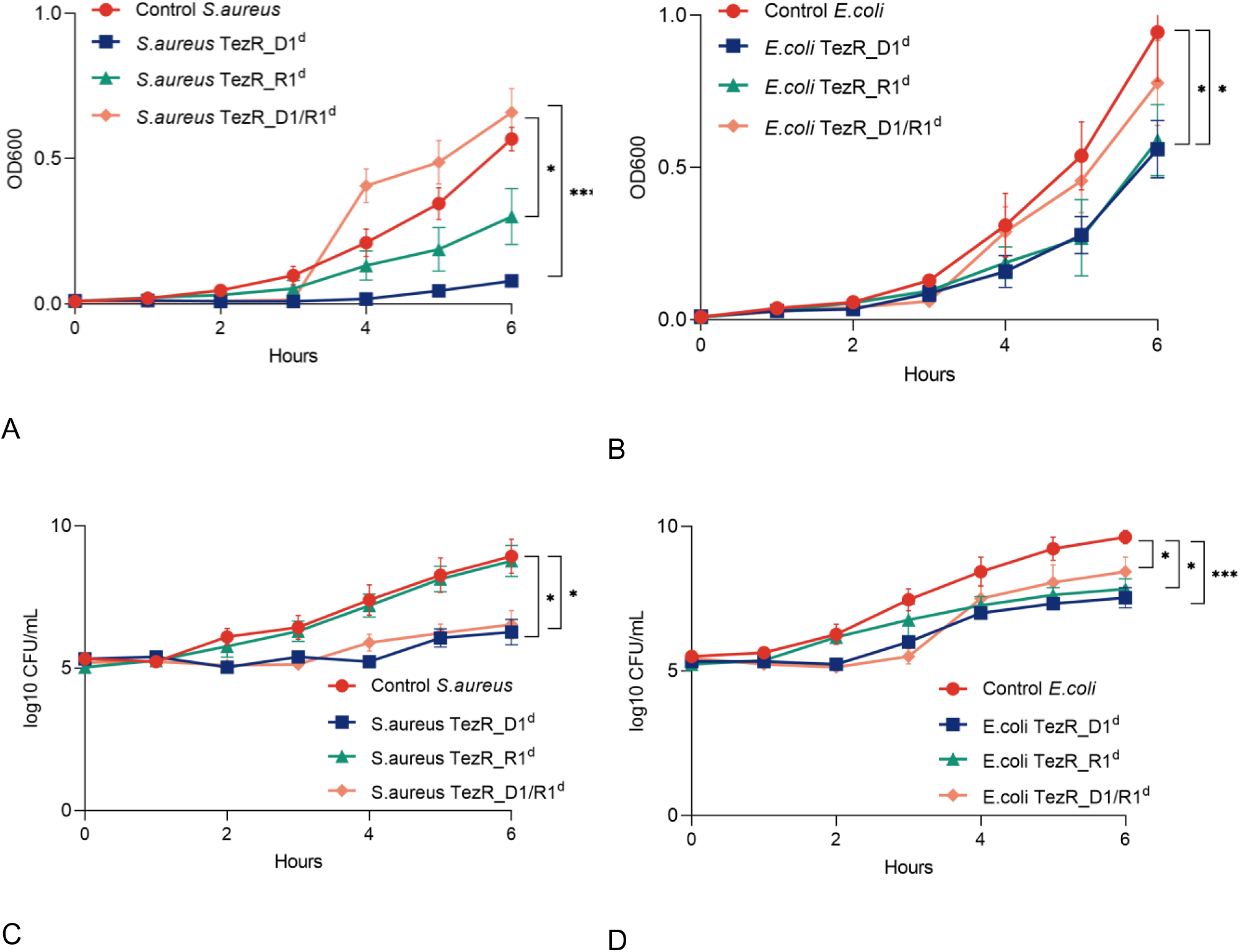
Role of TezRs in the regulation of bacterial growth. Growth comparison of control bacteria and bacteria lacking TezR_D1 (*S. aureus* TezR_D1^d^, *E. coli* TezR_D1^d^), TezR_R1 (*S. aureus* TezR_R1^d^, *E. coli* TezR_R1^d^) or TezR_D1 and TezR_R1 (*S. aureus* TezR_D1/R1^d^, *E. coli* TezR_D1/R1^d^). (A, B) Bacterial growth measured as OD600 over time in (A) *S. aureus* and (B) *E. coli*. (C, D) Bacterial growth measured as bacterial counts (log10 CFU/mL) in (C) *S. aureus* and (D) *E. coli*. Values representing the mean ± SD were normalized to the initial OD600 value. *p < 0.05, ***p < 0.001.

**Figure 4.**
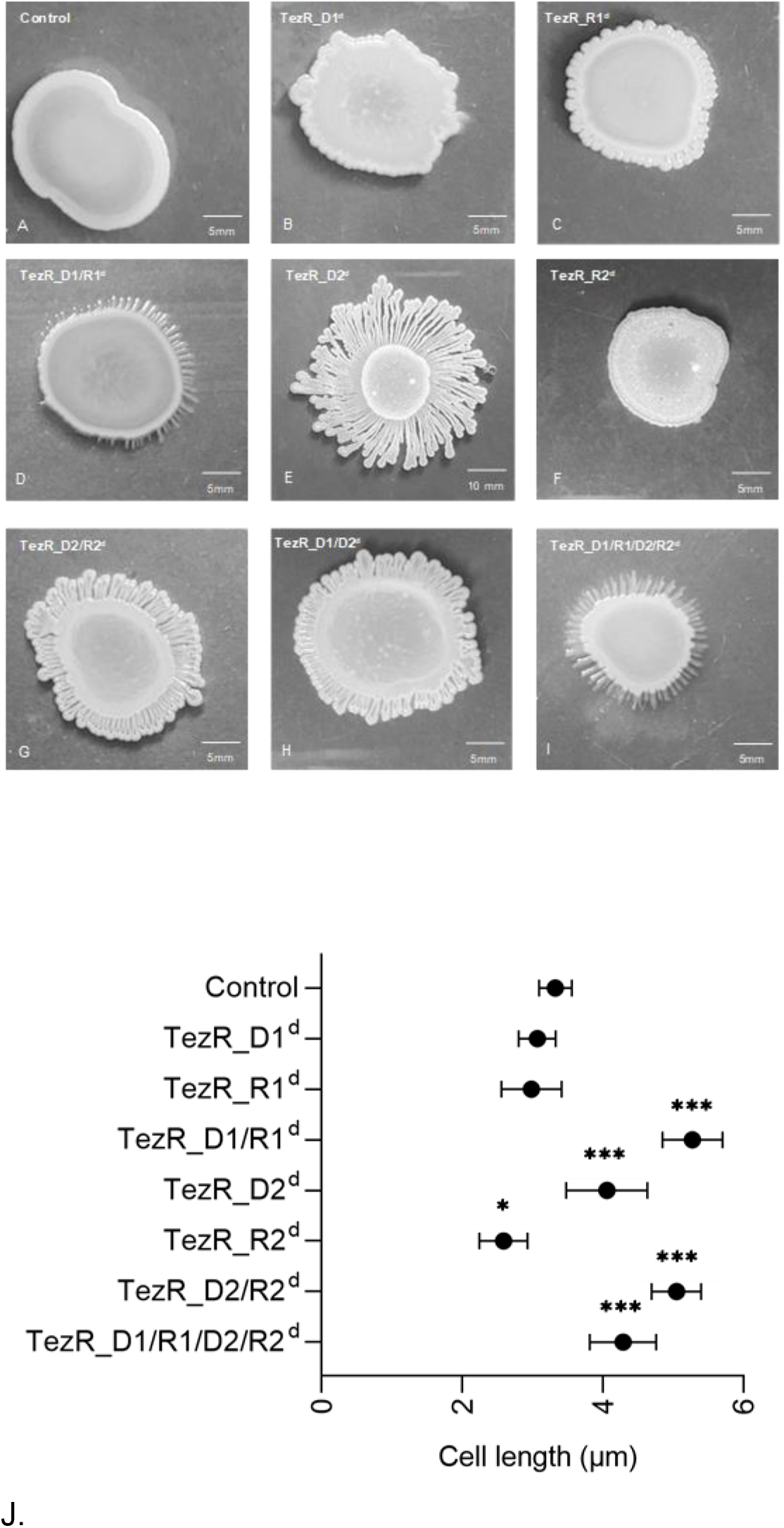
TezRs regulate biofilm morphology and cell size. Morphology of nuclease-treated or untreated 72-h-old biofilms. (A) Control *B. pumilus*. (B) *B. pumilus* TezR_D1^d^. (C) *B. pumilus* TezR_R1^d^. (D) *B. pumilus* TezR_D1/R1^d^. (E) *B. pumilus* TezR_D2^d^. (F) *B. pumilus* TezR_R2^d^. (G) *B. pumilus* TezR_D2/R2^d^. (H) *B. pumilus* TezR_D1/D2^d^. (I) *B. pumilus* TezR_D1/R1/D2/R2^d^. Scale bars indicate 5 or 10 mm. Representative images of three independent experiments are shown. (J). Cell length of bacteria grown on solid medium (µm). *p < 0.05, ***p < 0.001. Data represent the mean ± SD from three independent experiments.

Interestingly, the combined removal of other TezRs along with loss of TezR_D2 led to a striking difference compared to the large biofilms formed by *B. pumilus* TezR_D2^d^. The biofilms of both *B. pumilus* TezR_D2/R2^d^ and TezR_D1/D2^d^ were characterized by a structurally complex, densely branched morphology, but the dendrites were not so profound and the biofilm was not so spread out as in the case of *B. pumilus* TezR_D2^d^. The morphology of biofilms formed by bacteria devoid of both primary and secondary TezRs such as *B. pumilus* TezR_D1/R1/D2/R2^d^ was very similar to that of *B. pumilus* TezR_D1/R1^d^, with filamentous (filiform) margins but similar size as control *B. pumilus*.

In these experiments, nucleases added to the solid nutrient medium with the aim of removing secondary TezRs could potentially affect also cell surface-bound primary TezRs. However, a comparison of the morphology of biofilms formed by *B. pumilus* TezR_D1^d^ with those of *B. pumilus* TezR_D2^d^ and *B. pumilus* TezR_D1/D2^d^ (Fig. 4B, E, H) revealed clear differences, meaning that nucleases added to the agar did not alter primary TezRs, at least not in the same way as direct nuclease treatment did.

Moreover, the different size of biofilms formed by *B. pumilus* TezR_D2^d^ vs. *B. pumilus* TezR_D1/D2^d^ excludes the possibility that the increased colony size of the former resulted from greater swarming motility due to loss of extracellular DNA and decreased extracellular polysaccharide viscosity, because extracellular DNA was eliminated also in the latter (44). Collectively, these data allow us to conclude that different TezRs play an individual regulatory role in biofilm morphology.

Next, we found that loss of TezRs had divergent effects on bacterial size. The combined removal of primary TezRs, or secondary TezR_D2 alone or in combination with other TezRs, resulted in significantly increased cell sizes (p < 0.001). In comparison, individual loss of secondary TezR_R2s decreased the size of *B. pumilus* cells (p < 0.05). Further experiments could not confirm an association between cell size alteration and sporulation triggered by TezRs removal. Possibly, the observed greater mean cell length could result from incomplete cell division and elongation triggered by TezRs destruction (45).

### TezRs modulate sporulation

Given the significant alterations of biofilm morphology and transcriptome following TezRs loss, we sought evidence for their biological relevance in sporulation. We found that loss of TezR_D1, TezR_R1, and particularly TezR_R2 activated sporulation of *B. pumilus* VT1200 (all p < 0.001) (Fig. 5, Supplementary Table 3). In contrast, destruction of TezR_D2 completely repressed sporulation (p = 0.007) (Supplementary Table 3).

**Figure 5.**
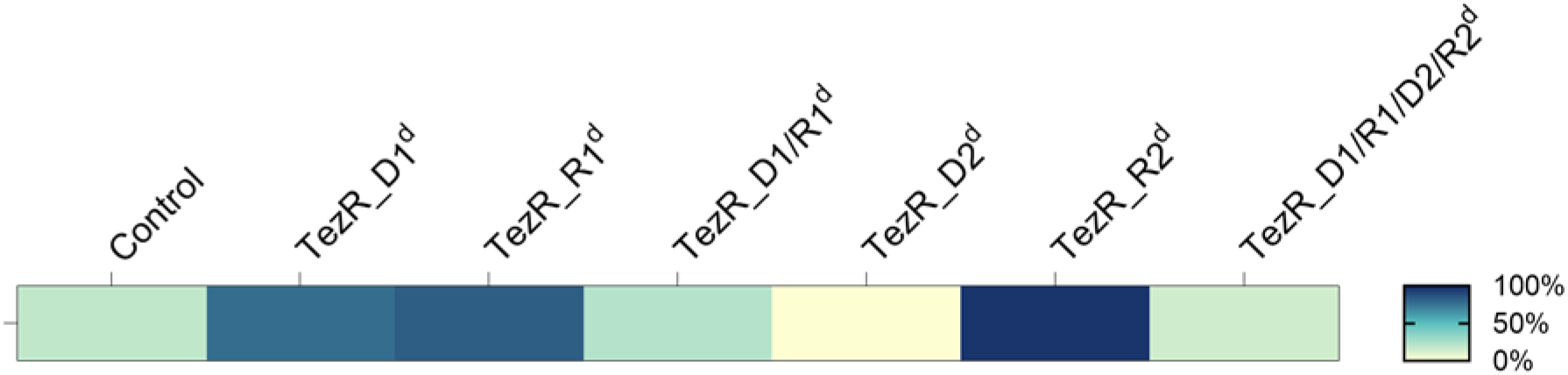
TezRs regulate sporulation. Heat map of sporulation intensity in cells with altered TezRs under normal conditions. Each cell indicates control *B. pumilus* or *B. pumilus* lacking TezRs. Color-coding indicates the ratio of spores to the total number of cells: white (0% sporulation), dark blue (100% sporulation).

Notably, sporulation was not affected in “drunk cells” lacking TezR_D1/R1, but was increased if either TezR_D1 or TezR_R1 were removed. This finding highlights the complex web of pathways dictating the responses of “drunk cells”, which do not simply reflect the additive effect of removing individual primary TezRs. Moreover, the result points to the various roles of TezRs in regulating bacterial sporulation.

### Role of TezRs in the regulation of stress responses

We next tested whether TezRs regulated also stress responses. The general stress response of control *B. pumilus* VT1200 manifested as increased sporulation (Fig. 6). Removal of TezR_R1 or TezR_R2 alone, or in combination with any other TezRs, upregulated the stress response and stimulated sporulation. Interestingly though, loss of TezR_D1 or TezR_D2 had the opposite effect (p < 0.001) (Supplementary Table 4). Hence, loss of TezR_D2 inhibited sporulation under both normal and stress conditions, confirming its implication in regulating the cell stress response.

**Figure 6.**
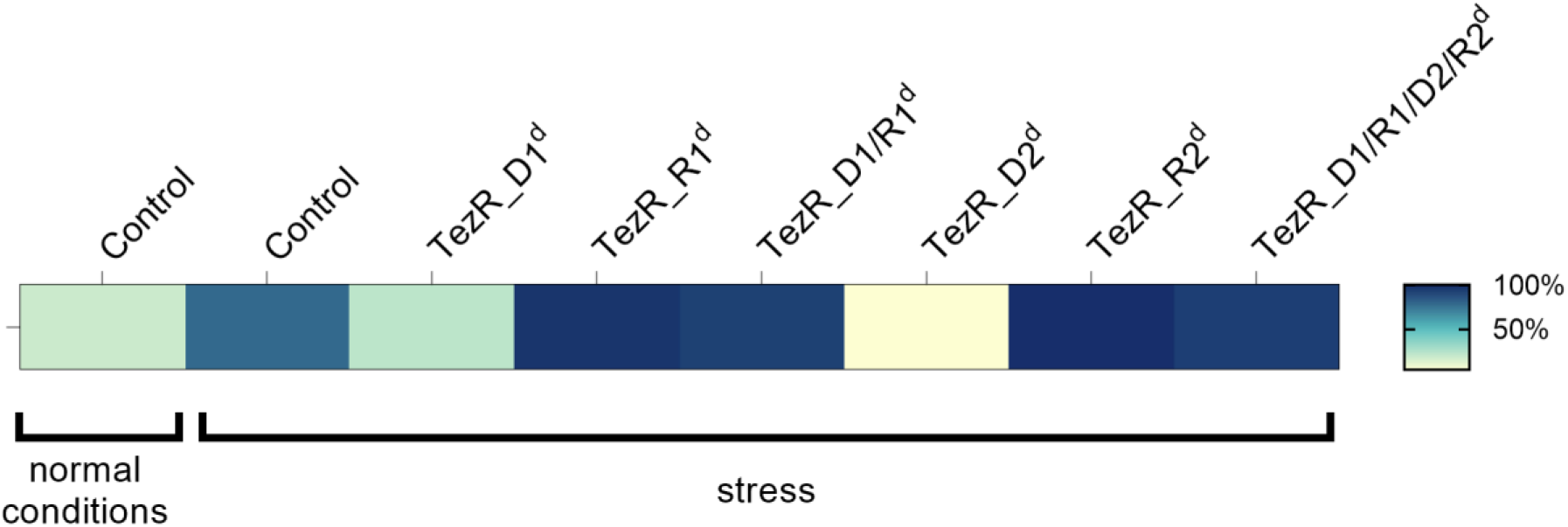
TezRs regulate sporulation under stress. Heat map of sporulation intensity in *B. pumilus* with altered TezRs under stress conditions. Each cell indicates control *B. pumilus* or *B. pumilus* lacking TezRs under stress conditions. Color-coding indicates the ratio of spores to the total number of cells: white (0% sporulation), dark blue (100% sporulation).

### TezRs removal results in increased temperature tolerance

Assessment of whether TezRs regulated bacterial thermotolerance revealed that control *S. aureus* VT209 exhibited maximum tolerance at up to 50 °C, whereas *S. aureus* lacking primary TezRs could survive at even higher temperatures. Specifically, *S. aureus* TezR_D1^d^ survived at up to 65 °C, *S. aureus* TezR_R1^d^ at up to 70 °C, and *S. aureus* TezR_D1/R1^d^ at up to 60 °C (Fig. 7A).

**Figure 7.**
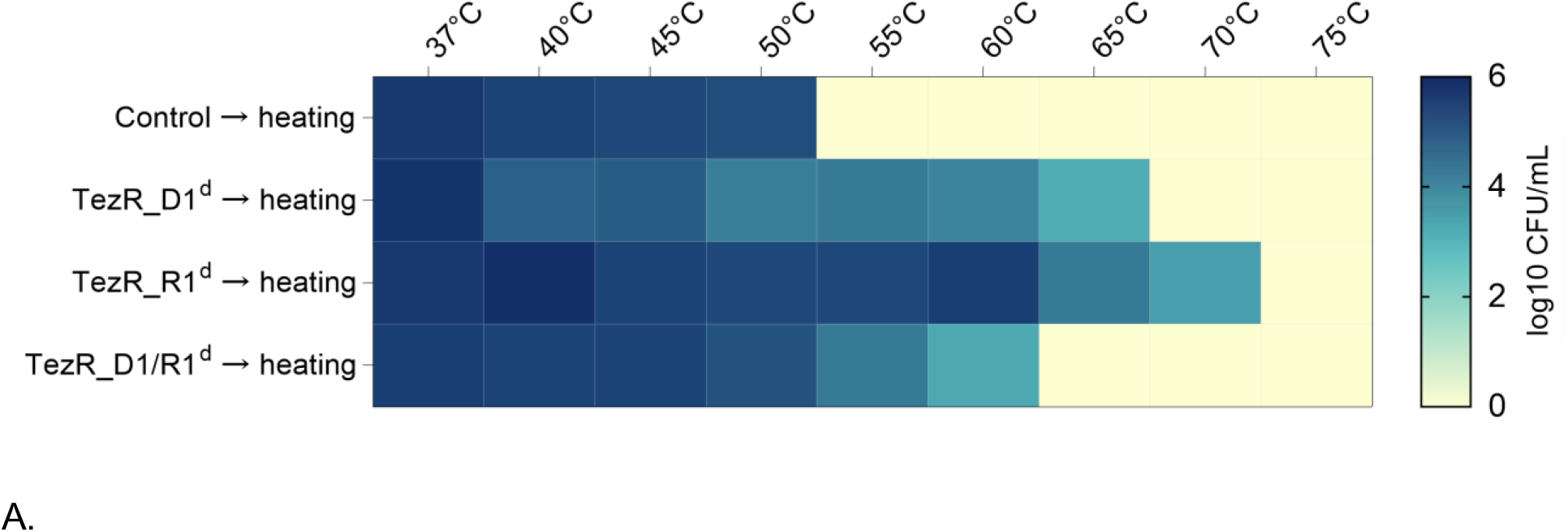
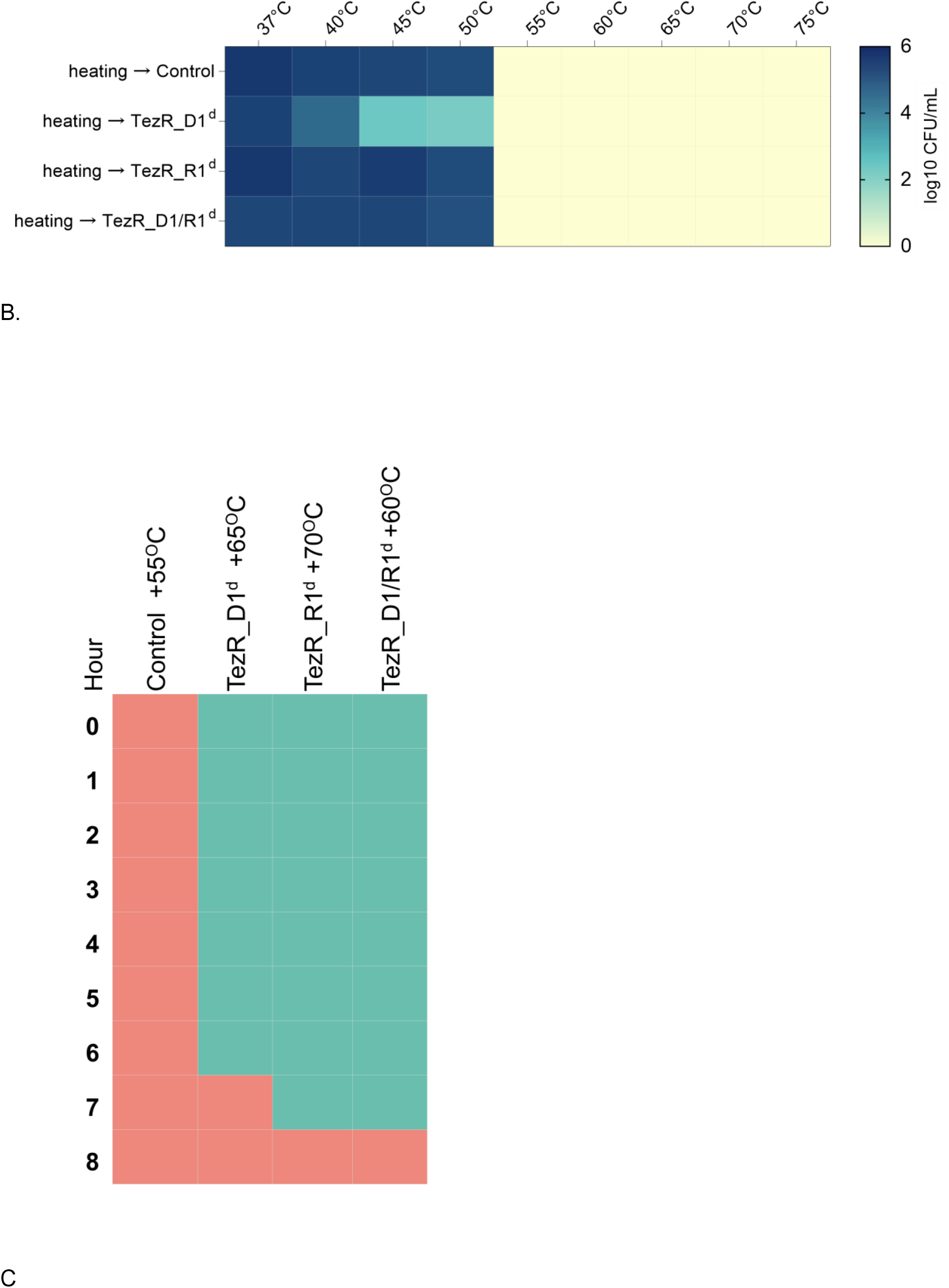
Role of TezRs in survival at the elevated temperature. (A) Heat map summarizing the effect of primary TezRs removal on survival of a *S. aureus* culture heated for 10 min at different temperatures. The color intensity represents the average log10 CFU/mL, from white (minimal) to blue (maximum). Values represent the average of three independed experiments. (B) Heat map summarizing the effect of primary TezRs removal on survival after heating of a *S. aureus* culture at different temperatures for 10 min. The color intensity represents the average log10 CFU/mL, from white (minimal) to blue (maximum). Values represent the average of three independed experiments. (C) Heat map representing the time required for the enhanced temperature tolerance of *S. aureus* to disappear in control, TezR_D1^d^ (65 °C), TezR_R1^d^ (70 °C), and TezR_D1/R1^d^ (60 °C) cells. Green squares denote bacterial growth following heating and indicate enhanced temperature survival. Red squares denote lack of bacterial growth following heating and indicate no change in temperature tolerance. Values represent the average of three independed experiments.

We sought to discern whether the observed enhanced temperature survival was attributable to transcriptome-level responses triggered by TezRs removal, or to the direct role of TezRs in sensing and regulation of temperature changes. To this end, we incubated control *S. aureus* at different temperatures and removed primary TezRs right after heating to trigger transcriptionally-induced alterations. Loss of primary TezRs after the heating step did not improve temperature tolerance (Fig. 7B). This result demonstrated that the response of bacteria to higher temperatures was regulated by primary TezRs and depended on their presence at the time of heating, rather than being induced by their loss.

Next, we evaluated how much time was required for bacteria, which became resistant to heating after primary TezRs removal, to recover normal temperature sensing. This information could be used as a surrogate marker of the time required for restoration of functionally active cell surface-bound TezRs. *S. aureus* TezR_D1^d^, TezR_R1^d^, and TezR_D1/R1^d^ were inoculated in culture broth and grown at the maximum temperature tolerated by bacteria following each specific TezR destruction (65, 70, and 60°C, respectively) (Fig. 7C). Control *S. aureus* were processed in the same way and heated at 55 °C as their next-to-lowest non-tolerable temperature. Each hour after heating, bacteria were inoculated in fresh LB broth to assess the presence or absence of growth after 24 h at 37 °C. Growth meant that bacteria still possessed enhanced temperature survival and the corresponding time indicated no restoration of functionally active primary TezRs. In turn, absence of growth could mean that functionally active primary TezRs were restored and bacteria could normally sense and respond to the higher temperature. After TezRs removal, it took from 7 to 8 h for *S. aureus* to restore functionally active primary TezRs and normal temperature tolerance (Fig. 7C). Taken together, these data demonstrate that TezRs participate in temperature sensing and the regulation of the corresponding response.

### TezRs regulate UV resistance

To determine whether TezRs participated in UV resistance, we exposed cells to UV light. Loss of TezR_D1 and TezR_D1/R1 had no statistically significant effect on the survival of *S. aureus* following UV irradiation compared to control bacteria (Fig. 8). Notably, loss of TezR_R1 protected bacteria from UV-induced death, and resulted in 2.4 log10 CFU/mL higher viable counts compared to control *S. aureus* following UV irradiation (p = 0.002). These data suggest that TezRs participate in sensing and response to UV irradiation.

**Figure 8.**
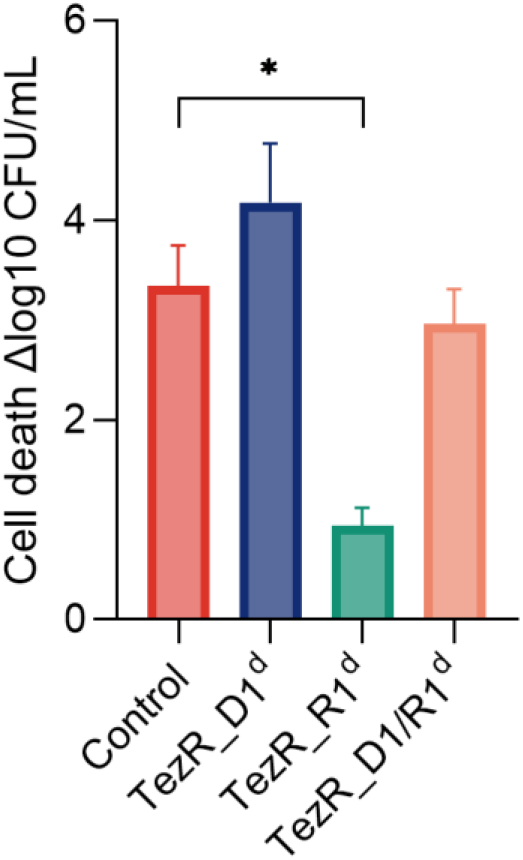
Role of primary TezRs in resistance of *S. aureus* to UV exposure. Comparison of live bacteria measured as bacterial counts (log10 CFU/mL) before and after UV exposure. Data represent the mean ± SD of three independent experiments. p < 0.05 was considered significant.

### Magnetoreception relies on TezRs

The magnetoreceptive function of TezRs was assessed by morphological changes at a macroscopic scale in agar-grown *B. pumilus* VT1200 biofilms following inhibition of the geomagnetic field (Fig. 9A, B).

**Figure 9.**
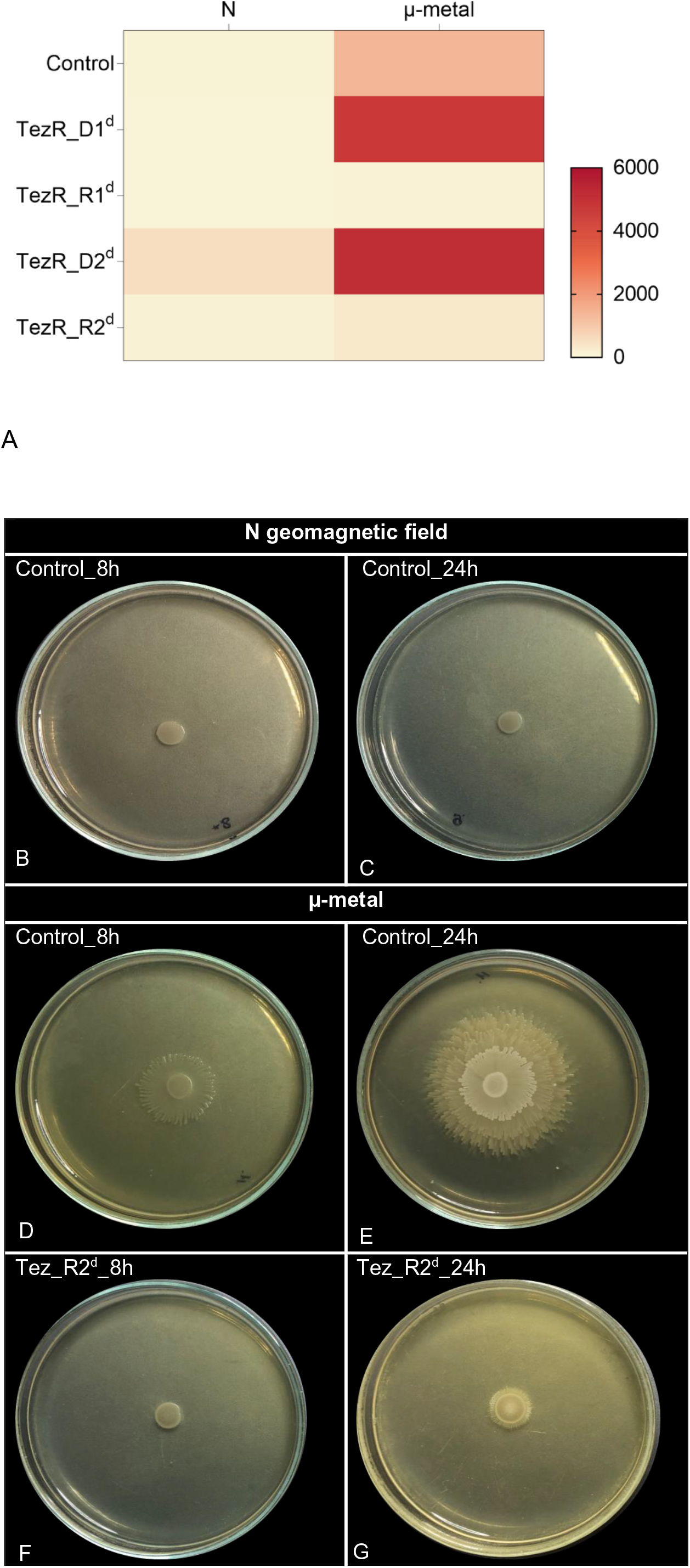
Role of TezRs in magnetoreception of *B. pumilus*. (A) Heat map representing the effect of TezRs loss on the size of the biofilm area under normal (N) and inhibited geomagnetic (µ-metal) fields after 24 h of growth. The size of the biofilm is represented by a color scale, from white (minimum) to red (maximum). (B–G) Dynamic changes to biofilm morphology in cells exposed to normal or inhibited geomagnetic field during 8 and 24 h of growth: (B, C) control *B. pumilus* under normal magnetic field; (D, E) control *B. pumilus* under inhibited (µ-metal) geomagnetic field; and (F, G) *B. pumilus* TezR_R2^d^ under inhibited (µ-metal) geomagnetic field.

Inhibition of the geomagnetic field promoted growth of control *B. pumilus* biofilms compared to cells grown under unaltered magnetic conditions (Fig. 9A, E). Loss of TezR_D1 or TezR_D2 stimulated bacterial growth in response to inhibition of the geomagnetic field across the entire plate (Fig. 9A). Instead, biofilms formed by *B. pumilus* following loss of TezR_R1 or TezR_R2 presented a strikingly diminished response to inhibition of the geomagnetic field. When compared with biofilms formed by control *B. pumilus*, those formed by *B. pumilus* TezR_R1^d^ or TezR_R2^d^ grown in a µ-metal cylinder for 24 h displayed only a negligible increase in size (Fig. 9A). However, they still exhibited minor changes in morphology compared with their counterparts grown under unaltered magnetic conditions (Fig. 9C, G).

To further elucidate the detailed role of RNA-based TezRs in sensing and responding to the geomagnetic field, we analyzed the time it took for morphological differences between control and *B. pumilus* TezR_R2^d^ biofilms placed in a µ-metal cylinder to occur. We found that already after 8 h, biofilms of control *B. pumilus* cultivated under inhibited geomagnetic field (Fig. 9D) presented an altered morphology with an increased size and irregular edge compared with those grown under normal conditions (Fig. 9B). In contrast, the morphology of *B. pumilus* TezR_R2^d^ biofilms was identical in the absence (Fig. 9F) or presence (Fig. 9B) of a regular geomagnetic field. These results showed that the alterations of biofilm morphology observed in *B. pumilus* TezR_R2^d^ in the inhibited geomagnetic field (Fig. 9G) occurred within 8–24 h. Together with our data pointing to the need for *S. aureus* for 8 h to restore normal temperature tolerance, these results add another line of evidence that bacteria started responding to geomagnetic field only after TezRs have been restored. Overall, RNA-based TezRs might be implicated in sensing and regulation of cell response to the geomagnetic field. These findings also highlight the complex web of interactions between different TezRs, as some of them adapt their regulatory role to the presence or absence of other TezRs.

### TezRs are required by bacteria for light sensing

Given the broad regulatory functions of TezRs in mediating the interaction between bacteria and the surrounding environment, we sought evidence for their biological relevance in sensing visible light. We analyzed differences in morphology of biofilms formed by control *B. pumilus* and *B. pumilus* following TezRs removal grown under light vs. dark conditions. Bacterial biofilms formed by either control *B. pumilus* or those lacking TezRs, except TezR_D2, responded to light by forming large biofilms with filamentous (filiform) margins (Fig. 10, Supplementary Fig. 2).

**Figure 10.**
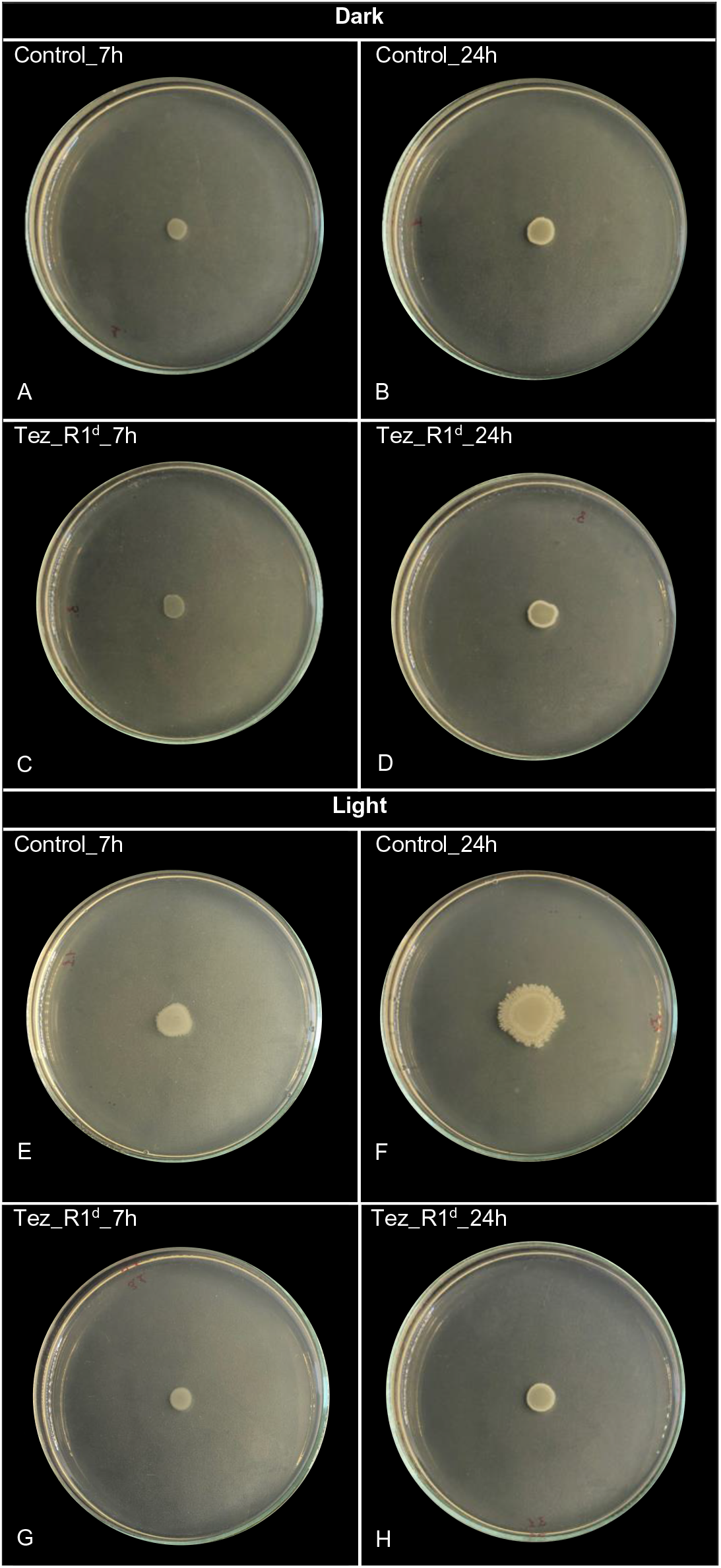
Role of TezRs in light sensing. (A - D) Images of (A, B) control *B. pumilus* (Control) and (C, D) *B. pumilus* TezR_R1^d^ (TezR_R1^d^) incubated in the dark for 7 h and 24 h. (E - H) Images of (E, F) control *B. pumilus* and (G, H) *B. pumilus* TezR_R1^d^ incubated in the light for 7 h and 24 h.

In contrast, *B. pumilus* TezR_D2^d^ grown under light exhibited reduced biofilm size compared to those grown under dark conditions (Supplementary Fig. 2). Strikingly, 24-h-old biofilms formed by *B. pumilus* TezR_R1^d^ and TezR_R2^d^ grown in the light presented altered margins, but their growth was contained compared with that of control *B. pumilus*.

As in the case of magnetoreception, we hypothesized that the reason for the observed phenotype was that *B. pumilus* TezR_R1^d^ and *B. pumilus* TezR_R2^d^ started responding to light only after 7 h, when either their RNA-based TezRs were restored or when the cell’s normal response was restored after TezR destruction. Therefore, we analyzed the morphology of 7-h-old biofilms grown under light conditions (Fig. 10). By that time, biofilms of control *B. pumilus* already had an altered morphology compared with those grown in the dark. In contrast, the morphology of *B. pumilus* TezR_R1^d^ was identical irrespective of illumination conditions. Accordingly, changes to biofilm morphology of *B. pumilus* TezR_R1^d^ occurred within 7–24 h of growth in the light, when TezR_R1 should have already been restored.

Together, the results imply that TezRs are involved in the regulation of microbial light sensing. Specifically, we found a positive association between the ability of bacteria to sense and respond to light, and the presence of RNA-based TezRs.

### TezRs regulate anaerobic survival of aerobes

Intuitively, we hypothesized that TezRs might regulate the bacterial response to a changing gas composition. To test this hypothesis, we used the obligate aerobe *P. putida*, generally known for its inability to perform anaerobic fermentation. Introduction of numerous additional genes, a massive restructuring of its transcriptome, and nutrient supplementation have been proposed as the only means to accommodate anoxic survival of this species (46–49).

Control *P. putida* and *P. putida* lacking TezRs were placed on agar and cultivated under anoxic conditions. While control *P. putida*, and *P. putida* deficient in TezR_D1 or TezR_D2 alone, or in combination with loss of RNA-based TezRs, could not grow under anaerobic conditions, loss of only RNA-based TezRs allowed for anaerobic growth of *P. putida* (Fig. 11A, B). *P. putida* TezR_R1^d^ and TezR_R2^d^ were characterized by microcolonies crowding (Fig. 11 A, B).

**Figure 11.**
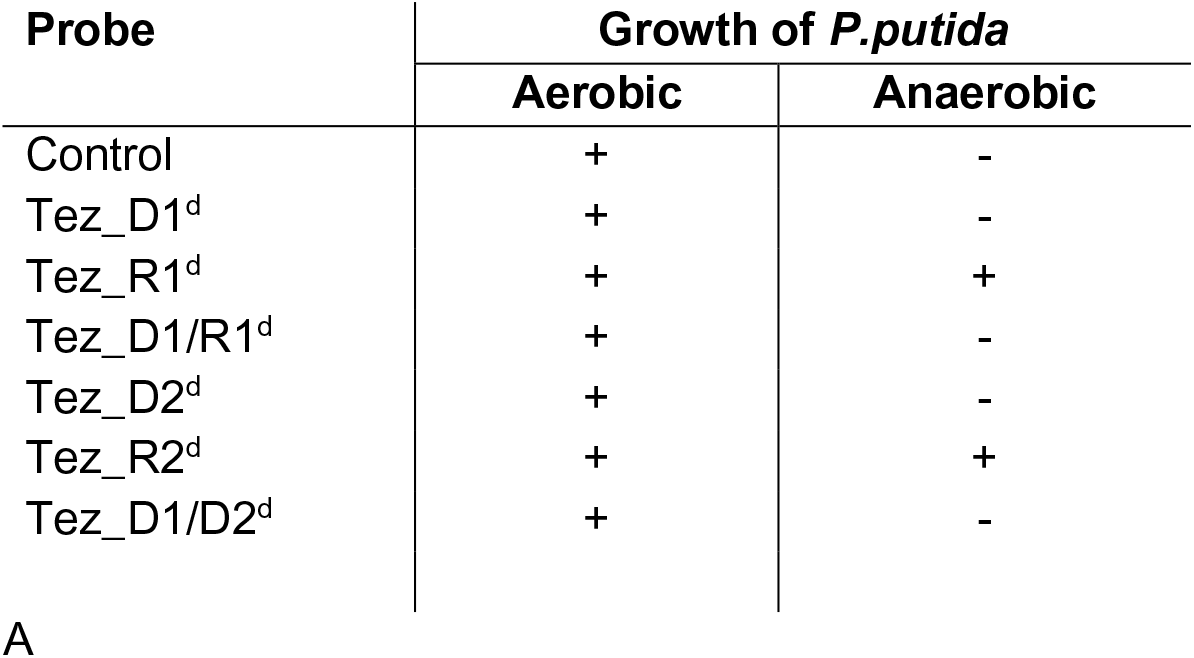
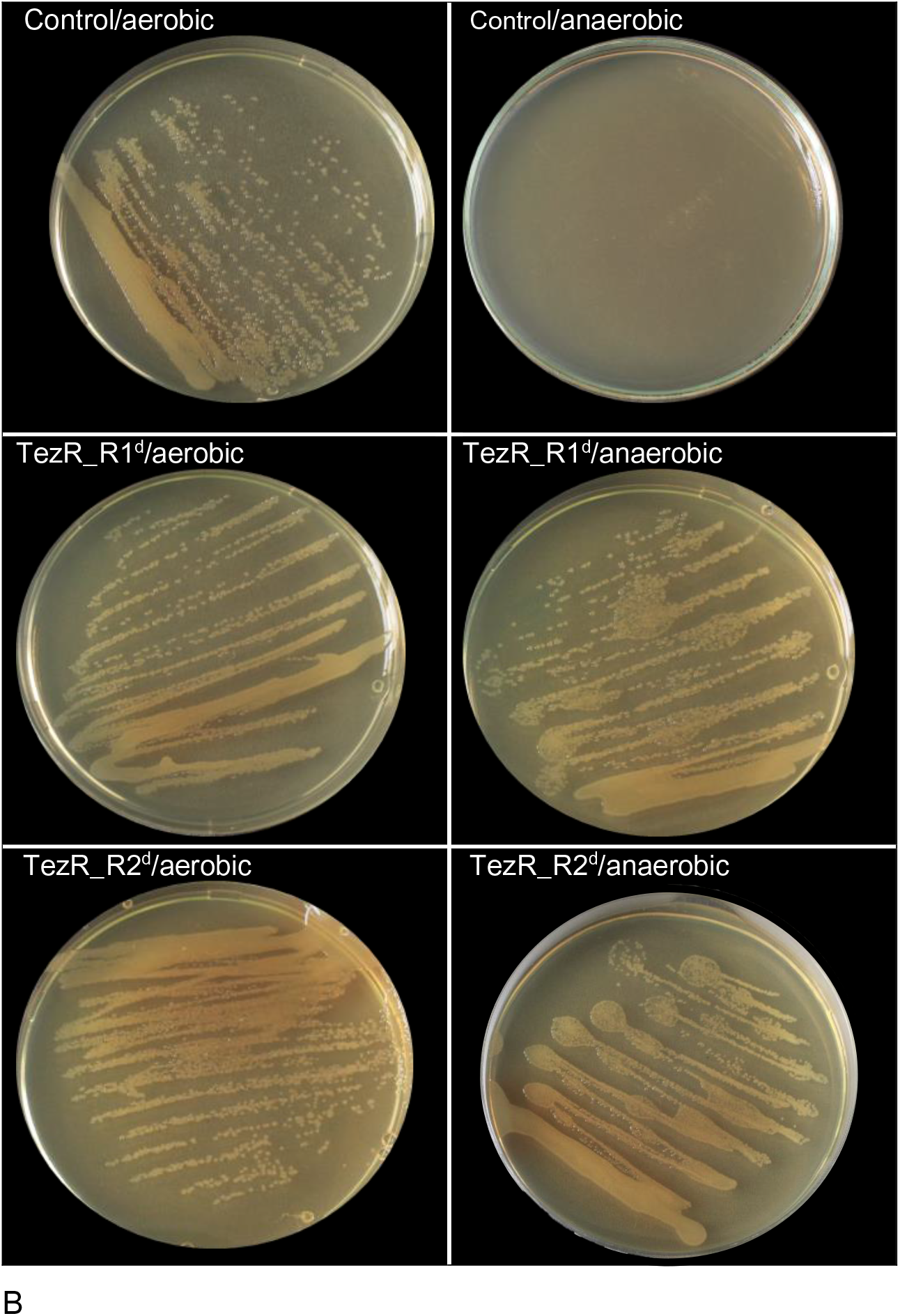
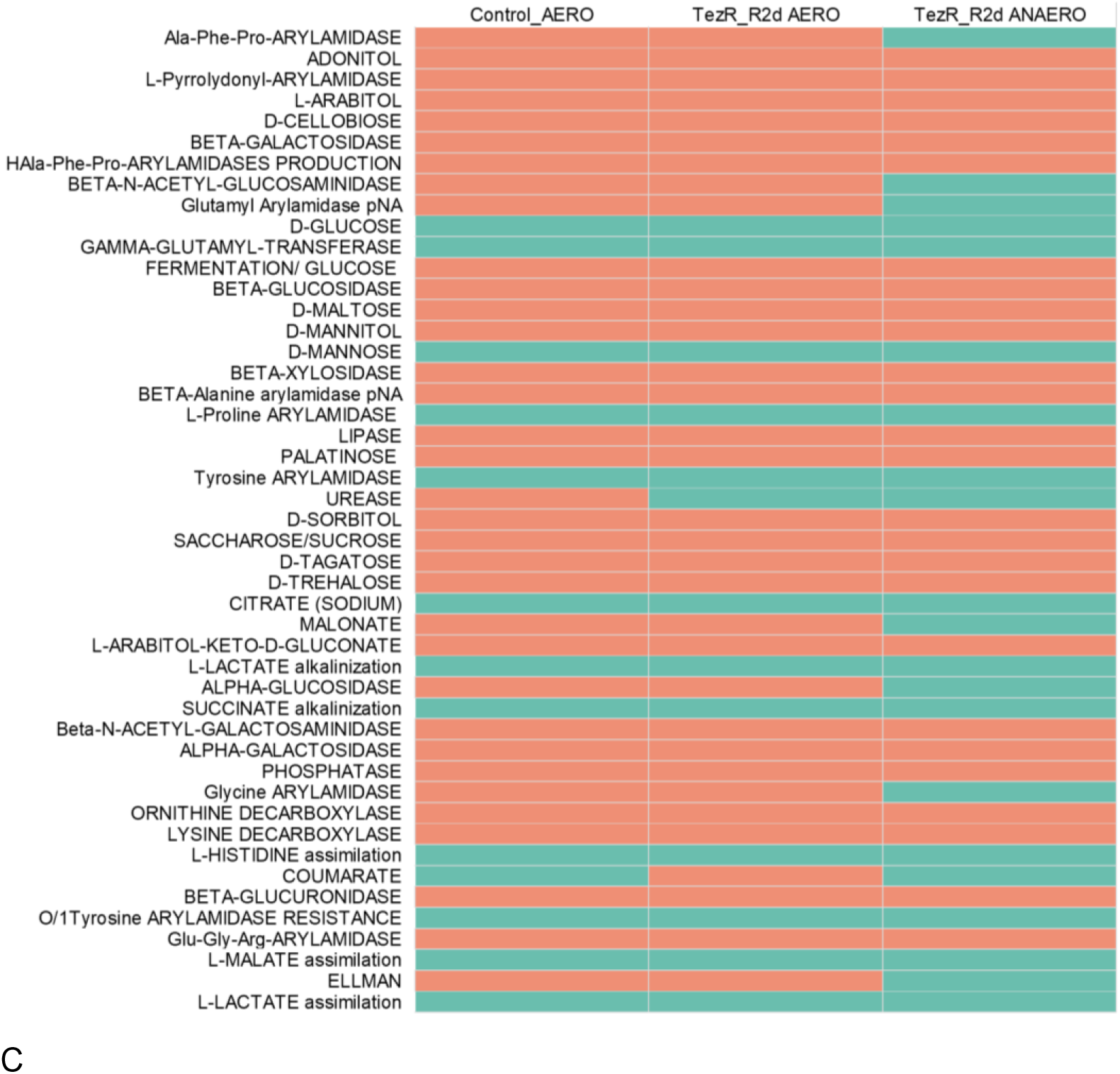
Role of TezRs in growth of *P. putida* under anaerobic conditions. (A) Effect of TezRs loss on the growth of *P. putida* under aerobic and anaerobic conditions. Presence of bacterial growth is marked with a “+” sign, absence of bacterial growth is marked with a “-“ sign. Values correspond to representative results of three independent experiments. (B) Growth of control *P. putida*, *P. putida* TezR_R1^d^, and *P. putida* TezR_R2^d^ under aerobic or anaerobic conditions for 24 h. (C) Biochemical profile of control *P. putida* grown under aerobic conditions (Control_aero) and *P. putida* TezR_R2^d^ cultivated under aerobic (TezR_R2^d^ _aero) and anaerobic (TezR_R2^d^_anaero) conditions in a VITEK® 2 system. Green color denotes positive test reaction results, red color denotes negative results. Values correspond to representative results of three independent experiments.

We compared the biochemical profile of *P. putida* TezR_R2^d^ grown in anoxic conditions with control *P. putida* and aerobically grown *P. putida* TezR_R2^d^ using the VITEK® 2 system (Fig. 11C). We observed activation of the urease enzyme in both aerobically and anaerobically grown *P. putida* TezR_R2^d^. This enzyme is considered essential for anaerobic fermentation in this species (46). Moreover, when *P. putida* TezR_R2^d^ were cultivated under anoxic conditions, we noted the activation of some aminopeptidases and glycolytic enzymes known to participate in microbial anaerobic survival in the absence of external electron acceptors such as oxygen (50–53).

Collectively, the findings point to a previously unknown sensing and regulatory function of the TRB-receptor system and, in particular, the role of TezR_R1 and TezR_R2 in adaptation to variations in gas composition. Importantly, loss of these TezRs enables obligatory aerobic *P. putida* to grow under anoxic conditions.

### Bacterial chemotaxis and biofilm dispersal are controlled by TezRs

Bacterial chemotaxis and biofilm dispersal are essential for colonizing various environments, allowing bacteria to escape stress, migrate to a nutritionally richer environment, and efficiently invade a host (54,55,56). Although *Bacillus* spp. is believed to rely on transmembrane chemoreceptors to detect environmental chemical stimuli and a kinase (CheA) and response regulator (CheY) to mediate downstream signals, it remains to be determined how the receptor senses such stimuli (57–59). Moreover, the gene network and signal transduction pathways controlling bacterial dispersal remain largely unexplored.

Here, we examined the role of TezRs in bacterial chemotaxis and dispersal in motile *B. pumilus* VT1200.

Control *B. pumilus* grew on the agar surface as round biofilms (Fig. 12A); however, addition of human plasma as a chemoattractant, triggered directional migration towards the plasma (Fig. 12B). Visual examination of biofilms revealed that *B. pumilus* TezR_D1^d^ lost their chemotaxis ability, while *B. pumilus* TezR_R1^d^ triggered biofilm dispersal within the chemoattractant zone (Fig. 12C–E). Biofilms formed by *B. pumilus* TezR_D2^d^ displayed marked chemotaxis towards plasma along with expanded biofilm growth, which appeared typical for this mutant even in the absence of chemoattractant (Fig. 12F). Loss of TezR_R2 induced marked biofilm dispersal towards the chemoattractant (Fig. 12G) and was accompanied by the formation of multiple separate colonies in the agar zone where plasma was added. Combined elimination of both DNA- and RNA-based secondary TezRs maintained biofilm expansion and chemotaxis behavior (Fig. 12H) typical of *B. pumilus* TezR_D2^d^; however, the primary community was characterized by zones of active sporulation (Supplementary Fig. 2).

**Figure 12.**
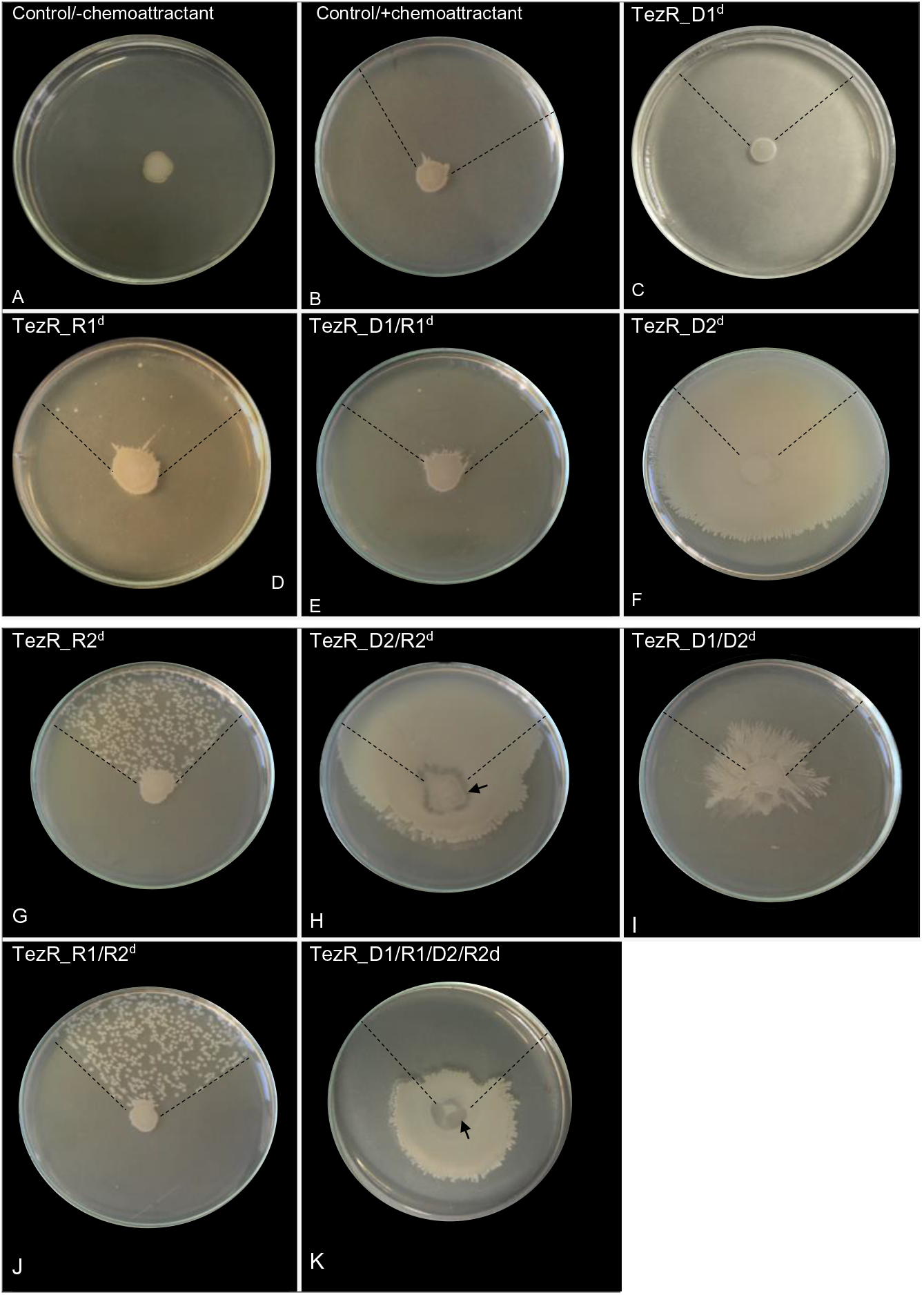
Effect of TezRs on *B. pumilus* chemotaxis to plasma and biofilm dispersal. (A) Control *B. pumilus* with no chemoattractant added. (B) Chemotaxis of control *B. pumilus* towards plasma as chemoattractant. (C) Chemotaxis of *B. pumilus* TezR_D1^d^. (D) Biofilm dispersal and chemotaxis of *B. pumilus* TezR_R1^d^. (E) Chemotaxis of *B. pumilus* TezR_D1/R1^d^. (F, H) Chemotaxis and visibly expanded biofilm growth of *B. pumilus* TezR_D2^d^ and *B. pumilus* TezR_D2/R2^d^. (G, J) Chemotaxis and intense biofilm dispersal of *B. pumilus* TezR_R2^d^ and *B. pumilus* TezR_R1/R2^d^. (I) Chemotaxis of *B. pumilus* TezR_D1/D2^d^. (K) Negative chemotaxis of *B. pumilus* TezR_D1/R1/D2/R2^d^. Black dotted lines denote the area in which plasma was placed. The black arrow points to zones of active sporulation. A chemotactic response is visualized as a movement of the biofilm away from the center towards the chemoattractant.

Interestingly, combined removal of primary and secondary DNA-based TezRs did not affect chemotaxis (Fig. 12I); however, *B. pumilus* TezR_D1/D2^d^ displayed geometrical swarming motility patterns with branched biofilm morphology, not observed in any other TezRs mutant of *B. pumilus*. Surprisingly, loss of all primary and secondary TezRs of *B. pumilus* prevented growth towards the chemoattractant, leading instead to negative chemotaxis away from plasma, and appearance of zones of active sporulation (Fig. 12K). These results point to the unique individual sensory and regulatory properties of TezRs in mediating chemotaxis, biofilm morphology, and dispersal. Biofilm dispersal triggered by the removal of TezR_R1 and TezR_R2 in the presence of chemoattractant occurred only in intact DNA-based TezRs. Hence, bacterial interaction with the chemoattractant is regulated by the TRB-receptor system through apparent cooperation between RNA- and DNA-based TezRs, as evidenced by the complex responses triggered by loss of multiple TezRs, and which cannot be accounted for by summing up the effect of individual TezRs losses.

### Functional responses induced by loss of TezRs include regulation of bacterial virulence

Membrane-damaging toxins that cause hemolysis or lecithin hydrolysis are critical for *S. aureus* virulence; however, regulation of their functioning remains poorly understood (60). In accordance with the observed pluripotent regulatory role of TezRs, we investigated the effect of TezRs loss on the hemolytic and lecithinase activities of *S. aureus* SA58-1. Loss of TezR_D1 or TezR_D2 alone, or in combination with other TezRs, statistically inhibited hemolysis (p < 0.05) and triggered the switch from α-hemolysis to β-hemolysis (Fig. 13A), pointing to the activation of genes encoding different hemolysins.

**Figure 13.**
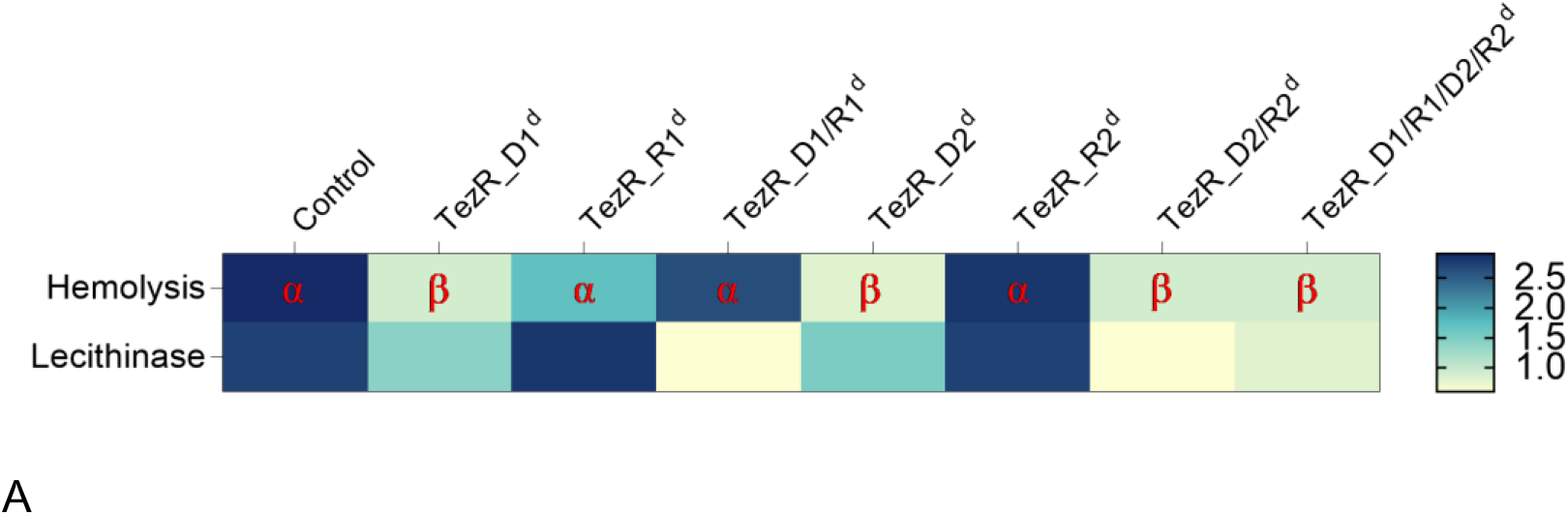
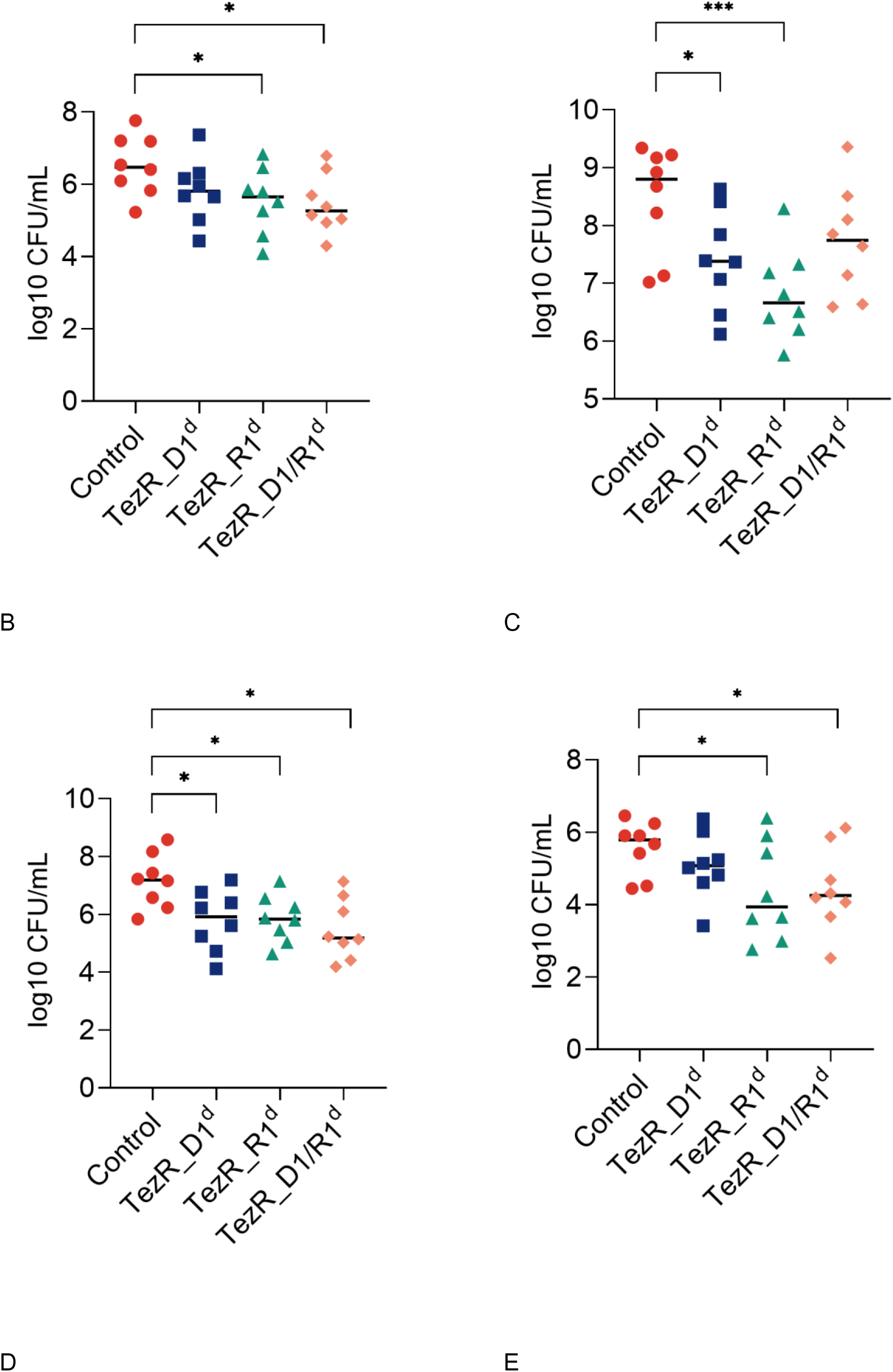
Role of TezRs in virulence. (A) Role of TezRs in the regulation of *S. aureus* hemolysis and lecithinase activities. Hemolytic activity of control *S. aureus* or *S. aureus* lacking TezRs is represented by a clear zone around the colonies on sheep blood agar plates. The presence of α- or β-hemolysis is marked with red letters. Lecithinase activity was analyzed by measuring a white diffuse zone surrounding the colonies. The extent of hemolysis and lecithinase zones (in mm) ranges from white (minimal) to dark blue (maximum). (B–E) Bacterial burden in animals intraperitoneally challenged either with control *S. aureus*, *S. aureus* TezR_D1^d^, *S. aureus* TezR_R1^d^ or *S. aureus* TezR_D1/R1^d^. Mice (n = 8) were euthanized 12 h after inoculation and *ex vivo* CFU were determined in (B) abdominal fluid, (C) liver, (D) spleen, and (E) kidneys. Values represent the mean ± SD. Each symbol corresponds to an individual mouse; horizontal bars denote the geometric mean. *p < 0.05, **p < 0.001.

A similar pattern was observed regarding the role of TezRs in regulating lecithinase activity (Fig. 13A), which was also inhibited following loss of DNA-based TezRs alone or in combination with RNA-based TezRs (p < 0.05). In contrast, loss of TezR_R1 or TezR_R2 alone caused no statistically significant alterations of hemolytic and lecithinase activities.

To further clarify the role of TezRs in virulence, we used a mouse model of *S. aureus* peritoneal infection. Mice were intraperitoneally challenged with 10.1 log10 CFU/mouse containing control *S. aureus*, *S. aureus* TezR_D1^d^, *S. aureus* TezR_R1^d^ or *S. aureus* TezR_D1/R1^d^ (Fig. 13B–E). All animals exhibited typical signs of acute infection within 12 h, including hypothermia, hunched posture and slightly reduced movement, piloerection, breathing difficulty, narrowed palpebral fissures, trembling, and reduced locomotor activity. Bacterial load was measured in the abdomen, spleen, liver, and kidneys 12 h post infection by aspiration from the abdomen or homogenization of organs, plating on selective *S. aureus* medium, and subsequent identification by microscopy.

Loss of any of the primary TezRs altered the host-parasite relationship, decreasing dissemination of *S. aureus*. The most pronounced decrease was observed in the liver, kidney, and spleen in the group challenged with *S. aureus* TezR_R1^d^. Reduction of *S. aureus* dissemination was less clear following infection with *S. aureus* TezR_D1^d^ or *S. aureus* TezR_D1/R1^d^, although it nevertheless resulted in a significant drop in viable counts in some organs. Taken together, these results imply that bacteria disseminated less effectively following loss of TezRs, which can be associated with their higher susceptibility to the host immune response or altered adaptation to the environment.

### Formation of bacterial persisters can be modulated by TezRs

To gain insight into how TezRs regulated the formation of persisters, we used *E. coli* ATCC 25922. Control *E. coli, E. coli* TezR_D1^d^, *E. coli* TezR_R1^d^, and *E. coli* TezR_D1/R1^d^ were normalized with respect to CFU, diluted in fresh ampicillin-containing medium, and incubated for 6 h (Fig. 14). The number of viable cells in the culture was determined by plating them on agar and overnight incubation.

**Figure 14.**
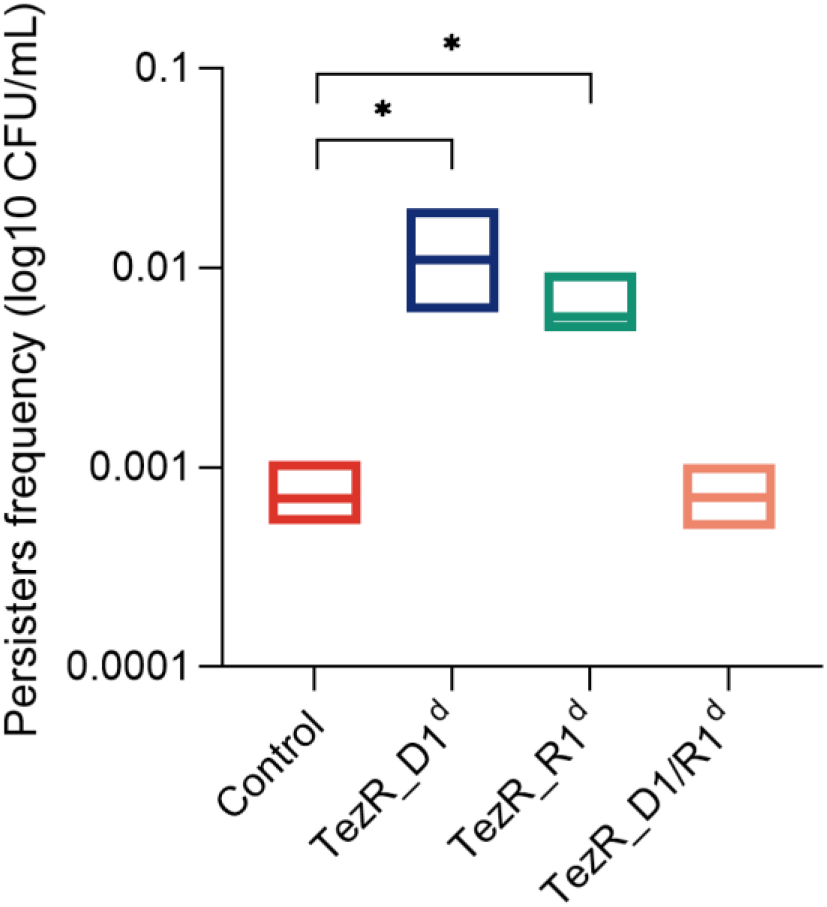
Impact of TezRs on persister formation. Control *E. coli*, *E. coli* TezR_D1^d^, *E. coli* TezR_R1^d^, and *E. coli* TezR_D1/R1^d^ were exposed to ampicillin for 6 h at 37 °C in LB broth and plated on LB agar without antibiotics to monitor CFU counts and colony growth. Values are representative of three independent experiments. Bars represent the mean ± SD. *p < 0.05.

As expected, only 1/1304 of original control *E. coli* cells were ampicillin tolerant. Primary TezRs regulated the rate at which cells entered dormancy and defined the persistence rate. The number of persisters was 155 times higher in *E. coli* TezR_D1^d^ and 8.5 times higher in *E. coli* TezR_R1^d^ (Fig. 14). Notably, the combined loss of both primary DNA- and RNA-based TezRs did not affect persister formation and there was no difference in the number of persisters between “drunk” *E. coli* TezR_D1/R1^d^ and the control.

### TezRs regulate spontaneous mutagenesis

Next, we examined how the destruction of different TezRs affected the rate of spontaneous mutagenesis. In these experiments, we measured spontaneous mutation frequency to rifampicin in *E. coli* ATCC 25922 by counting viable RifR mutants after cultivation on rifampicin-supplemented agar plates (Table 1). Spontaneous mutagenesis was inhibited in *E. coli* TezR_D1^d^, meaning that loss of TezR_D1 blocked the occurrence of replication errors, while loss of TezR_R1 did not affect this process. Surprisingly, the combined loss of TezR_D1/R1 triggered spontaneous mutagenesis and led to significantly more RifR mutants in “drunk” *E. coli* TezR_D1/R1^d^.

**Table 1.** Role of TezRs in spontaneous RifR mutagenesis.

| Probe | RifR mutants per 9<br>log10 *E. coli* cells<br>(mean ± SD)<sup>a</sup> | P value |
| --- | --- | --- |
| Control *E.coli* | 27 ± 5.79 |  |
| *E.coli* TezR_D1<sup>d</sup> | 0 ± 0 | 0.015 |
| *E.coli* TezR_R1<sup>d</sup> | 34 ± 8.84 | 0.249 |
| *E.coli* TezR_D1/R1<sup>d</sup> | 1050 ± 258.83 | 0.021 |

<sup>a</sup> Values represent the mean from at least three independent experiments.

### Loss of TezRs favors bacterial recombination

To determine the role of TezRs in bacterial recombination, we incubated control *E. coli* LE392 with λ phage (bearing Ampr and Kanr genes) for a time sufficient to cause phage adsorption and DNA injection. This was followed by treatment with nucleases to generate *E. coli* LE392 TezR_D1d, *E. coli* LE392 TezR_R1^d^, and *E. coli* LE392 TezR_D1/R1^d^ (61).

Control *E. coli* LE392 were incubated with λ phage, but were not treated with nucleases. Loss of any primary TezRs increased recombination frequency, as indicated by the increased rate at which phages lysogenized sensitive bacteria and, consequently, the higher number of antibiotic-resistant mutants (Fig. 15). The increase was statistically significant (p < 0.05) only in bacteria lacking TezR_R1 or those with combined loss of TezR_D1/R1. Taken together, these findings show that primary TezRs regulate recombination frequency and their loss can affect prophage formation.

**Figure 15.**
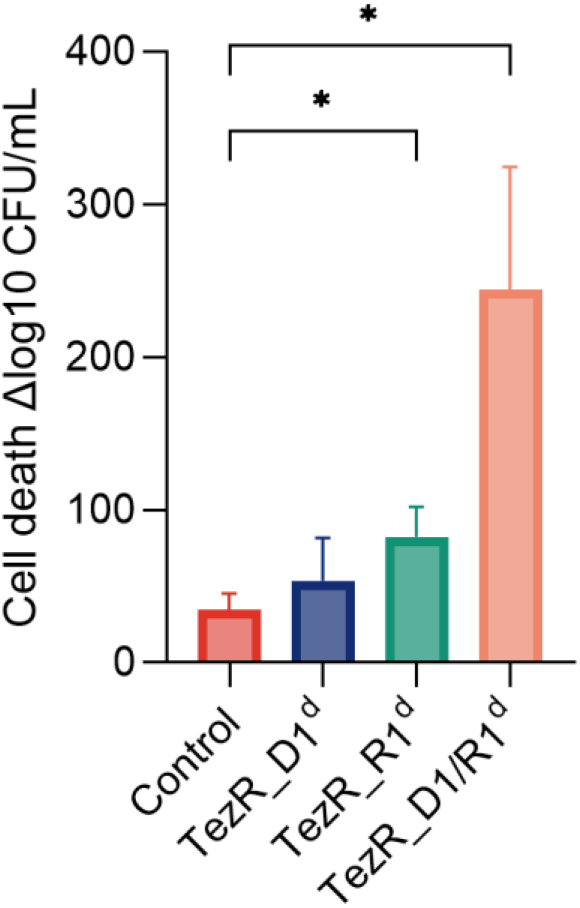
Role of TezRs in bacteriophage integration frequency. Data represent the mean of three independent experiments, error bars depict the standard deviation. *p < 0.05.

### TezRs are required for chemosensing and utilization of xenobiotics

To investigate the role of TezRs in xenobiotics sensing and utilization, control *B. pumilus* and *E. coli* or their counterparts lacking primary TezRs were inoculated in M9 minimal medium supplemented with the xenobiotic dexamethasone as the sole source of carbon and energy (62,63). We compared the lag phase, which comprises the time required for sensing and starting the utilization of these nutrients, between bacterial with unaltered and destroyed primary TezRs (64–66).

Loss of TezR_D1 in *E. coli* and *B. pumilus* did not affect the lag phase when bacteria were grown on media supplemented with dexamethasone. In marked contrast, the time lag of *E. coli* and *B. pumilus* devoid of TezR_R1 (Fig. 16A, B) was delayed by 3 and 2 h compared with that of control bacteria (p < 0.05), indicating a delay in the uptake and consumption of dexamethasone.

**Figure 16.**
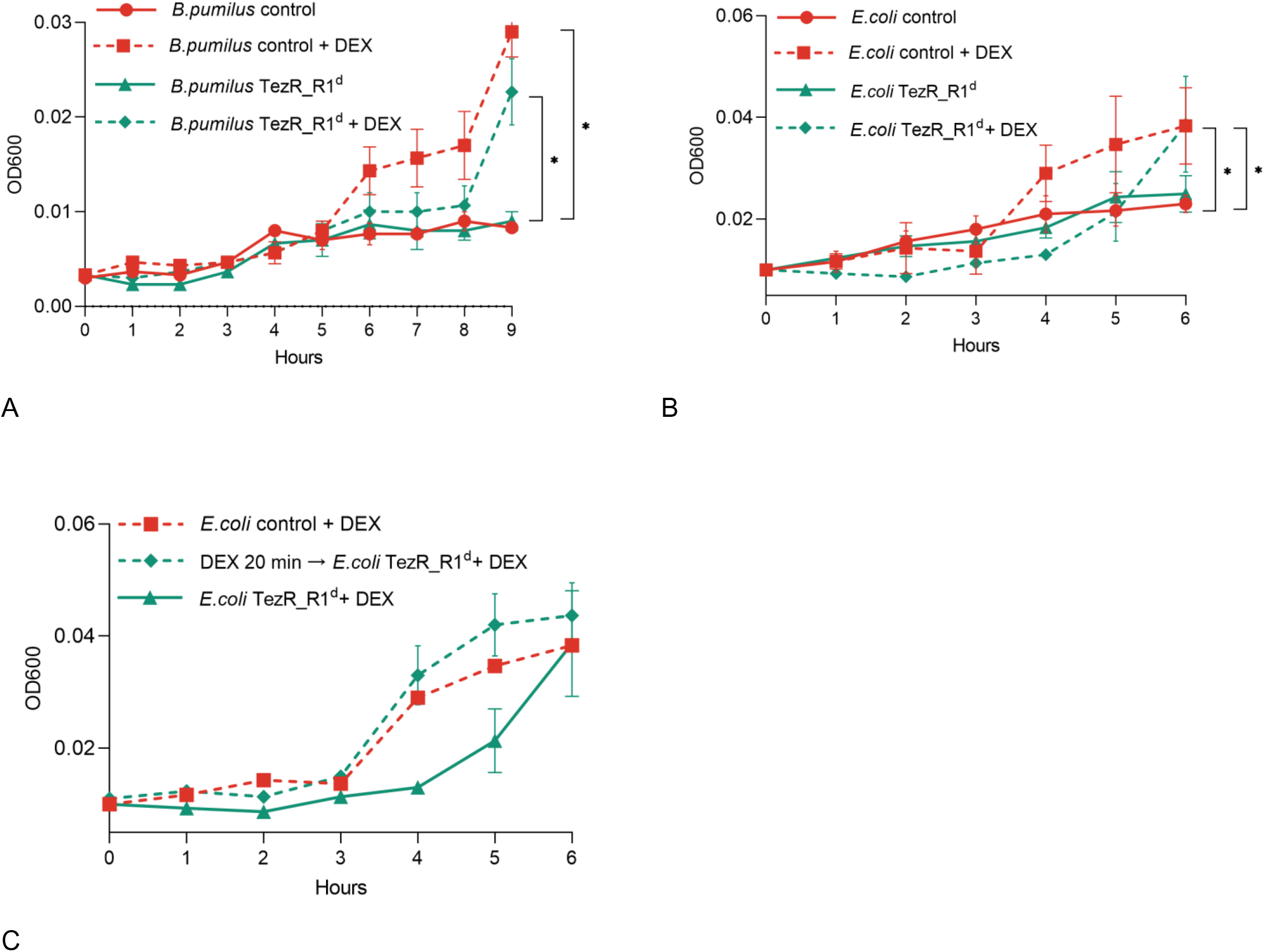
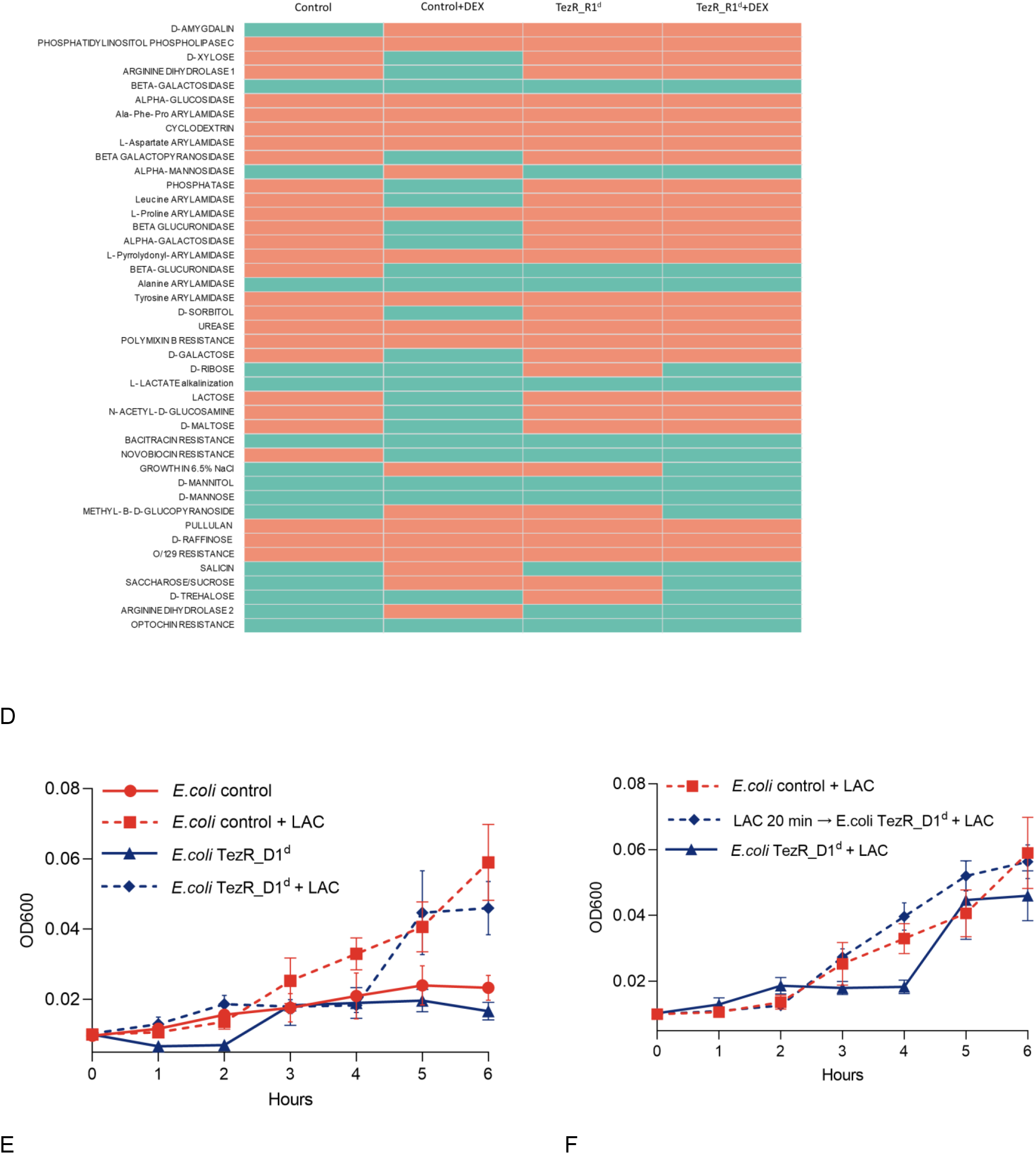
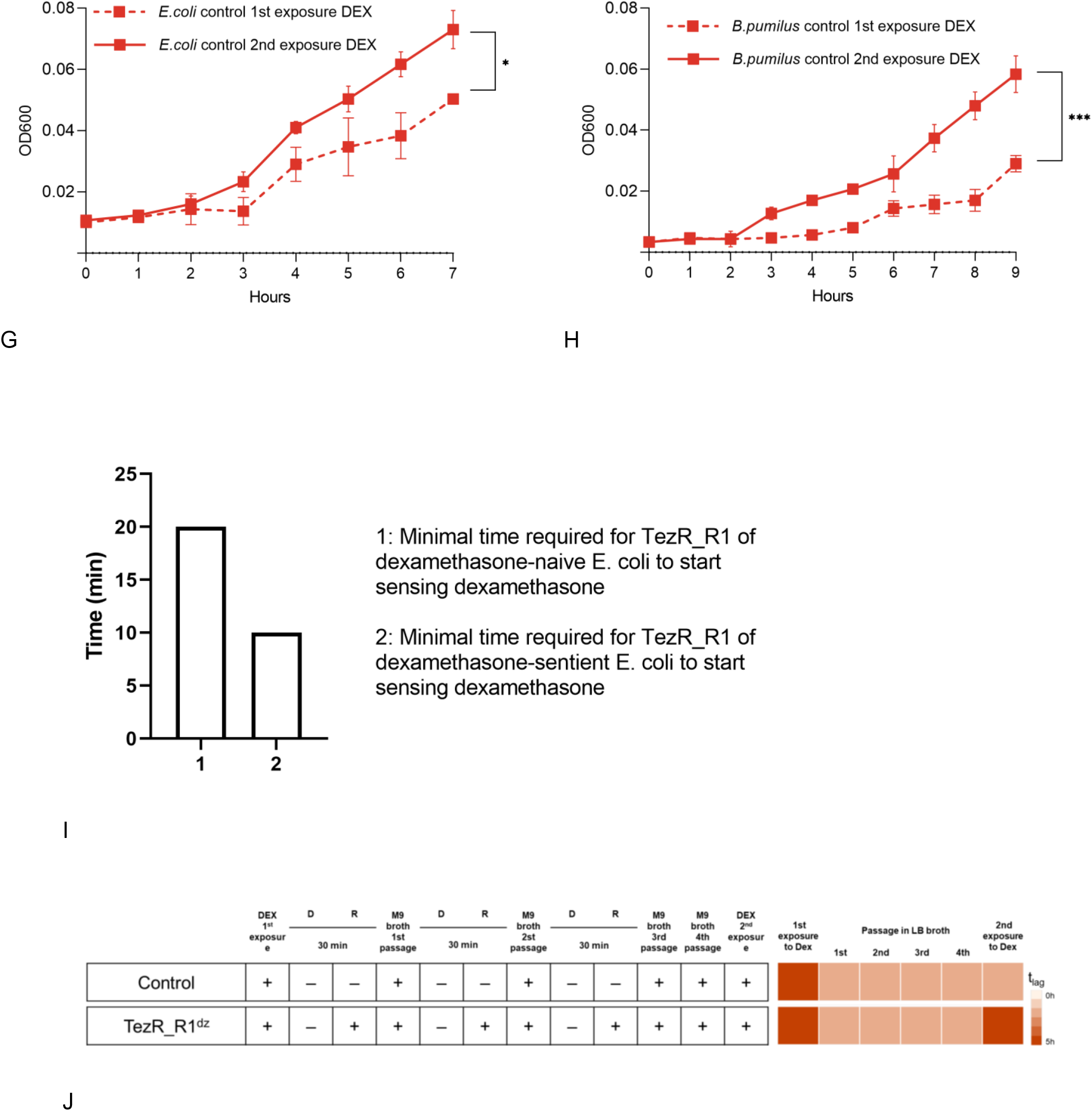
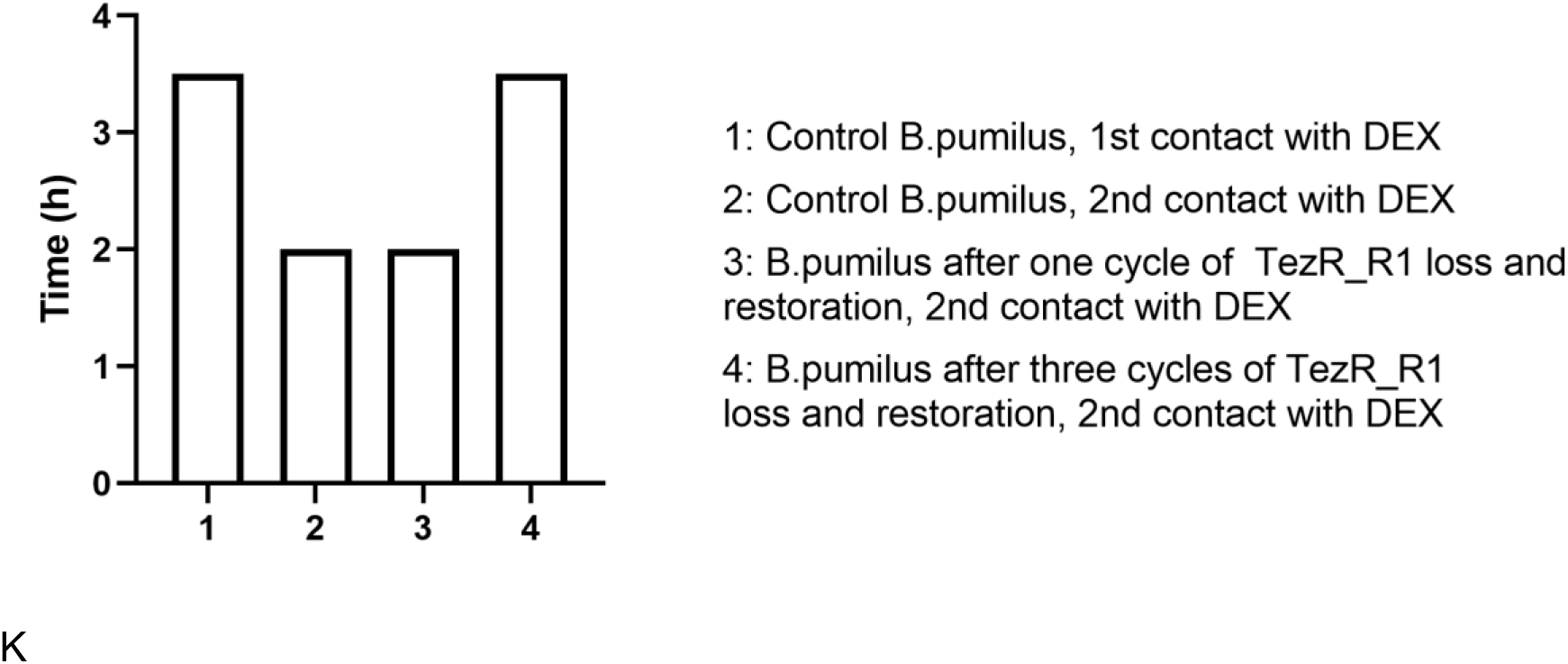
Role of TezRs in chemosensing and bacterial memory. Growth of control *B. pumilus* or *E. coli* and their counterparts lacking primary TezRs on M9 medium with and without dexamethasone (DEX) or lactose (LAC) was monitored over time. (A) Control *B. pumilus* and *B. pumilus* TezR_R1^d^ grown in M9 medium with or without dexamethasone. (B) Control *E. coli* and *E. coli* TezR_R1^d^ grown in M9 medium with or without dexamethasone. (C) Pretreatment of control *E. coli* with dexamethasone for 20 min followed by TezR_R1 removal and subsequent growth on M9 medium supplemented with dexamethasone. (D) Biochemical profile of control *B. pumilus* and *B. pumilus* TezR_R1^d^ grown on minimal M9 medium without (M9) or with dexamethasone (M9+DEX). Green denotes positive test reaction results, red denotes negative results. Values show representative results of three independent experiments. (E) Control *E. coli* and *E. coli* TezR_R1^d^ grown in M9 medium with or without lactose. (F) Pretreatment of control *E. coli* with lactose for 20 min followed by TezR_R1 removal and subsequent growth on M9 medium supplemented with dexamethasone. (G) Time required for dexamethasone-naïve and dexamethasone-sentient control *E. coli* to commence growth on M9 medium supplemented with dexamethasone. (H) Time required for dexamethasone-naïve and dexamethasone-sentient control *B. pumilus* to commence growth on M9 medium supplemented with dexamethasone. (I) Minimal time required for TezR_R1 of E. coli to start sensing dexamethasone. The X-axis represents the time lag of control *E. coli* upon initial and second exposure to DEX. (J) Time to the start of DEX utilization (tlag) by dexamethasone-naïve and dexamethasone-sentient *B. pumilus*. The experimental protocol is shown to the left. The tlag after each passage in M9 medium with or without dexamethasone is shown to the right as a heat map, whose color scale ranges from white (0 h) to red (5 h). (K) Minimal time required for TezR_R1 of *B. pumilus* to start sensing dexamethasone.

We hypothesized that the prolonged time required by bacteria lacking TezR_R1 to start using dexamethasone resulted from disruption of their role in sensing and nutrient consumption, rather than an alteration of transcriptional activity following their removal. To verify this hypothesis, we conducted an experiment designed to prove that if bacteria used TezR_R1 to sense dexamethasone, then *E. coli* pretreated with dexamethasone followed by TezR_R1 elimination and cultivation in M9 supplemented with dexamethasone would have the same time lag as wild-type *E. coli* in the same M9 medium. In other words, once bacteria sensed dexamethasone through TezR_R1, they would continue responding to it even if TezR_R1 was subsequently removed.

In agreement with this hypothesis, control *E. coli* exposed to dexamethasone for at least 20 min with subsequent TezR_R1 loss and inoculation in dexamethasone-supplemented M9 exhibited similar growth and time lag as control *E. coli* (Fig. 16C).

We also analyzed how loss of TezR_R1 altered the biochemical profile of *B. pumilus* grown on minimal M9 medium supplemented with dexamethasone (Fig. 16D). Addition of dexamethasone to control *B. pumilus* clearly induced a variety of enzymes known to participate in steroid metabolism including β-glucuronidase (67). This increase was less apparent in *B. pumilus* TezR_R1^d^, whereby no β-glucuronidase was detected. Lack of changes to the biochemical activity of bacteria devoid of TezRs following treatment with nutrients provides another line of evidence supporting the essential role of TezRs in the sensing and response to chemical factors, as well as recognition of xenobiotics.

### Utilization of lactose and functioning of the lac-operon are controlled by TezRs

To evaluate the potential universal role of primary TezRs in detecting exogenous nutrients, we examined their role in sensing lactose by cultivating the lac-positive strain *E. coli* ATCC 25922 in M9 medium supplemented with lactose as the sole source of carbon and energy. Surprisingly, unlike for dexamethasone, loss of TezR_R1 had no effect on lactose sensing. At the same time, loss of TezR_D1 increased the time lag by 2 h compared with control *E. coli*, indicating how utilization of lactose was regulated by these receptors (Fig. 16E). As with dexamethasone, when control *E. coli* were pre-exposed to lactose for 20 min, followed by TezR_D1 removal and subsequent cultivation on M9 medium supplemented with lactose, their behavior and time lag was similar to that of control *E. coli* (Fig. 16F). This finding further confirmed the lactose-sensing role of TezR_D1 and how functioning of the lac-operon relied on initial substrate recognition through TezRs.

### TezRs are implicated in bacterial memory and forgetting

We reasoned that, if TezRs participated in the sensing nutrients, they might also play a role in bacterial memory formation and verified this possibility using an ‘adaptive’ memory experiment (14, ^68^). We found that control *E. coli* and *B. pumilus* “remembered” the first exposure to dexamethasone, as indicated by shortening of the lag phase from 3 h upon first exposure to 2 h upon second exposure for *E. coli* and from 5 to 2 h for *B. pumilus* (Fig. 16G, H).

We next assessed whether TezRs implicated in the memorization of a previous engagement to nutrients required less time to trigger utilization of such a nutrient upon repeated sensing. To achieve the stated goal, we exposed “dexamethasone-naïve” and “dexamethasone-sentient” *E. coli* with unaltered TezRs to dexamethasone for different time periods. After that, TezR_R1 were destroyed and cells were placed in fresh M9 medium containing dexamethasone. Only the bacteria whose pre-exposure to dexamethasone prior to TezR_R1 destruction was enough to trigger its utilization were able to grow. In agreement with our hypothesis, we found that TezR_R1 required 20 min to sense and trigger the utilization of dexamethasone upon first exposure to it (Fig. 16I), but only 10 min upon second exposure (p < 0.05). The difference in time required for TezR_R1 to mount a response at first (20 min) and repeated (10 min) contact with dexamethasone points to the involvement of TezRs and the TRB-receptor system in long-term cell memory formation, enabling a faster response to repeated stimuli (^69^).

We next studied the role of TezRs in “forgetting”. We supposed that because TezRs participated in bacterial memory, their continued loss might result in no memory of past experiences, which would reflect in a longer time lag.

We found that control *B. pumilus* remembered the first exposure to dexamethasone, indicated by reduction of the lag phase from 5 h upon first exposure to 2 h upon second exposure. Dexamethasone-sentient *B. pumilus* with restored TezRs (following one- or two-time cycles of TezRs removal and subsequent restoration) maintained a time lag below 2 h (Fig. 16J), meaning that these one- or two-time cycles of TezRs loss did not affect bacterial memory. However, three repeated rounds of TezRs removal and restoration led to “forgetting” of any previous exposure to dexamethasone and the behavior of the corresponding *B. pumilus* became similar (5-h lag phase) to that of control *B. pumilus* upon first exposure to dexamethasone. We named these cells, whose memory had been erased by multiple cycles of TezRs loss “zero cells”.

Moreover, we found that after one or two-time removal of TezRs and subsequent restoration, TezRs continued to react faster to the substrate than at the very first contact (Fig. 16K). However, TezRs restored after three-time cycles destruction required the same contact time as naïve cells to sense the substrate. We reasoned that TezRs restored after one- or two-time cycles of destruction retained a type of “memory” (a reduced time required to sense and recognize substrate). This phenomenon appeared to depend on the role of TezRs in a bacterial intergenerational memory scheme capable of maintaining and losing past histories of interactions.

#### The effect of the binding of propidium iodine (PI) on the functionality of TezRs

To further confirm the role of TezRs in cell signaling we inactivated them using PI, which is known to bind both DNA and RNA without penetrating the live cells (70). Similar to the observation where both TezR_D1/R1^d^ were removed, PI-treated *B. pumilus* exhibited the identical pattern of increase in lag phase and delay in the uptake of dexamethasone when incubated in minimal media (Fig. 17). Thus, these results imply that not only TezRs destruction, but also abrogation of their functions by PI binding, modulates the sensory and regulatory activities of the cell.

**Figure 17.**
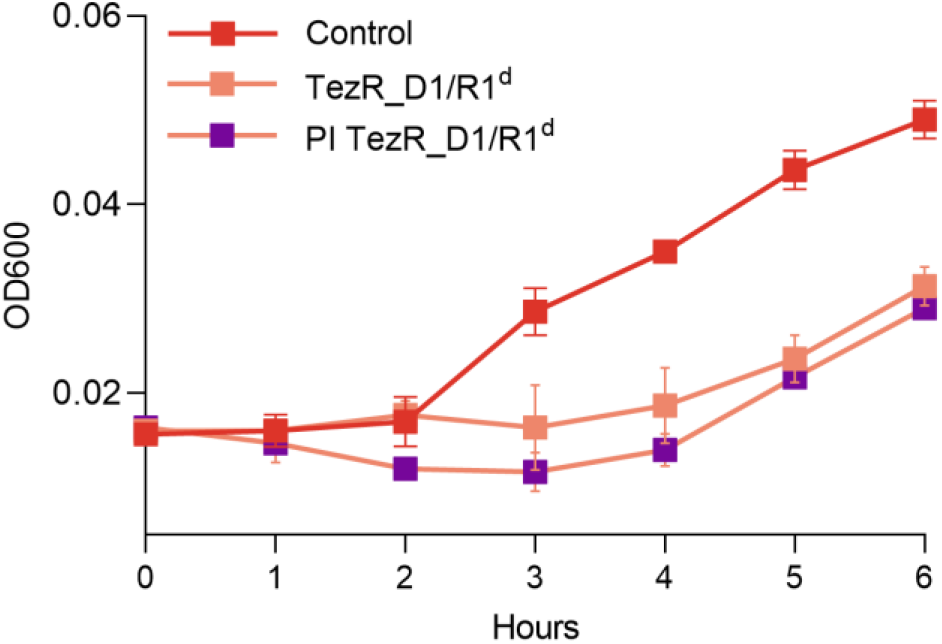
Inactivation of TezRs with PI. Time required for dexamethasone-sentient *B. pumilus* control (control), or *B. pumilus* following TezR_D1/R1 destruction (TezR_D1/R1^d^) or with TezR inactivated with PI (PI TezR_D1/R1^d^) to commence growth on M9 medium supplemented with dexamethasone.

### Role of reverse transcriptase and integrase in functioning of the TRB-receptor system

We hypothesized that formation and functioning of TezRs could be associated with reverse transcription and that affecting the corresponding enzymes might prevent the restoration of TezRs after their removal. Recent data suggest that non-nucleoside reverse transcriptase inhibitors (RTIs), originally designed to block HIV reverse transcriptase, interact non-specifically with different transcriptases (71,72). Here, we used non-nucleoside RTIs against control *S. aureus* and *S. aureus* lacking primary TezRs.

The RTIs etravirine and nevirapine did not exhibit any antibacterial activity against *S. aureus* and presented a MIC > 500 µg/mL (Supplementary Table 5). Thus, in this experiment we used very low doses of RTIs, more than 100 fold lower than their MICs.

Addition of RTIs to the medium did not alter growth dynamics of control *S. aureus* (measured as OD600), but affected growth of *S. aureus* lacking primary TezRs (Fig. 18A). Specifically, RTIs inhibited growth of *S. aureus* TezR_D1^d^ (p < 0.05 for all), but not *S. aureus* TezR_R1^d^. Even more surprisingly, treatment of *S. aureus* TezR_D1/R1^d^ with RTIs accelerated bacterial growth. We suggest that the inhibitory effect of RTIs on growth of bacteria lacking TezRs can be explained by the requirement for these receptors when cells are grown in liquid media.

**Figure 18.**
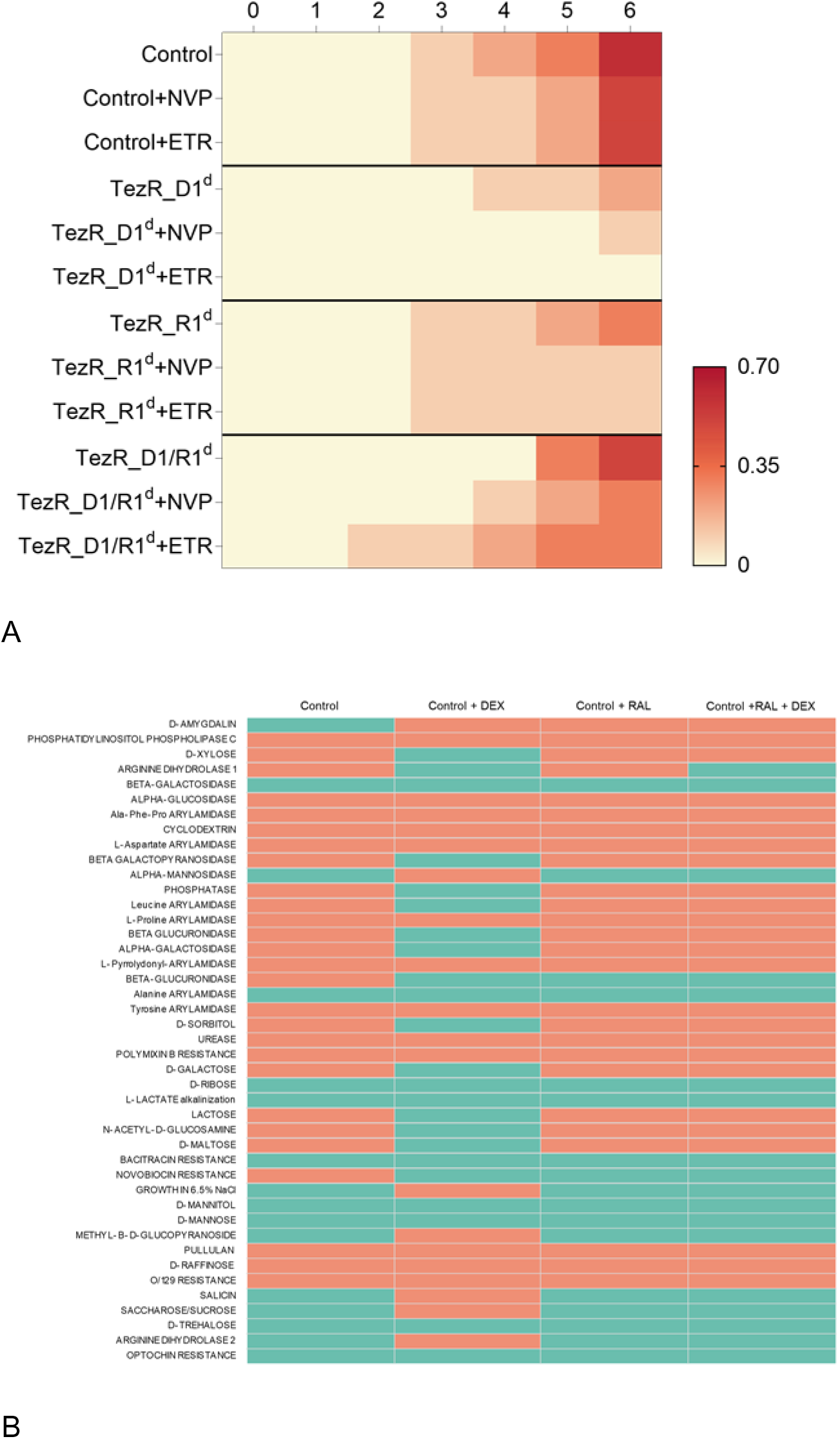
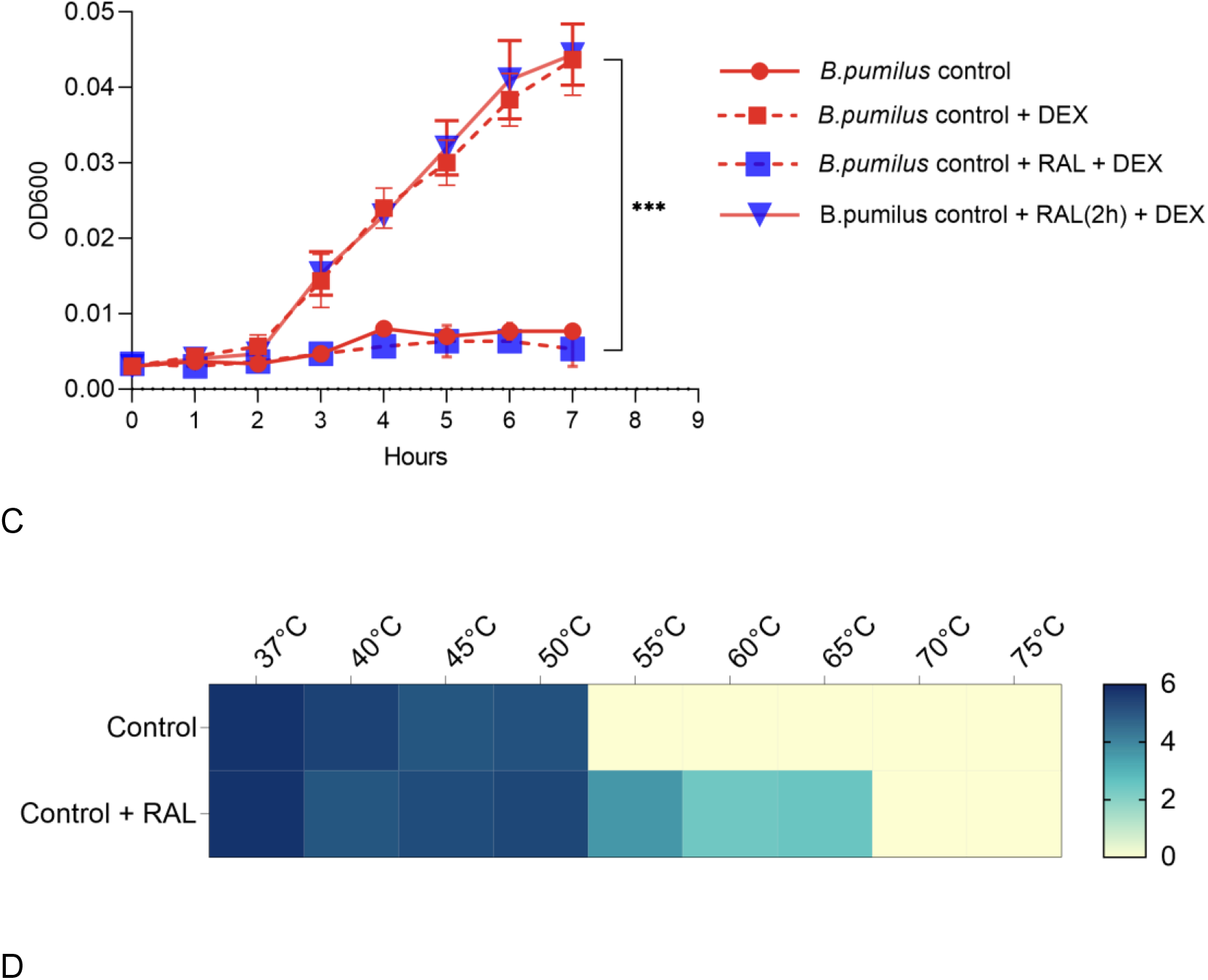
Role of reverse transcriptase and integrase in the TRB-receptor system. (A) Effect of RTIs on bacterial growth and memory. Heat map representation of growth by control *S. aureus*, *S. aureus* TezR_D1^d^, *S. aureus* TezR_R1^d^, and *S. aureus* TezR_D1/R1^d^ upon treatment with RTIs. Nevirapine (NVP) and etravirine (ETR) were added to the broth and OD600 was monitored hourly for 6 h at 37 °С. OD600 is labeled by a color scale, from white (minimal) to red (maximum). Values show representative results of three independent experiments. (B) Biochemical profile of control *B. pumilus* grown on M9 minimal medium without (M9) or with dexamethasone (M9+DEX), and with or without adding raltegravir (RAL). Green denotes positive test reaction results, red denotes negative results. Values show representative results of three independent experiments. (C) Raltegravir (RAL) added together with *B. pumilus* grown on M9 medium with dexamethasone (DEX) or 2 h after the plating of control *B. pumilus* on M9 with DEX (blue triangles with red line). (D) Heat map showing the effect of raltegravir on signal transduction from TezRs in relation to temperature tolerance in control and raltegravir-treated (control+RAL) cells. CFU are labeled by a color scale, from white (minimum) to blue (maximum). Values show representative results of three independent experiments.

Next, we investigated the onset of a signal transduction cascade following the interaction between TezRs and ligands. We hypothesized that the response to stimuli might also depend on recombinases. To verify this possibility, we used raltegravir, an inhibitor of viral integrase known to cross-react with bacterial recombinases due to structural and functional similarity with HIV integrase (73,74). Using a nontoxic concentration of raltegravir (Supplementary Table 5), we successfully blocked the activation of bacterial enzymes of control *B. pumilus* in response to dexamethasone (Fig. 18B). As a result, the biochemical profile of control *B. pumilus* grown on M9 medium supplemented with dexamethasone and raltegravir was almost identical to that of *B. pumilus* grown on M9 without dexamethasone. This allowed us to assume that raltegravir blocked signal transduction from TezRs following substrate recognition.

To confirm that the raltegravir-inducted response of *B. pumilus* to dexamethasone was not the result of any toxic effect, we measured OD600 of control *B. pumilus* when raltegravir was added to the medium at different time points (Fig. 18C). Addition of raltegravir to dexamethasone-sentient control *B. pumilus* grown on M9 with dexamethasone led to inhibition of bacterial growth only when it was added together with the cells, but lost its inhibiting function if added 2 h after growth had started (Fig. 18C). We believe that raltegravir inhibited signal transduction from TezRs occurring during the first 2 h, but had no control over it once the signal had already been relayed.

Given that we previously showed how the loss of TezRs enhanced survival at higher temperatures, we hypothesized that raltegravir might block signal transduction from TezRs and lead to higher heat tolerance even in bacteria with intact TezRs. *S. aureus* treated or not with raltegravir were gradually heated up to 65 °C and the presence of viable bacteria was analyzed. *S. aureus* treated with raltegravir could survive at temperatures over 15 °C higher than those of cells not treated with raltegravir (Fig. 18D). These data add another line of evidence supporting the involvement of the TRB-receptor system in intracellular signal trafficking.

## DISCUSSION

Here, we describe for the first time the most external receptive system in bacteria, named “TRB-receptor system”, which oversees almost all aspects of cell behavior and memory. Such a universal receptive system, implicated in sensing a wide range of chemical, physical, and biological factors, has not been described previously in eukaryotes or prokaryotes.

The system is composed of previously uncharacterized nucleic-acids based receptors capable of sensory and regulatory function, as well as reverse transcriptases and integrases. Our study shows a unique composition of these receptors, which we named TezRs. In contrast to known receptors formed by proteins, TezRs are formed by DNA and RNA molecules (75). The selective removal of different TezRs led to individual alterations in cell functioning and remarkably impacted the transcription of various genes, which highlights the specific role of each of the discovered TezRs.

We first showed that the TRB-receptor system functioned robustly across different bacterial types and played a previously unexplored and critical role in the regulation of microbial growth in liquid and solid media, as well as in collective behavior. These processes are known to be tightly regulated by numerous genes and post-transcriptional events (76). Loss of different TezRs resulted in changes to growth kinetics, biofilm formation, and cell size. The most significant alterations were noted for biofilms formed by motile bacteria lacking TezR_D2. These biofilms were characterized by formation of dendritic-like colony patterns, typical of cells with an increased swarming motility (77). Given that swarming motility is a hallmark of bacterial multicellularity, it is possible that TezRs participate in the regulation of this process (78).

Biofilm dispersal allows bacterial cells to leave a biofilm and migrate to a more favorable environment for resettlement. Previous evidence suggests that biofilm dispersal is modulated by the alteration of environmental conditions or gene activity (56,80,75). However, our data validated that this process is also modulated by TezRs, without any other direct dependency on exogenous or endogenous genetic stimulus.

Furthermore, we observed that TezRs controlled sporulation, which represents another important bacterial indicator of the interaction with the external environment (81). The TRB-receptor system exerted divergent effects on sporulation and loss of particular TezRs could either increase or totally inhibit this process. Given that sporulation is a stressful event for the cell and its initiation is tightly regulated in response to an unfavorable environment, increased sporulation following loss of TezR_D1, TezR_R1, and particularly TezR_R2 raises the question of the role of these TezRs in such context (82). The inhibition of sporulation following removal of TezR_D2 under normal and stressful conditions adds another line of evidence suggesting that TezRs supervise known regulatory pathways and known receptors responsible for sporulation.

We also found that primary TezRs regulated the rate at which cells entered dormancy and determined the persistence rate, thus defining a bet-hedging strategy of cells. Even though the molecular mechanisms underlying persister formation have been intensively studied and are believed to be achieved through the modulation of multidrug efflux pumps, DNA repair, and ROS production, as shown here the understanding of this important phenomenon is incomplete (83–85).

To evaluate the role of the TRB-receptor system in bacterial adaptation to a variety of chemical and physical factors, we began by looking at the regulation of bacterial survival at high temperatures. In a set of experiments, we showed that all primary TezRs and TezR_R1 in particular were key regulators of survival under thermal stress and their removal enabled cells to tolerate up to 20°C higher temperatures than those managed by control bacteria. We reasoned that, because loss of TezRs before the heating step but not after it increased survival, TezRs might be involved in thermosensing and supervise the corresponding response. This idea was supported by the notion that intracellular mRNA and RNA thermosensors could react to an altered temperature, which thus modulated translation (86). We found that the TRB-receptor system orchestrated the cell response to UV exposure. When bacteria are exposed to UV light, they respond to DNA damage by a highly regulated series of events known as the SOS response, which ultimately dictates whether the cell should survive or induce cell death (87,88). Loss of RNA-based TezRs increased survival after UV exposure, which can be explained by modulation of SOS-induced cell death (89).

An interesting finding regarding the regulation of cell responses to variations in gas composition was observed when the obligate aerobe *P. putida* could grow under anoxic conditions following the removal of TezR_R1 or TezR_R2. Notably, this finding is echoed by recent theoretical studies suggesting that growth of *P. putida* under anoxic conditions would require numerous additional genes and a massive restructuring of its transcriptome to find alternative means of ATP synthesis (46,48). We reasoned that RNA-based TezRs could be implicated in sensing of the gas content or stimulate genetic variability to enable the selection of clones capable of growing under anoxic conditions.

Examining the bacterial response to other physical factors, we found that TezRs were involved in sensing and regulation of the response to changes in the geomagnetic field (known as magnetoreception) and light. Non-magnetotactic and non-photosynthetic *B. pumilus* with intact TezRs sensed inhibition of the geomagnetic field and the presence of light in the environment, as manifested by changes in biofilm morphology and expanded growth. We found that RNA-based TezRs are implicated in sensing of the geomagnetic field and light and that, in the case of their loss, bacteria could not start responding to alterations in these factors for a few hours, most likely until these TezRs were restored. It is surprising, since until now, the identity of a magnetic sensor in non-magnetotactic bacteria remained enigmatic; however, some studies show that different bacteria even lacking magnetosomes are capable of sensing the geomagnetic field (25,26).

Interestingly, the ability of TezRs to interact with the magnetic field could be explained by the nucleic-acid structure of these receptors, owing to the alleged paramagnetic properties of nucleic acids and their ability to emit or transmit electromagnetic waves (90–94).

It has not escaped our attention that the observed altered responses to these physical factors by bacteria lacking RNA-based TezRs happened only as long as DNA-based TezRs were present. It is possible that different TezRs interact with each other to form functional complexes in which they affect each other’s functioning. This observation corroborates the fact that selective or combined removal of various TezRs triggered different transcriptomic clustering. Notably, the most significant impact on the transcriptome profiles, with the upregulation of the highest number of genes was triggered by the individual loss of RNA-based TezRs.

Studying the role of the TRB-receptor system in response to different chemical and physical factors, we were surprised by how cells lacking both RNA- and DNA-based TezRs continued responding to some of these factors. Although TezR_D1/R1^d^ bacteria displayed an increased survival at higher temperatures, their survival did not differ from that of control cells under altered UV, light, and gas content conditions. Indeed, combined cleavage of different TezRs triggered individual responses that were often more than just the sum of alterations triggered by the loss of each individual TezR. Thus, we named cells lacking primary DNA- and RNA-based TezRs that exhibited an unexpected response to stimuli “drunk cells.” The paradoxical behavior of “drunk cells” could be explained by the existence of internal (i.e., cytoplasmic) TezRs (TezR_i), which could be activated following the loss of primary TezRs. The existence of cytoplasmic receptors in bacteria was only recently shown, but these receptors are protein-based and respond only to chemosensing (95).

The present results also expanded our understanding of the TRB-receptor system in the control of mutational events and recombination frequency. We found that TezRs regulated spontaneous mutations and that it was possible to either inhibit this process through loss of TezR_D1 or increase it via combined removal of TezR_D1/R1. We did not look deeper into this phenomenon; however, we believe that alterations of these TezRs could possibly control the mismatch repair system, which is known to be responsible for spontaneous mutagenesis (96). The control of bacterial variability by the TRB-receptor system is also supported by increased recombination frequency following TezRs destruction during infections of bacteria by phages (97).

Our findings support a role for TezRs in microbial virulence and pathogenicity. TezRs regulate production of virulence factors, such as hemolysin and lecithinase, as well as *in vivo* bacterial dissemination. These properties are known to play an important role in the spreading of infections, but their underlying molecular mechanisms are only now beginning to be elucidated. In fact, given that loss of TezRs inhibited bacterial dissemination, nucleases produced by macroorganisms could actually constitute a protective mechanism (98,99).

Finally, we studied the role of TezRs in bacterial chemotaxis, which is one of the primary means of bacterial adaptation (37). We found that TezRs controlled chemotaxis and that removal of certain TezRs could either promote or inhibit this process, or even cause a switch from positive to negative chemotaxis. Because the loss of TezRs did not affect bacterial motility but modulated chemotaxis, we conclude that TezRs control and oversee the function of transmembrane methyl-accepting chemotaxis proteins, which are believed to be the primarily regulators of chemotaxis (100). In bacteria, chemotaxis can be viewed as an intrinsic element of chemoreception. Thus, not surprisingly, we discovered that TezRs played a primary role in both processes. We found that the existence of TezRs was a prerequisite for different bacteria to utilize well-recognized factors such as lactose, as well as synthetic xenobiotics. The fact that lactose utilization, which is one of the most well described examples of chemoreception, depends on TezRs can be explained by the overseeing function of the TRB-receptor system over the lac-operon. We reasoned that the controlling role of TezRs in sensing different substances including xenobiotics suggested how different bacterial chemoreceptors were under the control of the TRB-receptor system.

The ability of cells to sense environmental factors and nutrients is also related to cell memory. Participation of DNA- and RNA-based TezRs in cell memory formation to known nutrients and xenobiotics was supported by the difference in time required to sense and trigger substrate utilization by naïve and sentient bacteria. Given that genome rearrangement occurs during cell memory formation, we suggested, and for the first time confirmed, that bacterial memory formation could be blocked by recombinase inhibitors (101). Together, these results highlighted how loss of TezRs could modulate genome rearrangement during bacterial memory formation (101). Intriguingly, our results showed that TezRs of sentient bacteria exhibited faster substrate recognition than naïve cells and that this difference could be passed on through multiple generations. It is tempting to speculate that TezRs of sentient cells do not only maintain a memory of previous interactions, but that they also exhibit faster substrate recognition, which implies a selection of cells whose TezRs have higher affinity for previously encountered substrates. This characteristic shares similarity with the adaptive strategy of immune cells, whose secondary and more pronounced response is based on their affinity for antigens and the higher number of cells possessing relevant receptors (102–104).

We hypothesized that cell memory formation included several processes. First, substrate is sensed by TezRs. Second, this event triggers gene expression or rearrangement to utilize the substrate. Third, TezRs with a memory of this substrate and ability to recognize it in follow-up contacts are formed. The formation of TezRs with memory to previous events was proved by the possibility to erase this memory via loss of TezRs in substrate-sentient cells. Indeed, three repeated rounds of TezRs loss led to “forgetting” of the initial contact with the substrate. We named such cells “zero cells”. “Zero cells” did not “remember” previous interactions with the substrate and required the same time to start its utilization as substrate-naïve cells. We concluded that removing TezRs and forming “zero cells” altered the activity of genes or triggered genetic networks rearrangements. Therefore, we report for the first time that, by affecting TezRs, it is possible to control memory formation and “forgetting”, both of which are critical aspects of memory regulation. This finding opens a wide range of possibilities for directed cellular programming (105).

To address the question of how TezRs were formed, we hypothesized that this process involved different types of DNA and RNA transcription events (106). Even though reverse transcriptases have been found in a wide range of bacteria, their structure and function remain enigmatic (107). Bacterial retroelements with reverse transcription activity (mainly represented by group II introns associated with the CRISPR-Cas system), diversity-generating retroelements (producing hypervariable proteins mediating adaptation to a changing environment), Abi-related reverse transcriptases, and retron reverse transcriptases encoding extrachromosomal satellite multicopy single-stranded RNA/DNA structures remain all poorly understood (108–111). In addition, there are various reverse transcriptases of unknown function. In support of this idea, we observed inhibition of bacterial growth when cells lacking primary DNA-based TezRs (and not control, vehicle-treated cells) were treated with reverse transcriptase inhibitors. Accordingly, we speculated that this occurred due to inhibition of TezRs restoration by reverse transcriptases.

We have not specifically investigated the mechanism of TezRs translocation to the cell surface, but the observed upregulation of proteins associated with type VII secretion system (T7SS) following the loss of DNA-based TezRs alone or in combination with RNA-based TezRs, raises the question about T7SS involvement in translocation of DNA-based TezRs. Although, T7SS has not yet been fully characterized, and the intricate molecular mechanisms underlying its function remains elusive, the T7SS secretion machinery is attributed to bacterial pathogenicity and is also known to be a part of curli biogenesis machinery that requires extracellular DNA (112,113).

Trying to answer the question of how the signal from TezRs was processed further downstream in the cells, we found that the integrase inhibitor raltegravir blocked the bacterial response to the xenobiotic dexamethasone (74). As consumption of the latter was found to be controlled by TezRs, this finding suggested that bacterial recombinases might be implicated in the processing of stimuli from TezRs. Taken together, these results allowed us to conclude that recombinases and reverse transcriptases were part of the TRB-receptor system.

Taking into consideration the nucleic acids-based chemical nature of TezRs, it is worthwhile revisiting some of the existing paradigms of microbiology associated with nucleic acids. Thus, some biological effects so far associated with the action of nucleases against bacterial biofilms and inhibition of bacterial adhesion, might actually stem from previously overlooked changes to TezRs with subsequent loss of their receptive and regulatory function (44,114,115). Our data might also shed the light on the role of nucleic acids identified on cell surfaces, which have been described in some organisms but their contribution to cell functioning remained poorly defined (116–118).

The model used in this study and based on the use of nucleases to remove TezRs relevant to natural conditions. Many bacteria secrete nucleases in the extracellular environment, suggesting that the destruction of TezRs may be a conserved and previously overlooked mechanism to gain a fitness advantage over competing strains (99,119).

Along with the nucleases we used PI which is known to bind both DNA and RNA without penetrating live cells (120). As expected, bacteria following PI treatment behaved similarly as the “drunk cells” after the destruction of primary DNA and RNA formed TezRs. Therefore, not only their destruction but also their inactivation due to PI binding could significantly affect the receptive and regulatory functions of TezRs.

Future studies of the TRB-receptor system will require the development of new tools, coupled with an interdisciplinary approach that bridges microbiology and molecular biology. They should focus on the structural aspects of TezRs, as well as the molecular mechanisms of their formation and translocation to the cell surface. The functioning of bacterial TezRs across different organisms, as well as the mechanisms of their interaction with ligands and signal transduction should also receive attention.

Considering the various cell features that are regulated by TezRs, we hypothesize that their specific functions stem from their physical characteristics, such as length and presence of specific loops or nucleic acids conformations (121,122). A better understanding of these properties could lead to further and more accurate sub-classification of TezRs.

In follow-up studies, it will be critical to pay attention to the association of primary and secondary TezRs with the cell surface, and the way signals from these TezRs are transmitted further downstream in the cells. Based on our data, we speculate that secondary TezRs may exist as free receptors not bound to cell structures. However, we could not determine how TezRs interacted with protein receptors performing the same function. One can assume that some TezRs might be an integral, sensing (i.e. ligand-binding) part of such a protein receptors.

Moreover, given the recently discovered ability of DNA molecules to modify and misfold proteins, it is intriguing whether TezRs could possess a similar chaperoning function (27,123–125).

We are only starting to understand the sensory, receptive, and regulatory roles, as well as the structure of TezRs. Nevertheless, the need to deepen our knowledge in this field does not diminish the importance of the present observations. Finally, we believe that upcoming studies will expand our understanding of the whole set of sensing and regulatory processes involving TezRs.

## Conclusion

In this study, we describe for the first time the most external bacterial receptive and regulatory system, which enables sensing and response to numerous chemical (including xenobiotics), physical, and biological stimuli. This system consists of DNA- or RNA-based receptors, which we termed TezRs and classified based on the type of nucleic acids and localization. Besides TezRs, the system includes also reverse transcriptases and integrases.

Through removal of different TezRs, it is possible to modulate the cells’ responses to external stimuli, as well as that of known receptor-mediated signaling pathways. Importantly, loss of TezRs can cause unexpected activity and rapid changes to cell properties; we termed these cells “drunk cells”.

We characterized also the role of TezRs in cell memory formation and maintenance. Importantly, by affecting the TRB-receptor system, it is possible to erase the memory of previous events, leading to “zero cells”, whose existence opens new possibilities for regulating bacterial cells and populations.

In summary, the discovered TRB-receptor system enables the regulation of diverse cellular processes, including those whose modulation was previously poorly explored. Crucially, it also enables bacteria to survive in the face of constant changes to surrounding environmental factors.

## MATERIALS AND METHODS

### Bacterial and phage strains and culture conditions

*Bacillus pumilus* VT1200, *Staphylococcus аureus* MSSA VT209, *Staphylococcus aureus* SA58-1, *Pseudomonas putida* VT085, and *Escherichia coli* LE392 infected with bacteriophage λLZ1 [gpD-GFP b::ampR, kanR] bearing ampicillin and kanamycin resistance were obtained from a private collection (provided by Dr. V. TRB). *Escherichia coli* ATCC 25922 was purchased from the American Type Culture Collection (Manassas, VA, USA). Bacterial strains were passaged weekly on Columbia agar (BD Biosciences, Franklin Lakes, NJ, USA) and stored at 4 °C. All subsequent liquid subcultures were derived from colonies isolated from these plates and were grown in Luria-Bertani (LB) broth (Oxoid, Hampshire, UK; Sigma-Aldrich, St Louis, MO, USA), Columbia broth (BD Biosciences) or nutrient broth (CM001; Oxoid), if not stated otherwise. Other liquid media included M9 Minimal Salts (Sigma-Aldrich). For experiments on sold media, bacteria were cultured on Columbia agar, nutrient agar (CM003; Oxoid), TGV agar (TGV-Dx, Human Microbiology Institute, New York, NY, USA), LB agar (Sigma-Aldrich), Aureus ChromoSelect Agar Base (Sigma-Aldrich), tryptic soy agar (Sigma-Aldrich), and egg-yolk agar (Hardy Diagnostics, Santa Maria, CA, USA). Sheep red blood cells were purchased from Innovative Research (Peary Court, MI, USA). All cultures were incubated aerobically at 37 °C in a Heracell 150i incubator (Thermo Scientific, Waltham, MA, USA) if not stated otherwise. For anaerobic growth experiments, *P. putida* VT085 was plated on agar and cultivated in AnaeroGen 2.5-L Sachets (Oxoid) placed inside a CO2 incubator (Sanyo, Kitanagoya, Aichi, Japan) at 37 °C for 24 h.

### Reagents

Bovine pancreatic DNase I with a specific activity of 2,200 Kunitz units/mg and RNase A (both Sigma-Aldrich) were used at concentrations of 1 to 100 µg/mL. Ampicillin, kanamycin, rifampicin, vancomycin, nevirapine, etravirine, raltegravir, lactose, povidone iodine and dexamethasone were obtained from Sigma-Aldrich.

### Classification and nomenclature of TezRs

We classified TezRs based on the structural features of their DNA- or RNA-containing domains, as well as association with the bacterial cell surface determined by the possibility of being washed into culture medium or matrix (Table 2).

**Table 2.** Classification of TezRs in bacteria.

| Name of the receptor | Description of the receptor |
| --- | --- |
| Primary TezRs |  |
| TezR_D1 | DNA-based receptors located outside the membrane; they participate in cell regulation and are stably associated with the cell surface. |
| TezR_R1 | RNA-based receptors located outside the membrane; they participate in cell regulation and are stably associated with the cell surface. |
| Secondary TezRs |  |
| TezR_D2 | DNA-based receptors located outside the membrane; they participate in cell regulation and can be easily washed out along with culture medium or matrix. |
| TezR_R2 | RNA-based receptors located outside the membrane; they participate in cell regulation and can be easily washed out along with culture medium or matrix. |

To describe bacteria with certain destroyed TezR, we marked them with the superscript letter “d” – meaning destroyed.

An example of *E.coli* with destroyed primary DNA formed TezR will be designated as “E. coli TezR_D1^d^”, where TezR stands for TRB receptor and is followed by an underscore, then a capital letter representing the type of nucleic acid (D for DNA), followed by an Arabic numeral representing that it is a primary receptor, and “d” superscript meaning that this receptor was destroyed. The same principle of naming is applicable for bacteria with other destroyed TezRs. Cells with multiple cycles of TezRs destruction and restoration were named “zero cells” and are designated by a superscript letter “z” placed after the letter “d”.

### Removal of TezRs

To remove primary TezRs, bacteria were harvested by centrifugation at 4000 rpm for 15 min (Microfuge 20R; Beckman Coulter, La Brea, CA, USA), the pellet was washed twice in phosphate-buffered saline (PBS, pH 7.2) (Sigma-Aldrich) or nutrient medium to an optical density at 600 nm (OD600) of 0.003 to 0.5. Bacteria were treated for 30 min at 37 °C with nucleases (DNase I or RNase A), if not stated otherwise, washed three times in PBS or broth with centrifugation at 4000 × *g* for 15 min after each wash, and resuspended in PBS or broth. Bacteria, whose TezRs were deleted or made non-functional, were marked with the superscript letter “d”.

To study secondary TezRs, 1.5% TGV agar was used. After autoclaving at 121 °C for 20 min, the agar was cooled down to 45 °C and DNase I or RNase A, or a mixture of the two, was added, mixed, and 20 mL of the solution was poured into 90-mm glass Petri dishes.

For biofilm formation assays, bacteria were separated from the extracellular matrix by washing three times in PBS or broth with centrifugation at 4000 × *g* for 15 min after each wash. Then, 25 µL of suspension containing 7.5 log10 cells was inoculated into the center of the prepared solid medium surface supplemented or not with nucleases and incubated at 37 °C for different times.

### Inactivation of TezRs

To inactivate primary TezRs, bacteria were harvested by centrifugation at 4000 x g for 15 min (Microfuge 20R; Beckman Coulter, La Brea, CA, USA). The pellet was washed twice in PBS, pH 7.2 (Sigma-Aldrich). Bacteria were treated with PI for 30 min at 37 °C. If not stated otherwise, the PI-treated cells were washed three times in PBS with subsequent centrifugation at 4000 × *g* for 15 min, and resuspended in PBS or nutrient medium.

### Growth curve

For growth rate determination at the various time points, stationary phase bacteria were washed from the extracellular matrix, treated with nucleases (10 µg/mL), and 5.5 log10 cells were inoculated into 4.0 mL Columbia broth. OD600 was measured on a NanoDrop OneC spectrophotometer (Thermo Scientific).

### Bacterial viability test

To evaluate bacterial viability, bacterial suspensions were serially diluted and 100 µL of the diluted suspension was spread onto agar plates. Plates were incubated at 37 °C overnight and colony forming units (CFU) were counted the next day.

### Biofilm morphology

To culture bacterial biofilms, we prepared glass Petri dishes containing TGV agar supplemented or not with 100 µg/mL DNase I or RNase A, or a mixture of the two. Then, 25 µL of a suspension containing 5.5 log10 cells was inoculated in the center of the agar and the dishes were incubated at 37 °C for different times. The biofilms were photographed with a digital camera (Canon 6; Canon, Tokyo, Japan) and analyzed with Fiji/ImageJ software (126).

### Fluorescence microscopy

Differential interference contrast (DIC) and fluorescence microscopy were used to confirm the destruction of primary TezRs with nucleases. Bacteria treated or not with nucleases were sampled at OD600 of 0.1, washed from the matrix, fixed in 4% paraformaldehyde/PBS (Sigma-Aldrich) for 15 min at room temperature, and stored at 4 °C until use. Bacteria were centrifuged at 14,000 × *g* and cell pellets were dispersed in 10 μL PBS, incubated with SYTOX Green at a final concentration of 2 μM, and mounted in Fluomount mounting medium. Cells were imaged using an EVOS FL Auto Imaging System (Thermo Scientific) equipped with a 60× or 100× objective and 2× digital zoom.

Membrane-impermeable SYTOX Green stained cell surface-bound DNA and RNA. A reduction of green fluorescence compared to the untreated control, enabled the visualization of alterations elicited by nuclease treatment. Dead cells with permeable membranes showed a higher level of green fluorescence and were discarded from the analysis. No post-acquisition processing was performed; only minor adjustments of brightness and contrast were applied equally to all images. ImageJ software was used to quantify the signal intensity per cell; at least five representative images (60× field) were analyzed for each case (127).

### Light microscopy-based methods

Samples were imaged on an Axios plus microscope (Carl Zeiss, Jena, Germany) equipped with an ApoPlan ×100/1.25 objective. Images were acquired using a Canon 6 digital camera. Cell size was determined by staining cell membranes with methylene blue or Gram staining (both Sigma-Aldrich) and quantification in Fiji/ImageJ software. Values were expressed in px2 (126).

### Assays of RNase internalization

The internalization of RNase A was visualized in *B. pumilus*. *B. pumilus* (5.5 log_10_ cells/ml) in PBS were incubated with fluorescein isothiocyanate (FITC) labeled RNase A at 37 °C for 15 or 60 minutes as previously described (128). Bacteria were washed three times with PBS to remove any unbound protein. After washing the bacteria is cultivated for 2h in LB broth, washed to remove residual media components, and placed on a microscope slide for visualization. Fluorescence was monitored using a fluorescence microscope (Axio Imager Z1, Carl Zeiss, Germany). To visualize the internalization of RNase A, the biofilms of *B. pumilus* incubated with 100 µg/mL fluorescein-labeled RNase A were obtained as described earlier. After 24 h of growth at 37 °C, bacteria were washed three times with PBS to remove unbound proteins, and placed on a microscope to monitor the fluorescence using a fluorescence microscope (Axio Imager Z1, Carl Zeiss, Germany).

### Generation of RNA sequencing data

To isolate RNA, the cell suspension obtained 2.5h post-nuclease treatment were washed thrice in PBS,pH 7.2 (Sigma) and centrifuged each time at 4000 × *g* for 15 min (Microfuge 20R, Beckman Coulter) followed by resuspension in PBS.

RNA was purified using RNeasy Mini Kit (Qiagen) according to the manufacturer’s protocol. The quantity and quality of RNA was spectrophotometrically evaluated by measuring the UV absorbance at 230/260/280 nm with the NanoDrop OneC spectrophotometer (ThermoFisher Scientific).

Transcriptome sequencing (RNA-Seq) libraries were prepared using an Illumina TruSeq Stranded Total RNA Library Prep kit. RNA was ribodepleted using the Epicenter Ribo-Zero magnetic gold kit (catalog no. RZE1224) according to the manufacturer’s guidelines. The libraries were pooled equimolarly and sequenced in an Illumina NextSeq 500 (Illumona, San Diego CA) platform with paired 150-nucleotide reads (130MM reads max).

### Analysis of RNA sequencing data

Sequencing reads were mapped corresponding to the reference genome of *S. aureus* NCTC 8325 (NCBI Reference Sequence: NC_007795), and expression levels were estimated using Geneious 11.1.5. Transcripts with an adjusted P value of < 0.05 and log_2_ fold change value of ± 0.5 were considered for significant differential expression. PCA, volcano plots and Euclidean distances plots were generated using the ggplot2 package in R, and the Venn diagram was obtained using BioVenn (129).

### Sporulation assay

Sporulation was analyzed under the microscope by counting cells and spores in 20 microscope fields and three replicates. For each image, we calculated the number of spores and the number of cells. Then, we plotted the ratio of spores to the combined number of cells and spores in each bin. Sporulation under stress conditions was carried out by heating the bacterial culture at 42 °C for 15 min.

### Modulation of thermotolerance

Overnight *S. aureus* VT209 cultured in LB broth supplemented or not with raltegravir (5 µg/mL) was separated from the extracellular matrix by washing in PBS and then diluted with PBS to OD600 of 0.5. Bacteria were left untreated or treated with nucleases to remove primary TezRs and 5.5 log10 CFU/mL were placed in 2-mL microcentrifuge tubes (Axygen Scientific Inc., Union City, CA, USA). Each tube was heated to 37, 40, 45, 50, 55, 60, 65, 70 or 75 °C in a dry bath (LSETM Digital Dry Bath; Corning, Corning, NY, USA) for 15 min. After heating, control *S. aureus* were immediately treated with nucleases to delete primary TezRs, washed three times to remove nucleases, serially diluted, plated on LB agar, and the number of CFU was determined within 24 h.

### Modulation of thermotolerance restoration after TezRs loss

To determine the time it took for thermotolerance to be restored in bacteria following TezRs removal, overnight *S. aureus* VT209 cultures were treated with 10 µg/mL DNase I or RNase A, or a mixture of the two. Bacteria lacking TezRs were inoculated in LB broth and sampled hourly for up to 8 h. The samples were heated at the maximum temperature tolerated by the bacteria and viability was assessed as described in the previous section. Untreated *S. aureus* were used as a control and were processed the same way by heating at the lowest non-tolerable temperature, serially diluted, plated on LB agar, and assessed for CFUs within 24 h. Complete restoration of normal temperature tolerance coincided with growth inhibition at higher temperatures. The experiment was not extended beyond this time point.

### Bacteriophage infection assay

An overnight *E. coli* LE392 culture was diluted 1:1000 and grown in liquid LB broth supplemented with 0.2% maltose and 10 mM MgSO4 at 30 °C for 18–24 h, until OD600 of 0.4. Cells were separated from the extracellular matrix by three washes in PBS and centrifugation at 4000 × *g* for 15 min and 20 °C after each wash, followed by resuspension in ice-cold LB broth supplemented with 10 mM MgSO4 to OD600 of 1.0. Approximately 10 μL of plaque-forming units of the purified λ phage was added to 200 μL *E. coli* LE392 with intact TezRs. The suspension was incubated for 30 min on ice and another 90 min at room temperature to ensure that the phage genome entered the cells (Single-Cell Studies of Phage λ: Hidden Treasures Under Occam’s Rug). The remaining phages were removed by three washes in PBS and centrifugation at 4000 x g for 15 min and 20 °C after each wash.

Bacteria were treated with nucleases to destroy primary TezRs, followed by three centrifugation steps at 4000 x g for 15 min and 20 °C. Control *E. coli* were not treated with nucleases. After that, 100 μL of bacterial suspension was plated as a lawn on LB agar supplemented with 10 μg/mL kanamycin and 100 μg/mL ampicillin, incubated for 24 h at 30 °C, and the number of Amp/Kanr colonies was determined.

### Persister assay

*E. coli* ATCC 25922 were treated with nucleases to remove primary TezRs, inoculated in LB broth supplemented with ampicillin (150 μg/mL), and incubated at 37 °C for 6 h. Samples taken before and after incubation with ampicillin were plated on LB agar without antibiotics to determine the CFU (^85^). The frequency of persisters was calculated as the number of persisters in a sample relative to the number of cells before antibiotic treatment in each probe.

### Analysis of virulence factors production

*S. aureus* SA58-1 were treated with nucleases to remove primary TezRs and resuspended in PBS to 6.0 log10 CFU/mL.

The hemolytic test was performed as previously described with minor modifications (130). Briefly, bacterial cells were plated in the center of Columbia agar plates supplemented with 5% sheep red blood cells and incubated at 37 °C for 24 h. A greenish zone around the colony denoted α-hemolysin activity; whereas β-hemolysin (positive) and γ-hemolysin (negative) activities were indicated by the presence or absence of a clear zone around the colonies. The size of the hemolysis zone (in mm) was measured.

Lecithinase activity by bacteria with intact TezRs or lacking TezRs was determined by plating cells on egg-yolk agar and incubation at 37 °C for 48 h. The presence of the precipitation zone and its diameter were evaluated (131).

### UV assay

*S. aureus* VT209 were treated with nucleases to remove primary TezRs. Control probes were left untreated. Bacteria at 8.5 log10 CFU/mL in PBS were added to 9-cm Petri dishes, placed under a light holder equipped with a new 254-nm UV light tube (TUV 30W/G30T8; Philips, Amsterdam, The Netherlands), and irradiated for different times at a distance of 50 cm. After treatment, bacteria were serially diluted, plated on nutrition agar plates, incubated for 24 h, and CFU were determined.

### Animal models

All animal procedures and protocols were approved by the institutional animal care and use (IACUC) committee at the Human Microbiology Institute (protocol: # T-19-204) and all efforts were made to minimize animal discomfort and suffering. Adult C57BL/6 mice weighing from 18 to 20 g (Jackson Laboratories, Bar Harbor, ME, USA) were fed ad libitum and housed in individual cages in a facility free of known murine pathogens. Animals were cared for in accordance with National Research Council recommendations, and experiments were carried out in accordance with the Guide for the Care and Use of Laboratory Animals (132).

Animals were randomly designated to four groups of eight mice each, which were used to measure the load of *S. aureus* SA58-1. Mice were anesthetized with 2% isoflurane, and intraperitoneally injected with nuclease-treated *S. aureus* at 10.1 log10 to 10.2 log10 CFU/mouse. Control animals received untreated *S. aureus* SA58-1. After 12 h, mice were euthanized by CO2 and cervical dislocation, and the bacterial load in the peritoneum, liver, spleen, and kidneys was determined by serial dilution and CFU counts after 48 h of culture on plates with selective *S. aureus* agar. Cell morphology was determined under an Axios plus microscope, following staining with a Gram stain kit (Merck, Darmstadt, Germany).

### Magnetic exposure conditions

The effect of the TRB-receptor system on regulation of *B. pumilus* VT1200 growth when exposed to regular magnetic and shielded geomagnetic fields was assessed. *B. pumilus* lacking primary and secondary TezRs were obtained as previously discussed. Final inoculi of 5.5 log10 CFU/mL in 25 µL were dropped in the center of agar-filled Petri dishes. Magnetic exposure conditions were modulated by placing the Petri dish in a custom-made box made of five layers of 10-µm-thick μ metal (to shield geomagnetic field) at 37 °C for 24 h. Biofilm surface coverage was analyzed using Fiji/ImageJ software and expressed as px2 (127,133).

In a second experimental, *B. pumilus* VT1200 with intact TezRs and missing TezR_R1 were exposed to regular magnetic conditions or a shielded geomagnetic field as described above, and colony morphology was analyzed after 8 and 24 h. Images of the plates were acquired using a Canon 6 digital camera.

### Estimation of spontaneous mutation rates

To calculate the number of mutation events, we used *E. coli* ATCC 25922, treated with nucleases to remove primary TezRs or untreated controls, and standardized at 9.0 log10 cells. The number of spontaneous mutations to RifR was used to estimate the mutation rate. This was determined by counting the number of colonies formed on Mueller-Hinton agar supplemented or not with rifampicin (100 µg/mL). After incubation at 37 °C for 48 h, CFU as well as rifampicin resistant mutants were counted and the mutation rate was calculated by the Jones median estimator method (134).

### Light exposure experiments

*B. pumilus* VT1200 lacking primary and secondary TezRs were obtained as described previously. An aliquot containing 5.5 log10 bacteria in 25 µL was placed in the center of Columbia agar plates, which were then incubated at 37 °C for 7 or 24 h while irradiated with halogen lamps of 150 W (840 lm) (Philips, Shanghai, China). Colonies were photographed with a Canon 6 digital camera. The distance between the light source and the sample was 20 cm. Control probes were processed the same way, but were grown in the dark.

### Chemotaxis and dispersal measurements

The assay was performed as described previously with some modifications. Briefly, assay plates containing TGV agar were prepared by adding 250 µL fresh human plasma to a sector comprising 1/6 of the plate. The plasma was filtered through a 0.22-μm pore-size filter (Millipore Corp., Bedford, MA, USA) immediately prior to use. Written informed consent was obtained from all patients to use their blood samples for research purposes, and the study was approved by the institutional review board of the Human Microbiology Institute (# VB-021420).

*B. pumilus* VT1200 devoid of primary and secondary TezRs were obtained as described previously. An aliquot containing 5.5 log10 cells in 25 µL was placed in the center of the plates, which were then incubated at 37 °C for 24 h and photographed with a Canon 6 digital camera. Chemotaxis was evaluated by measuring the migration of the central colony towards the plate sector containing plasma. Colony dispersal was assessed based on the appearance of small colonies on the agar surface.

### Effect of reverse transcriptase inhibitors and integrase on bacterial growth

Minimum inhibitory concentrations (MICs) of nevirapine and etravirine against *S. aureus* VT209 were evaluated. *S. aureus* VT209 with intact or missing primary TezRs were obtained as described previously. Bacteria were incubated in LB broth supplemented or not with nevirapine (5 µg/mL) or etravirine (5 µg/mL). These values corresponded to > 1/100 their MICs. Growth was monitored by measuring OD600 during the first 6 h of incubation at 37 °С and recorded at hourly intervals on a NanoDrop OneC spectrophotometer.

### Biochemical analysis

Biochemical tests were carried out using the colorimetric reagent cards GN (gram-negative) and BCL (gram-positive spore-forming bacilli) of the VITEK® 2 Compact 30 system (BioMérieux, Marcy l’Étoile, France) according to the manufacturer’s instructions. The generated data were analyzed using VITEK® 2 software version 7.01, according to the manufacturer’s instructions.

### Recognition of lactose and dexamethasone

The role of the TRB-receptor system in the recognition of lactose and dexamethasone was investigated with *E. coli* ATCC 25922 and *B. pumilus* VT1200. Bacterial suspensions of control bacteria and those lacking primary TezRs were adjusted to a common CFU value and incubated in fresh M9 medium supplemented or not with 146 mM lactose or 127 mM dexamethasone.

The lag phase, representing the period between inoculation of bacteria and the start of biomass growth, was measured by monitoring OD600. The lag phase reflects the time required for the onset of nutrient utilization (64,65).

### Cell memory formation experiments

The onset o bacterial memory was defined as the time required for dexamethasone to start being consumed (time lag) in dexamethasone-naïve and dexamethasone-sentient *B. pumilus* VT1200 and *E. coli* ATCC 25922. To study the first exposure to dexamethasone, *B. pumilus* or *E. coli* with intact TezRs were incubated in fresh M9 medium supplemented or not with 127 mM dexamethasone for 24 h. To study the second exposure to dexamethasone, bacteria were taken after 24 h of cultivation from the first exposure to dexamethasone and washed three times in PBS with centrifugation at 4000 x g for 15 min and 20°C after each wash. Bacteria were adjusted to a common OD600 and incubated again in fresh M9 medium supplemented with dexamethasone. During the first and second exposures to dexamethasone, samples were taken at hourly intervals for the first 6 h and OD600 was measured with a NanoDrop OneC spectrophotometer to determine the lag phase. The different time lag between the first and second exposures to dexamethasone represented the formation of memory (135).

### Evaluation of the role of TezRs in memory formation

To study the role of TezR_R1 in remembering previous exposures to nutrients, we assessed the difference in the time required for TezR_R1 of dexamethasone-naïve and dexamethasone-sentient *E. coli* ATCC 25922 to sense and trigger dexamethasone utilization. The two *E. coli* cell types with intact TezRs were pretreated with 127 mM dexamethasone for 5, 10, 15 or 20 min. Next, bacteria were treated with RNase A to remove TezR_R1, and inoculated in fresh M9 medium supplemented with dexamethasone. The lag phase prior to dexamethasone consumption was determined by monitoring OD600 every hour.

### Memory loss experiments

THE role of TezRs in bacterial memory loss was studied by comparing the lag phase of dexamethasone-naïve and dexamethasone-sentient *B. pumilus* VT1200 with intact TezRs (14). Bacteria were cultivated in M9 medium supplemented with 127 mM dexamethasone for 24 h, centrifuged at 4000 x g for 15 min, and washed in M9 medium without dexamethasone. The cells then underwent repeated rounds of TezR_R1 removal and restoration, followed by growth in M9 broth without dexamethasone. After 24 h of cultivation at 37 °C, bacteria were isolated from the medium, TezR_R1 were removed again, and bacteria were re-inoculated in fresh M9 broth. In total, cultivation in M9 broth followed by TezR_R1 removal was repeated three times. Samples were taken prior to every TezR_R1 removal step, bacteria were washed, inoculated in M9 broth supplemented with dexamethasone, and the time lag to dexamethasone consumption was assessed by monitoring OD600.

After the third set of cultivation in M9 broth, bacteria were centrifuged and inoculated in fresh M9 broth. They were then cultivated for 24 h, centrifuged, washed, and inoculated in M9 broth supplemented with dexamethasone to mimic a second contact with dexamethasone. The time lag to dexamethasone consumption was assessed by monitoring OD600. Bacteria from the control group were processed the same way, but without undergoing TezR_R1 removal.

### Raltegravir in cell memory formation experiments

The MIC of raltegravir against *S. aureus* VT209 was evaluated. To determine the effect of raltegravir on bacterial memory, *B. pumilus* VT1200 were grown on fresh M9 medium supplemented or not with 127 mM dexamethasone, with or without additionally supplementation with raltegravir (5 µg/mL, a 100-times lower concentration than the MIC). The biochemical profile of cells was analyzed with a VITEK® 2.

To evaluate the maximal time required for raltegravir to affect dexamethasone utilization, *B. pumilus* VT1200 were grown in M9 broth supplemented with 127 mM dexamethasone, while raltegravir was added at 0 h, 15 min, 30 min, 1 h or 2 h. The samples were taken at hourly intervals for the first 6 h to measure OD600 and determine the lag phase.

### Statistics

At least three biological replicates were performed for each experimental condition unless stated otherwise. Each data point was denoted by the mean value ± standard deviation (SD). A two-tailed *t*-test was performed for pairwise comparisons and p ≤ 0.05 was considered significant. Bacterial quantification data were log10-transformed prior to analysis. Statistical analyses for the biofilm assays and hemolysin test were performed using Student’s *t*-test. Data from animal and sporulation studies were calculated using a two-tailed Mann-Whitney U test. GraphPad Prism version 9 (GraphPad Software, San Diego, CA, USA) or Excel 10 (Microsoft, Redmond, WA, USA) were applied for statistical analysis and illustration.

---

Supplementary Table 1. Effect of primary TezR_D1/R1 removal on bacterial size.

Supplementary Table 2. Effect of primary TezRs removal on the size of *B. pumilus* VT1200 biofilm.

Supplementary Table 3. Effect of TezR removal on sporulation under normal conditions.

Supplementary Table 4. Effect of TezR removal on sporulation under stress conditions.

Supplementary Table 5. MICs of tested reverse transcriptase inhibitors and integrase inhibitor against control *S. aureus*.

Supplementary Figure S1. Absence of RNase A internalization in *B. pumilus*.

Supplementary Figure S2. Effect of TezRs removal on light sensing.

## Author Contributions

VT and GT designed experiments. VT and GT supervised data analysis, analyzed data and wrote the manuscript.

## ACKNOWLEDGMENTS

We gratefully acknowledge Dr. You Zhou, Microscopy facility at the Center for Biotechnology in University of Nebraska-Lincoln for help in microscopy; Kristina Kardava, Marya Vecherkovskaya and Tatiana Lazareva for setting some experiments; Genome Technology Center (GTC) for expert library preparation and sequencing, and the Applied Bioinformatics Laboratories (ABL) for providing bioinformatics support and helping with the analysis and interpretation of the data. GTC and ABL are shared resources partially supported by the Cancer Center Support Grant P30CA016087 at the Laura and Isaac Perlmutter Cancer Center. This work has used computing resources at the NYU School of Medicine High Performance Computing (HPC) Facility.

## References

1. Wadhams, G. H. & Armitage, J. P. Making sense of it all: bacterial chemotaxis. Nat. Rev. Mol. Cell Biol. 5, (2004).

2. Ortega, Á., Zhulin, I. B. & Krell, T. Sensory Repertoire of Bacterial Chemoreceptors. Microbiol. Mol. Biol. Rev. 81, (2017).

3. Bi, S., Jin, F. & Sourjik, V. Inverted signaling by bacterial chemotaxis receptors. Nat. Commun. 9, (2018).

4. Rayo, J., Amara, N., Krief, P. & Meijler, M. M. Live Cell Labeling of Native Intracellular Bacterial Receptors Using Aniline-Catalyzed Oxime Ligation. J. Am. Chem. Soc. 133, 7469–7475 (2011).

5. Falke, J. J. & Hazelbauer, G. L. Transmembrane signaling in bacterial chemoreceptors. Trends Biochem. Sci. 26, (2001).

6. Ng, W.-L. et al. Probing bacterial transmembrane histidine kinase receptor-ligand interactions with natural and synthetic molecules. Proc. Natl. Acad. Sci. 107, (2010).

7. Falke, J. J. Cooperativity between bacterial chemotaxis receptors. Proc. Natl. Acad. Sci. 99, (2002).

8. Hazelbauer, G. L., Falke, J. J. & Parkinson, J. S. Bacterial chemoreceptors: high-performance signaling in networked arrays. Trends Biochem. Sci. 33, (2008).

9. Yang, Y. & Sourjik, V. Opposite responses by different chemoreceptors set a tunable preference point in *Escherichia coli* pH taxis. Mol. Microbiol. 86, (2012).

10. Machuca, M. A. et al. Helicobacter pylori chemoreceptor TlpC mediates chemotaxis to lactate. Sci. Rep. 7, (2017).

11. Li, H. & Wang, H. Activation of xenobiotic receptors: driving into the nucleus. Expert Opin. Drug Metab. Toxicol. 6, (2010).

12. Sourjik, V. & Berg, H. C. Functional interactions between receptors in bacterial chemotaxis. Nature 428, (2004).

13. Jacquin, J. et al. Microbial Ecotoxicology of Marine Plastic Debris: A Review on Colonization and Biodegradation by the “Plastisphere”. Front. Microbiol. 10, (2019).

14. Wolf, D. M. et al. Memory in Microbes: Quantifying History-Dependent Behavior in a Bacterium. PLoS One 3, (2008).

15. Kordes, A. et al. Establishment of an induced memory response in Pseudomonas aeruginosa during infection of a eukaryotic host. ISME J. 13, (2019).

16. Gosztolai, A. & Barahona, M. Cellular memory enhances bacterial chemotactic navigation in rugged environments. Commun. Phys. 3, (2020).

17. Andersson, S. G. E. Stress management strategies in single bacterial cells. Proc. Natl. Acad. Sci. 113, (2016).

18. Lambert, G. & Kussell, E. Memory and Fitness Optimization of Bacteria under Fluctuating Environments. PLoS Genet. 10, (2014).

19. Stock, J. B. & Zhang, S. The biochemistry of memory. Curr. Biol. 23, (2013).

20. Vashistha, H., Kohram, M. & Salman, H. Non-genetic inheritance restraint of cell-to-cell variation. Elife 10, (2021).

21. Yang, C.-Y. et al. Encoding Membrane-Potential-Based Memory within a Microbial Community. Cell Syst. 10, (2020).

22. . Matsunaga, T., et al. Complete Genome Sequence of the Facultative Anaerobic Magnetotactic Bacterium Magnetospirillum sp. strain AMB-1. DNA Res. 12, (2005).

23. McCausland, H. C. & Komeili, A. Magnetic genes: Studying the genetics of biomineralization in magnetotactic bacteria. PLOS Genet. 16, (2020).

24. Scheffel, A. et al. An acidic protein aligns magnetosomes along a filamentous structure in magnetotactic bacteria. Nature 440, (2006).

25. Nordmann, G. C., Hochstoeger, T. & Keays, D. A. Magnetoreception—A sense without a receptor. PLOS Biol. 15, (2017).

26. Monteil, C. L. & Lefevre, C. T. Magnetoreception in Microorganisms. Trends Microbiol. 28, (2020).

27. Blank, M. & Goodman, R. DNA is a fractal antenna in electromagnetic fields. Int. J. Radiat. Biol. 87, (2011).

28. Berashevich, J. & Chakraborty, T. How the Surrounding Water Changes the Electronic and Magnetic Properties of DNA. J. Phys. Chem. B 112, (2008).

29. Nikiforov, V. N., Koksharov, Y. A. & Irkhin, V. Y. Magnetic properties of “doped” DNA. J. Magn. Magn. Mater. 459, (2018).

30. Yoney, A. & Salman, H. Precision and Variability in Bacterial Temperature Sensing. Biophys. J. 108, (2015).

31. Chursov, A., Kopetzky, S. J., Bocharov, G., Frishman, D. & Shneider, A. RNAtips: analysis of temperature-induced changes of RNA secondary structure. Nucleic Acids Res. 41, (2013).

32. Sengupta, P. & Garrity, P. Sensing temperature. Curr. Biol. 23, (2013).

33. Barria, C., Malecki, M. & Arraiano, C. M. Bacterial adaptation to cold. Microbiology 159, (2013).

34. Abatedaga, I. et al. Integration of Temperature and Blue-Light Sensing in Acinetobacter baumannii Through the BlsA Sensor. Photochem. Photobiol. 93, 805–814 (2017).

35. Golic, A. E. et al. BlsA Is a Low to Moderate Temperature Blue Light Photoreceptor in the Human Pathogen Acinetobacter baumannii. Front. Microbiol. 10, (2019).

36. Briegel, A. et al. New Insights into Bacterial Chemoreceptor Array Structure and Assembly from Electron Cryotomography. Biochemistry 53, (2014).

37. Bi, S. & Sourjik, V. Stimulus sensing and signal processing in bacterial chemotaxis. Curr. Opin. Microbiol. 45, (2018).

38. Parkinson, J. S., Hazelbauer, G. L. & Falke, J. J. Signaling and sensory adaptation in Escherichia coli chemoreceptors: 2015 update. Trends Microbiol. 23, (2015).

39. Beyer, Szöllössi, Byles, Fischer & Armitage. Mechanism of Signalling and Adaptation through the Rhodobacter sphaeroides Cytoplasmic Chemoreceptor Cluster. Int. J. Mol. Sci. 20, (2019).

40. Irigoyen, J. P., Muñoz-Cánoves, P., Montero, L., Koziczak, M. & Nagamine, Y. The plasminogen activator system: biology and regulation. Cell. Mol. Life Sci. 56, (1999).

41. Bohn, C. et al. Experimental discovery of small RNAs in Staphylococcus aureus reveals a riboregulator of central metabolism. Nucleic Acids Res. 38, 6620–6636 (2010).

42. Kengmo Tchoupa, A., et al. The type VII secretion system protects Staphylococcus aureus against antimicrobial host fatty acids. Sci. Rep. 10, 14838 (2020).

43. Taylor, J. C. et al. A type VII secretion system of Streptococcus gallolyticus subsp. gallolyticus contributes to gut colonization and the development of colon tumors. PLOS Pathog. 17, e1009182 (2021).

44. Whitchurch CB, T.-N. T. R. P. M. J. Extracellular DNA required for bacterial biofilm formation. Science (80-. ). 295, 1487 (2022).

45. Ingham, C. J. & Jacob, E. Swarming and complex pattern formation in Paenibacillus vortex studied by imaging and tracking cells. BMC Microbiol. 8, (2008).

46. Kampers, L. F. C. et al. A metabolic and physiological design study of Pseudomonas putida KT2440 capable of anaerobic respiration. BMC Microbiol. 21, (2021).

47. Glasser, N. R., Kern, S. E. & Newman, D. K. Phenazine redox cycling enhances anaerobic survival in *P seudomonas aeruginosa* by facilitating generation of ATP and a proton-motive force. Mol. Microbiol. 92, (2014).

48. Nikel, P. I. & de Lorenzo, V. Engineering an anaerobic metabolic regime in Pseudomonas putida KT2440 for the anoxic biodegradation of 1,3-dichloroprop-1-ene. Metab. Eng. 15, (2013).

49. Eschbach, M. et al. Long-Term Anaerobic Survival of the Opportunistic Pathogen *Pseudomonas aeruginosa* via Pyruvate Fermentation. J. Bacteriol. 186, (2004).

50. Fuchs, S., Pané-Farré, J., Kohler, C., Hecker, M. & Engelmann, S. Anaerobic Gene Expression in *Staphylococcus aureus*. J. Bacteriol. 189, (2007).

51. Kadowaki, T. et al. Porphyromonas gingivalis Proteinases as Virulence Determinants in Progression of Periodontal Diseases. J. Biochem. 128, (2000).

52. Saunders, S. H. et al. Extracellular DNA Promotes Efficient Extracellular Electron Transfer by Pyocyanin in Pseudomonas aeruginosa Biofilms. Cell 182, (2020).

53. Ciemniecki, J. A. & Newman, D. K. The Potential for Redox-Active Metabolites To Enhance or Unlock Anaerobic Survival Metabolisms in Aerobes. J. Bacteriol. 202, (2020).

54. Rashid, M. H. & Kornberg, A. Inorganic polyphosphate is needed for swimming, swarming, and twitching motilities of Pseudomonas aeruginosa. Proc. Natl. Acad. Sci. 97, (2000).

55. Fraser, G. M. & Hughes, C. Swarming motility. Curr. Opin. Microbiol. 2, (1999).

56. Hagai, E. et al. Surface-motility induction, attraction and hitchhiking between bacterial species promote dispersal on solid surfaces. ISME J. 8, (2014).

57. Abee, T., Kovács, Á. T., Kuipers, O. P. & van der Veen, S. Biofilm formation and dispersal in Gram-positive bacteria. Curr. Opin. Biotechnol. 22, (2011).

58. Bartolini, M. et al. Regulation of Biofilm Aging and Dispersal in *Bacillus subtilis* by the Alternative Sigma Factor SigB. J. Bacteriol. 201, (2019).

59. McDougald, D., Rice, S. A., Barraud, N., Steinberg, P. D. & Kjelleberg, S. Should we stay or should we go: mechanisms and ecological consequences for biofilm dispersal. Nat. Rev. Microbiol. 10, (2012).

60. Velasco, E. et al. A new role for Zinc limitation in bacterial pathogenicity: modulation of α-hemolysin from uropathogenic Escherichia coli. Sci. Rep. 8, (2018).

61. Golding, I. Single-Cell Studies of Phage λ: Hidden Treasures Under Occam’s Rug. Annu. Rev. Virol. 3, (2016).

62. Huang, Y. J. and B. L.. C32 LUNG INJURY, ARDS, AND SEPSIS: The Effects Of inhaled Glucocorticoids On Growth Of Pseudomonas AerugINOSa. Am. J. Respir. Crit. Care Med. 195 (2017).

63. DeNiro, M. & Epstein, S. Mechanism of carbon isotope fractionation associated with lipid synthesis. Science (80-. ). 197, (1977).

64. Paliy, O. & Gunasekera, T. S. Growth of E. coli BL21 in minimal media with different gluconeogenic carbon sources and salt contents. Appl. Microbiol. Biotechnol. 73, (2007).

65. Fernández de las Heras, L., García Fernández, E., María Navarro Llorens, J., Perera, J. & Drzyzga, O. Morphological, Physiological, and Molecular Characterization of a Newly Isolated Steroid-Degrading Actinomycete, Identified as Rhodococcus ruber Strain Chol-4. Curr. Microbiol. 59, (2009).

66. Basan, M. et al. A universal trade-off between growth and lag in fluctuating environments. Nature 584, (2020).

67. Shibasaki, H., Tanabe, C., Furuta, T. & Kasuya, Y. Hydrolysis of conjugated steroids by the combined use of β-glucuronidase preparations from helix pomatia and ampullaria: determination of urinary cortisol and its metabolites. Steroids 66, (2001).

68. Zarkan, A. et al. Indole Pulse Signalling Regulates the Cytoplasmic pH of E. coli in a Memory-Like Manner. Sci. Rep. 9, (2019).

69. Lyon, P. The cognitive cell: bacterial behavior reconsidered. Front. Microbiol. 6, (2015).

70. García-López, V. et al. Molecular machines open cell membranes. Nature 548, 567–572 (2017).

71. Szilvay, A. M., Stern, B., Blichenberg, A. & Helland, D. E. Structural and functional similarities between HIV-1 reverse transcriptase and the *Escherichia coli* RNA polymerase β′ subunit. FEBS Lett. 484, (2000).

72. Sciamanna, I., De Luca, C. & Spadafora, C. The Reverse Transcriptase Encoded by LINE-1 Retrotransposons in the Genesis, Progression, and Therapy of Cancer. Front. Chem. 4, (2016).

73. Spanopoulou, E. et al. The Homeodomain Region of Rag-1 Reveals the Parallel Mechanisms of Bacterial and V(D)J Recombination. Cell 87, (1996).

74. Nishana, M., Nilavar, N. M., Kumari, R., Pandey, M. & Raghavan, S. C. HIV integrase inhibitor, Elvitegravir, impairs RAG functions and inhibits V(D)J recombination. Cell Death Dis. 8, (2017).

75. Hershko, A. & Ciechanover, A. THE UBIQUITIN SYSTEM. Annu. Rev. Biochem. 67, (1998).

76. MartÃ-nez, L. C. & Vadyvaloo, V. Mechanisms of post-transcriptional gene regulation in bacterial biofilms. Front. Cell. Infect. Microbiol. 4, (2014).

77. Kearns, D. B. A field guide to bacterial swarming motility. Nat. Rev. Microbiol. 8, (2010).

78. Claessen, D., Rozen, D. E., Kuipers, O. P., Søgaard-Andersen, L. & van Wezel, G. P. Bacterial solutions to multicellularity: a tale of biofilms, filaments and fruiting bodies. Nat. Rev. Microbiol. 12, (2014).

79. Kaplan, J. B. Biofilm Dispersal: Mechanisms, Clinical Implications, and Potential Therapeutic Uses. J. Dent. Res. 89, (2010).

80. Güvener, Z. T. & Harwood, C. S. Subcellular location characteristics of the Pseudomonas aeruginosa GGDEF protein, WspR, indicate that it produces cyclic-di-GMP in response to growth on surfaces. Mol. Microbiol. 071119190133004-??? (2007) doi:10.1111/j.1365-2958.2007.06008.x.

81. Tetz, G. & Tetz, V. Introducing the sporobiota and sporobiome. Gut Pathog. 9, 38 (2017).

82. Errington, J. Regulation of endospore formation in Bacillus subtilis. Nat. Rev. Microbiol. 1, 117–126 (2003).

83. Dubnau, D. & Losick, R. Bistability in bacteria. Mol. Microbiol. 61, 564–572 (2006).

84. Wilmaerts, D., Windels, E. M., Verstraeten, N. & Michiels, J. General Mechanisms Leading to Persister Formation and Awakening. Trends Genet. 35, 401–411 (2019).

85. Svenningsen, M. S., Veress, A., Harms, A., Mitarai, N. & Semsey, S. Birth and Resuscitation of (p)ppGpp Induced Antibiotic Tolerant Persister Cells. Sci. Rep. 9, 6056 (2019).

86. Loh, E., Righetti, F., Eichner, H., Twittenhoff, C. & Narberhaus, F. RNA Thermometers in Bacterial Pathogens. Microbiol. Spectr. 6, (2018).

87. Krishna, S., Maslov, S. & Sneppen, K. UV-Induced Mutagenesis in Escherichia coli SOS Response: A Quantitative Model. PLoS Comput. Biol. 3, e41 (2007).

88. Wadhawan, S. & Gautam, S. Rescue of Escherichia coli cells from UV-induced death and filamentation by caspase-3 inhibitor. Int. Microbiol. 22, 369–376 (2019).

89. Erental, A., Kalderon, Z., Saada, A., Smith, Y. & Engelberg-Kulka, H. Apoptosis-Like Death, an Extreme SOS Response in Escherichia coli. MBio 5, (2014).

90. Irkhin, V. Y. & Nikiforov, V. N. Quantum effects and magnetism in the spatially distributed DNA molecules. J. Magn. Magn. Mater. 459, 345–349 (2018).

91. Savelyev, I.V., Zyryanova, N.V., Polesskaya, O.O. and Myakishev-Rempel, M., 2019. On the existence of the DNA resonance code and its possible mechanistic connection to the neural code. NeuroQuantology, 17(2), P. 56. & Savelyev IV, Zyryanova NV, Polesskaya OO, M.-R. M. On The Existence of The DNA Resonance Code and Its Possible Mechanistic Connection to The Neural Code. NeuroQuantology 17, 56 (2019).

92. Yi, J. Emergent paramagnetism of DNA molecules. Phys. Rev. B 74, 212406 (2006).

93. Montagnier, L., Aïssa, J., Ferris, S., Montagnier, J.-L. & Lavalléee, C. Electromagnetic signals are produced by aqueous nanostructures derived from bacterial DNA sequences. Interdiscip. Sci. Comput. Life Sci. 1, (2009).

94. Zhang, Q., Throolin, R., Pitt, S. W., Serganov, A. & Al-Hashimi, H. M. Probing Motions between Equivalent RNA Domains Using Magnetic Field Induced Residual Dipolar Couplings: Accounting for Correlations between Motions and Alignment. J. Am. Chem. Soc. 125, 10530–10531 (2003).

95. Briegel, A. et al. Structure of bacterial cytoplasmic chemoreceptor arrays and implications for chemotactic signaling. Elife 3, (2014).

96. Schaaper, R. M. & Dunn, R. L. Spectra of spontaneous mutations in Escherichia coli strains defective in mismatch correction: the nature of in vivo DNA replication errors. Proc. Natl. Acad. Sci. 84, (1987).

97. Canchaya, C., Fournous, G., Chibani-Chennoufi, S., Dillmann, M.-L. & Brüssow, H. Phage as agents of lateral gene transfer. Curr. Opin. Microbiol. 6, 417–424 (2003).

98. Yang, D. et al. Human Ribonuclease A Superfamily Members, Eosinophil-Derived Neurotoxin and Pancreatic Ribonuclease, Induce Dendritic Cell Maturation and Activation. J. Immunol. 173, 6134–6142 (2004).

99. Sumby, P. et al. Extracellular deoxyribonuclease made by group A Streptococcus assists pathogenesis by enhancing evasion of the innate immune response. Proc. Natl. Acad. Sci. 102, (2005).

100. Lux, R., Jahreis, K., Bettenbrock, K., Parkinson, J. S. & Lengeler, J. W. Coupling the phosphotransferase system and the methyl-accepting chemotaxis protein-dependent chemotaxis signaling pathways of Escherichia coli. Proc. Natl. Acad. Sci. 92, (1995).

101. Sheth, R. U. & Wang, H. H. DNA-based memory devices for recording cellular events. Nat. Rev. Genet. 19, (2018).

102. Kurosaki, T., Kometani, K. & Ise, W. Memory B cells. Nat. Rev. Immunol. 15, 149–159 (2015).

103. McHeyzer-Williams, M., Okitsu, S., Wang, N. & McHeyzer-Williams, L. Molecular programming of B cell memory. Nat. Rev. Immunol. 12, 24–34 (2012).

104. Raychaudhuri, S. The Problem of Antigen Affinity Discrimination in B-Cell Immunology. ISRN Biomath. 2013, 1–18 (2013).

105. Chowdhury, S. et al. Programmable bacteria induce durable tumor regression and systemic antitumor immunity. Nat. Med. 25, 1057–1063 (2019).

106. Berg, P., Kornberg, R. D., Fancher, H. & Dieckmann, M. Competition between RNA polymerase and DNA polymerase for the DNA template. Biochem. Biophys. Res. Commun. 18, (1965).

107. Lim, D. & Maas, W. K. Reverse transcriptase in bacteria. Mol. Microbiol. 3, (1989).

108. Toro, N., Martínez-Abarca, F. & González-Delgado, A. The Reverse Transcriptases Associated with CRISPR-Cas Systems. Sci. Rep. 7, (2017).

109. Toro, N. & Nisa-Martínez, R. Comprehensive Phylogenetic Analysis of Bacterial Reverse Transcriptases. PLoS One 9, e114083 (2014).

110. Lampson, B. C., Inouye, M. & Inouye, S. Retrons, msDNA, and the bacterial genome. Cytogenet. Genome Res. 110, (2005).

111. Simon, D. M. & Zimmerly, S. A diversity of uncharacterized reverse transcriptases in bacteria. Nucleic Acids Res. 36, (2008).

112. Costa, T. R. D. et al. Secretion systems in Gram-negative bacteria: structural and mechanistic insights. Nat. Rev. Microbiol. 13, 343–359 (2015).

113. Erskine, E., MacPhee, C. E. & Stanley-Wall, N. R. Functional Amyloid and Other Protein Fibers in the Biofilm Matrix. J. Mol. Biol. 430, 3642–3656 (2018).

114. Tetz, G. V., Artemenko, N. K. & Tetz, V. V. Effect of DNase and Antibiotics on Biofilm Characteristics. Antimicrob. Agents Chemother. 53, (2009).

115. Guerrier-Takada, C., Gardiner, K., Marsh, T., Pace, N. & Altman, S. The RNA moiety of ribonuclease P is the catalytic subunit of the enzyme. Cell 35, (1983).

116. Huang, N. et al. Natural display of nuclear-encoded RNA on the cell surface and its impact on cell interaction. Genome Biol. 21, (2020).

117. Doyle, R. J., Koch, A. L. & Carstens, P. H. Cell wall-DNA association in Bacillus subtilis. J. Bacteriol. 153, (1983).

118. Hall, M. R., Meinke, W., Goldstein, D. A. & Lerner, R. A. Synthesis of Cytoplasmic Membrane-associated DNA in Lymphocyte Nucleus. Nat. New Biol. 234, 227–229 (1971).

119. Terekhov, S. S. et al. A kinase bioscavenger provides antibiotic resistance by extremely tight substrate binding. Sci. Adv. 6, (2020).

120. Rosenberg, M., Azevedo, N. F. & Ivask, A. Propidium iodide staining underestimates viability of adherent bacterial cells. Sci. Rep. 9, 6483 (2019).

121. Bacolla, A., Wang, G. & Vasquez, K. M. New Perspectives on DNA and RNA Triplexes As Effectors of Biological Activity. PLOS Genet. 11, e1005696 (2015).

122. Herbert, A. et al. Special Issue: A, B and Z: The Structure, Function and Genetics of Z-DNA and Z-RNA. Int. J. Mol. Sci. 22, 7686 (2021).

123. Tetz, G. & Tetz, V. Bacterial Extracellular DNA Promotes β-Amyloid Aggregation. Microorganisms 9, (2021).

124. Tetz, G. et al. Bacterial DNA promotes Tau aggregation. Sci. Rep. 10, (2020).

125. Tetz, V. & Tetz, G. Bacterial DNA induces the formation of heat-resistant disease-associated proteins in human plasma. Sci. Rep. 9, (2019).

126. Schindelin, J. et al. Fiji: an open-source platform for biological-image analysis. Nat. Methods 9, (2012).

127. Rueden, C. T. et al. ImageJ2: ImageJ for the next generation of scientific image data. BMC Bioinformatics 18, (2017).

128. Wang, X. et al. Hyaluronic acid modification of RNase A and its intracellular delivery using lipid-like nanoparticles. J. Control. Release 263, 39–45 (2017).

129. Hulsen, T., de Vlieg, J. & Alkema, W. BioVenn – a web application for the comparison and visualization of biological lists using area-proportional Venn diagrams. BMC Genomics 9, 488 (2008).

130. Manukumar, H. M. & Umesha, S. MALDI-TOF-MS based identification and molecular characterization of food associated methicillin-resistant Staphylococcus aureus. Sci. Rep. 7, (2017).

131. Bennett, R. W. & Monday, S. R. Saureus in International handbook of foodborne pathogens. in (ed. Miliotis MD, B. J.) 41–59 (2003).

132. Guide for the Care and Use of Laboratory Animals. (National Academies Press, 1996). doi:10.17226/5140.

133. Choudhry, P. High-Throughput Method for Automated Colony and Cell Counting by Digital Image Analysis Based on Edge Detection. PLoS One 11, (2016).

134. Jones ME, T. S. R. A. Luria-Delbrück fluctuation experiments: design and analysis. Genetics 136, 1209–1216 (1994).

135. Zhu L, Y. Z. Y. Q. T. Z. M. L. S. Z. L. X. Degradation of dexamethasone by acclimated strain of Pseudomonas Alcaligenes. Int. J. Clin. Exp. Med. 8, 10971 (2015).

