## Supplementary figure 1 for "Novel prokaryotic sensing and regulatory system employing previously unknown nucleic acids-based receptors"

Supplementary Figure S1. Absence of RNase A internalization in *B. pumilus*.

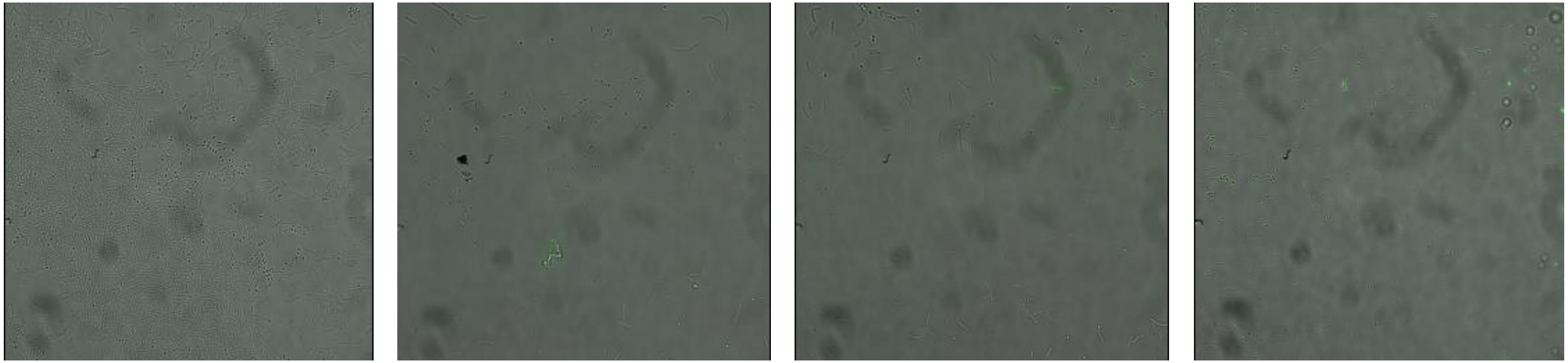

A B C D

*B. pumilus* cells were (A) untreated control (B) treated with fluorophore-labeled RNase A (100 µg/mL) for 15 min (C) treated with fluorophore-labeled RNase A (100 µg/mL) for 60 min (D) cultivated for 24h on agar supplemented with fluorophore-labeled RNase A (100 µg/mL)for 24h supplemented with labeled RNase A (100 µg/mL).
