## Supplementary figure 2 for "Novel prokaryotic sensing and regulatory system employing previously unknown nucleic acids-based receptors"

Supplementary Figure 2. Effect of TezRs removal on light sensing.

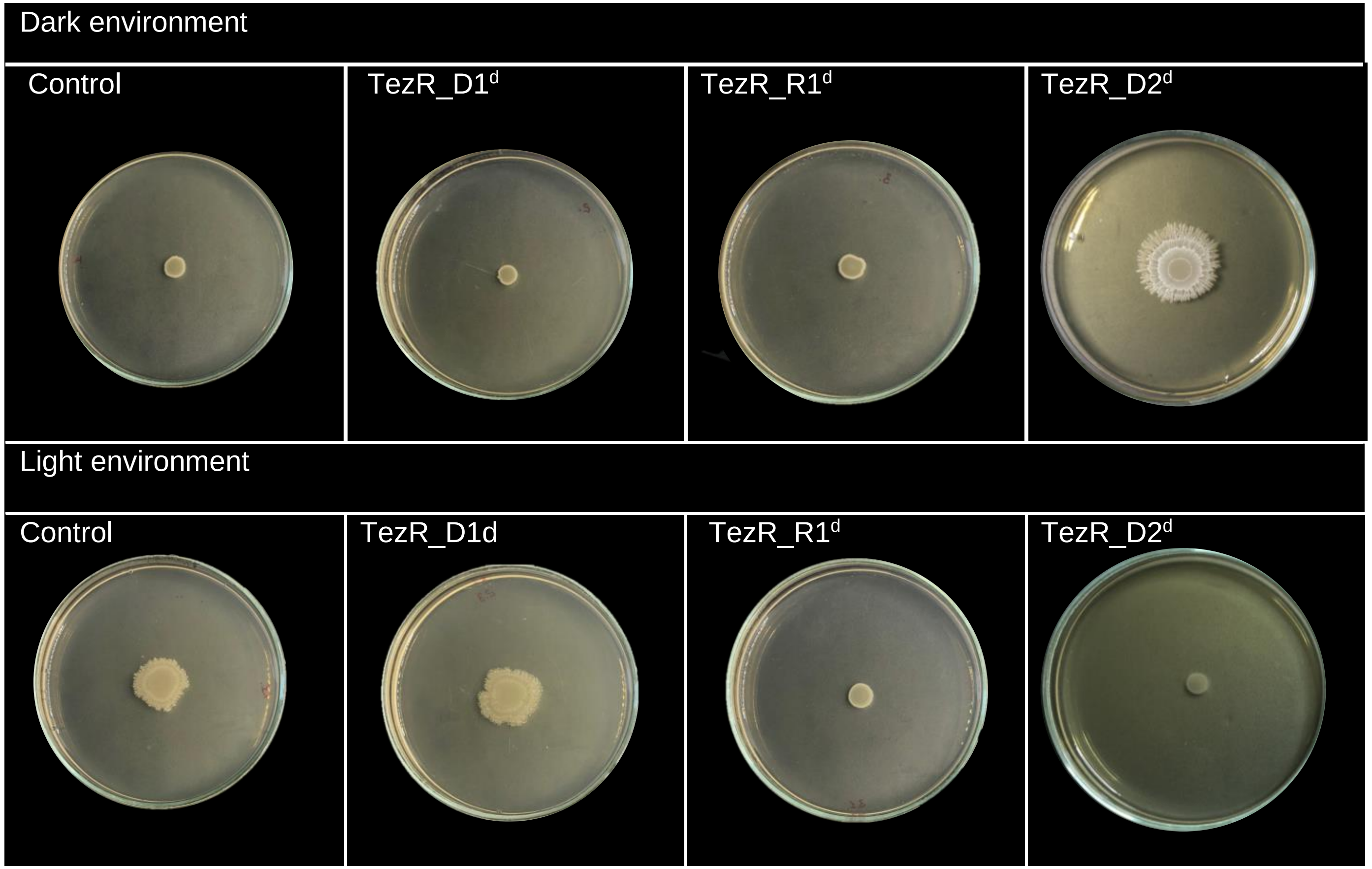
