## Supplementary table 1 for "Novel prokaryotic sensing and regulatory system employing previously unknown nucleic acids-based receptors"

Supplementary table 1. Effect of primary TezR_D1/R1 removal on bacterial size.

| Bacteria | Bacterial size (px2) | SD | p |
| --- | --- | --- | --- |
| Control *S. aureus* | 52.3 | 2.08 |  |
| *S. aureus* TezR_D1/R1^d^ | 59.1 | 6.94 | <0.001 |
| Control *E. coli* | 40.1 | 3.62 |  |
| *E. coli* TezR_D1/R1^d^ | 42.2 | 5.77 | 0.055 |
