## Supplementary table 2 for "Novel prokaryotic sensing and regulatory system employing previously unknown nucleic acids-based receptors"

Supplementary table 2. Effect of primary TezRs removal on the size of *B. pumilus* VT1200 biofilm.

| Bacteria | Biofilm size | SD | p |
| --- | --- | --- | --- |
| Control | 109290.7 | 17343.48 |  |
| TezR_D1^d^ | 158213.3 | 18154.98 | 0.028 |
| TezR_R1^d^ | 138990 | 6451.97 | 0.083 |
| TezR_D1/R1^d^ | 103975.67 | 11843.53 | 0.687 |
| TezR_D2^d^ | 1076281 | 64089.26 | <0.001 |
| TezR_R2^d^ | 137660 | 17984.35 | 0.121 |
| TezR_D2/R2^d^ | 288876.3 | 19735.85 | <0.001 |
| TezR_D1/D2^d^ | 256159.6 | 32023.40 | 0.006 |
| TezR_D1/R1/D2/R2^d^ | 134314 | 15963.79 | 0.14 |
