## Supplementary table 3 for "Novel prokaryotic sensing and regulatory system employing previously unknown nucleic acids-based receptors"

Supplementary table 3. Effect of TezR removal on sporulation under normal conditions.

| Bacteria | Sporulation (%) | SD | p |
| --- | --- | --- | --- |
| Control | 17.67 | 2.62 |  |
| TezR_D1^d^ | 76.33 | 5.312 | <0.001 |
| TezR_R1^d^ | 82 | 6.16 | <0.001 |
| TezR_D1/R1^d^ | 21.67 | 2.05 | 0.11 |
| TezR_D2^d^ | 0 | 0 | 0.007 |
| TezR_R2^d^ | 96 | 4.32 | <0.001 |
| TezR_D1/R1/D2/R2^d^ | 13.67 | 1.89 | 0.105 |
