## Supplementary table 4 for "Novel prokaryotic sensing and regulatory system employing previously unknown nucleic acids-based receptors"

Supplementary table 4. Effect of TezR removal on sporulation under stress conditions.

| Bacteria | Sporulation (%) | SD | p |
| --- | --- | --- | --- |
| Control normal conditions | 17.67 | 2.62 |  |
| Control stress conditions | 77.67 | 6.18 |  |
| TezR_D1^d^ | 21.67 | 3.40 | <0.001 |
| TezR_R1^d^ | 95.67 | 3.30 | 0.02 |
| TezR_D1/R1^d^ | 92.33 | 3.40 | 0.035 |
| TezR_D2^d^ | 4 | 2.16 | <0.001 |
| TezR_R2^d^ | 97.67 | 2.05 | 0.022 |
| TezR_D1/R1/D2/R2^d^ | 96.33 | 2.36 | 0.023 |
