## Supplementary table 5 for "Novel prokaryotic sensing and regulatory system employing previously unknown nucleic acids-based receptors"

Supplementary table 5. MICs of tested reverse transcriptase inhibitors and integrase inhibitor against control *S. aureus*.

| **Drug** | **Concentration of drug in µg/mL** | | | | | | | | | | | | |
| --- | --- | --- | --- | --- | --- | --- | --- | --- | --- | --- | --- | --- | --- |
|  | **512** | **256** | **128** | **64** | **32** | **16** | **8** | **4** | **2** | **1** | **0.5** | **0.25** | **0.125** |
| Etravirine | +* | + | + | + | + | + | + | + | + | + | + | + | + |
| Nevirapine | + | + | + | + | + | + | + | + | + | + | + | + | + |
| Raltegravir | + | + | + | + | + | + | + | + | + | + | + | + | + |

* “+” was used to mark the presence of bacterial growth
